# DeepDOX1: A Dual-Drive Framework Integrating Deep Learning and First-Principles Physics for Drug-Protein Affinity Prediction

**DOI:** 10.64898/2025.12.12.693818

**Authors:** Zheng Liu, Hao Sun, Yuliang Wang, Yanliang Ren, Li Rao, Zeyue Huang, Hongxuan Cao, Xiuqi Hu, Xinyue Zhu, Meng Li, Jian Wan

## Abstract

In this work, we present DeepDOX1, a dual-drive drug-protein affinity (DPA) prediction tool features the tight integration of a concise AI architecture and a quantum mechanics based representation. The first-principle physics generated features incorporating the interactions between the drug and the protein pocket allows a relatively simple CNN model trained on a relatively small training set to exhibit exceptional generalization capabilities across extensive testing. Notably, DeepDOX1 outperforms popular AI models in the tests simulating real-world drug design scenario and a highly challenging test set featuring covalent ligands, halogenated ligands and metalloproteins. In addition, we designed a series of novel covalent inhibitors targeting the diabetes target *hu*-FBPase using DeepDOX1. Subsequent experimental validation including enzyme-level bioactivity assays and crystal structure determination confirmed the DeepDOX1’s effectiveness in real-world drug design applications. It is conceivable that the combination of AI and first-principle QM might be one of the next breakthrough points of DPA prediction.

## Main

The evaluation of drug-protein binding affinity is a crucial step in drug design for screening and refining drug candidates. Computational methods have long been expected to be a cost-effective alternative to traditional experimental approaches for measuring drug-protein binding affinities, which typically demand significant time and resources. From relatively straightforward models such as Quantitative Structure - Activity Relationship (QSAR)^1^ and docking^2^, to the more physically advanced and intricate MM/PBSA^3^, free energy perturbation (FEP) techniques^4^, a diverse array of methods has been investigated for this purpose, yet none has emerged as the perfect solution. Recent years has seen a surge of artificial intelligence (AI) based drug protein affinity (DPA) prediction tools. In 2017, Ballester et al. introduced Δ_vina_RF20^5^, a Random Forest-based method for scoring the strength of DPA. Its subsequent version, Δ_vina_XGB^6^, was improved upon by implementing eXtreme Gradient Boosting (XGBoost). In 2016, Costa et al. developed a kernel learning-based DPA scoring function, KronRLS-MKL^7^, while Ragoza et al. pioneered the first convolutional neural network (CNN)-based DPA scoring function, demonstrating the feasibility of CNN architectures for this task^8^. Li et al. proposed DeepDTAF, a new CNN framework for binding affinity prediction in 2021^9^. Wegner et al. introduced DeepDock in the same year, leveraging graph convolutional networks (GCNs) to predict small-molecule conformations and partial binding affinities^10^. Several models were proposed in 2022, including PIGNet^11^, a physics-informed model for binding affinity prediction; PLA-MoRe^12^, a Transformer-based approach integrating structural and bioactive properties for enhanced binding affinity prediction; TankBind^13^, a GNN model combined with multilayer perceptrons (MLPs). In 2023, Hou et al. developed GenScore^14^ which is a Transformer-GCN hybrid, while Qiu et al. proposed MBP^15^, a multi-task learning-based scoring function. Concurrently, Wang et al. introduced the GNN-based model PLANET^16^, and Batista et al. reported HAC-Net^17^, an attention-enhanced CNN model. By 2025, AI-based DPA models will experience a further surge, including AdptDilatedGCN^18^, DTIAM^19^, SableBind^20^, and Boltz-2^21^. In this study, we were able to deploy and validated some of these popular models, utilizing their performance metrics as benchmarks for the validation of our method. The models validated in this work are summarized in **Table 1**.

**Table 1.** The three essential elements of some representative machine-learning based DTA prediction methods validated in this work, in comparison with DeepDOX1 developed in this work.

| Method | AI Architecture | Representation | Training Set Size |
| --- | --- | --- | --- |
| $\Delta_{vina}XGB^6$ | XGBoost | Drug: Vina Drug dependent terms;<br>Protein: bSASA features;<br>Interaction: Vina Interaction terms | 14,406 binders from PDBbind |
| TankBind<sup>13</sup> | Trigonometry-Aware Neural Networks | Drug: Graph generated by TorchDrug;<br>Protein: proximity 3D graph | 17,787 binders from PDBbind |
| DeepDock<sup>10</sup> | GCN | Drug: 2D graph based on atom and bond feature;<br>Protein: a polygon mesh of surface with electrostatics, hydrophathy, hydrogen bond, shape index encoded. | 15,000 complexes from PDBbind |
| SableBind<sup>20</sup> | Transformer | Drug: Pre-trained representation based on atom types, atom-atom distances and spatial positions;<br>Protein: sequence and structural information. | 19M molecules for drug representation pre-training, ~19000 binders from PDBbind for binding affinity training |
| DeepDOX1 | QM+CNN | Drug: atomic feature, RDKit predicted drug feature;<br>Protein: pocket residue feature;<br>Interaction: QM predicted binding energy and residue-drug interaction. | 9938 binders from PDBbind |

The effectiveness of an AI-based model hinges on three crucial factors: architecture, representation, and training data. As mentioned above, recently developed DPA prediction tools predominantly employ then state-of-the-art AI architectures. The representations used in DPA predictions mainly characterize information about proteins and drugs. Some early examples are protein sequence, drug chemical structure (often encoded as SMILES), drug physicochemical properties, and drug fingerprint information. The emerging Graph Neural Network (GNN) technology has spurred the application of graph-based representations for proteins and drugs^10^ ^13^. Recently, several studies reported the using of pre-trained information matrix generated by neural network for representation, exemplified by SableBind^20^ and Boltz-2^21^. The escalating complexity of both representations and architectures has driven an exponential demand for training dataset. The most widely used DPA training set is the PDBBind database, which contains both experimental crystal binding structure information and binding affinity information in terms of *K*_d_, *K*_i_ or IC ^22,23^. Its sample size increased from 2013 year’s 1,600 to 27,385 of its most recent version, barely enough to feed those data-hungry AI models. In addition, SableBind used 19 million unique molecules from the Uni-Mol project for the drug representation pre-training ^20^. Boltz-2 embodies the zenith of current training dataset scale in DPA prediction by using1.2 million binding affinity data gathered from PubChem, ChEMBL^24^ and BindingDB for the training of its binding affinity module. To sum up, the competition of literatures in this area has evolved to an arm race of advanced architectures, information-dense representations and massive training data. Consequently, AI-powered DPA prediction tools likely become more complex and data-driven in the future.

In contrast to AI largely driven by big data for its operation, quantum mechanics(QM) theories represent the extreme of physics based theoretical tool. QM provides physics description that most aligns with the electronic structure originated nature of protein-drug interactions including hydrogen bond^25^, van der Waals force (vdW)^26^, covalent bond^27^, halogen bond^28^ and coordination bond^29^, etc.

Although limited by their demanding computational cost, the application of QM methods in DPA prediction has a long history and was considered as “the logical next step in the evolution of this field” ^30^. See Ryde and Merz’s reviews for summaries of QM approaches involved in DPA prediction^30–33^. Previously, we have reported a series of work concerning protein-drug binding structure prediction and binding affinity prediction employing first principle QM^34–36^. In these works, density functional theories(DFT) were combined with divide and conquer strategy to provide reliable and affordable description of intermolecular interactions. Long years of experience taught us that with the computational cost problem largely been solved by the rapid development of computer technology and hybrid chemistry, the real challenge lies in the dynamics of solvents and protein. We found that sufficient sampling and explicit treatment of water molecules in protein cavity was essential for accurate assessment of solvation effect in DPA calculation, which led to a relatively expensive DPA prediction method DOX_BDW with a computational cost up to ∼500 core hours per calculation^34^. Simulation of protein-drug binding dynamics requires massive sampling of protein motion, too expensive for QM level DPA calculations. Therefore, the dynamics effect simulation requiring massive sampling is the shortest plank in the barrel for QM based drug design(QMDD), as shown in **Figure 1a**.

**Figure 1.**
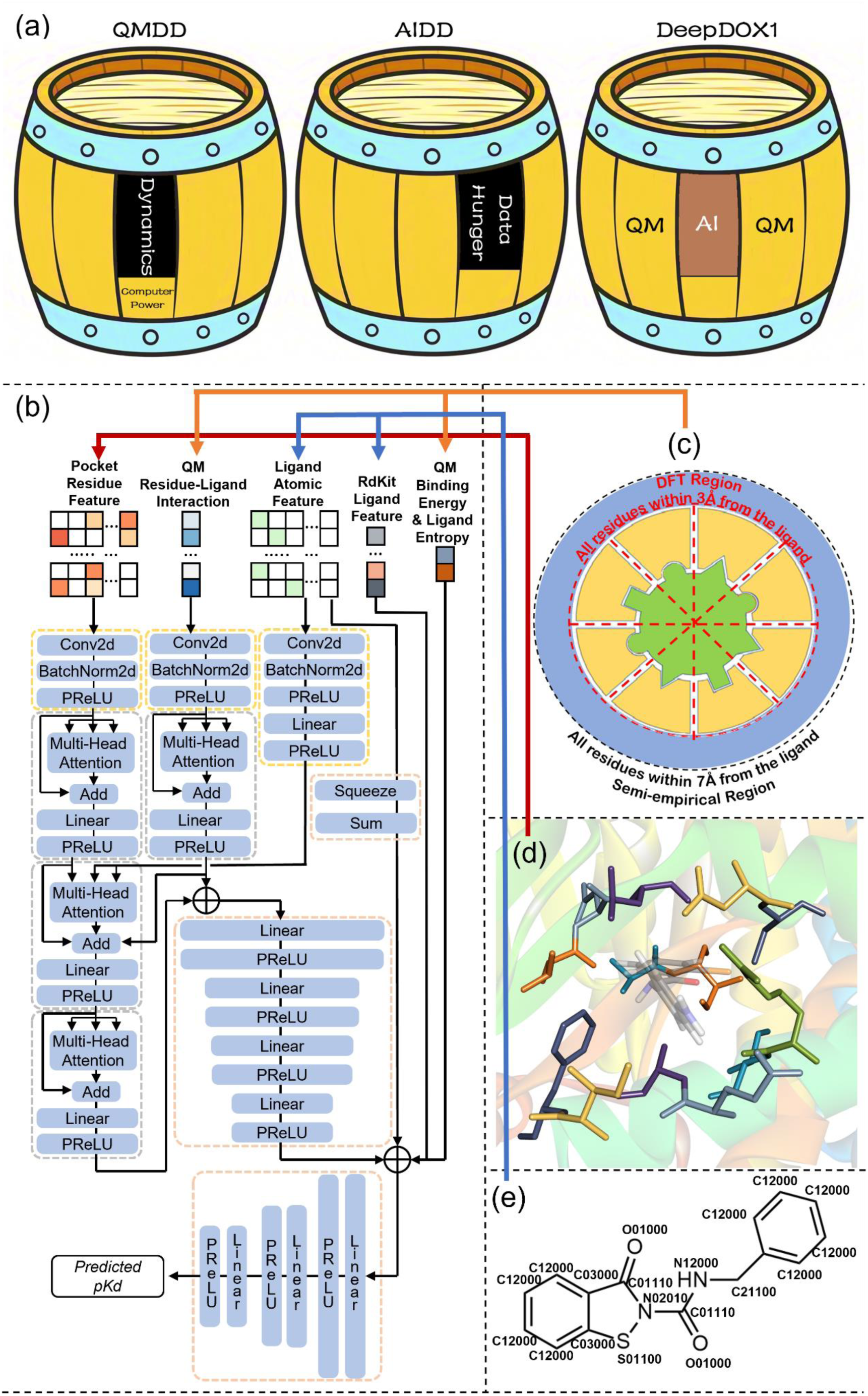
A schematic illustration of DeepDOX1. (a) Conceptual framework of combining QM and AI; (b) Architecture of DeepDOX1; (c) Illustration of QM binding energy computation; (d) Pocket residue denotes the protein residues within 3.5Å from any atom of the drug molecule; (e) Ligand representation consists of RDKit predicted molecular features and a novel atomic type defined by element type and chemical bonding environment, see **Supplementary Information Section 8** for a detailed illustration.

In this work, we developed a dual-drive framework DPA prediction tool, named as DeepDOX1. DeepDOX1 features the tight integration of a concise AI architecture and an interpretable, first principal QM based representation. Similar idea has been proven to be successful in the prediction of heats of formation^37,38^, NMR spectrum^39^ and molecular property prediction ^40^, but never been used in DPA prediction. This innovative approach leverages a succinct representation derived from QM-calculated protein pocket residue-ligand interaction energies, the pocket residue type and structural property and the ligand atomic type defined by its chemical environment, with the latter two closely related to the dynamics of protein ligand binding. From another perspective, we are using AI to predict the dynamics correction to QM simulated binding affinity based on stationary structure. Trained with a relatively small dataset (9938 protein-ligand binders), DeepDOX1 exhibit exceptional generalization capabilities in real-world drug design scenarios, according to a comprehensive validation against 1281 protein-ligand binders and application in *hu-*FBPase covalent inhibitor design. The details are discussed in the following sections.

## Results

### Overview of DeepDOX1 performance

An overview of DeepDOX1 performance is shown in Figure 2, in comparison with the performance of other AI models summarized in **Table 1**. Figure 2a shows the Spearman correlation and Pearson correlation obtained in the infamous CASF 2016 coreset test. Figure 2b shows the weighted average of Pearson correlation obtained on Merck-FEP test set, consists of 264 protein-ligand binders attributed to 8 protein-based subsets. Figure 2c shows the weighted average of Pearson correlation obtained in tests against a total of 608 protein-drug binding complexes belonging to 13 hit to lead optimization simulation subsets. Figure 2d shows the averaged Spearman correlation obtained in tests against 124 ligands featuring covalent ligands, halogenated ligands and metalloprotein ligands, indicating the performance on some challenging systems in DPA prediction. The performance of DeepDOX1 either surpasses all four AI representatives validated in this work or at least be comparable with the front runner in every tests. Considering that DeepDOX1 is a relatively simple architecture trained with smaller training dataset comparing with these AI model representatives, such performance was adequately good. Also, we would like to underscore that DeepDOX1’s performance is stable across all test sets, while the AI models’ performance exhibit significant volatility across different tests. This implies that the main forte of a QM/AI hybrid model lies in its extrapolability and universality.

**Figure 2.**
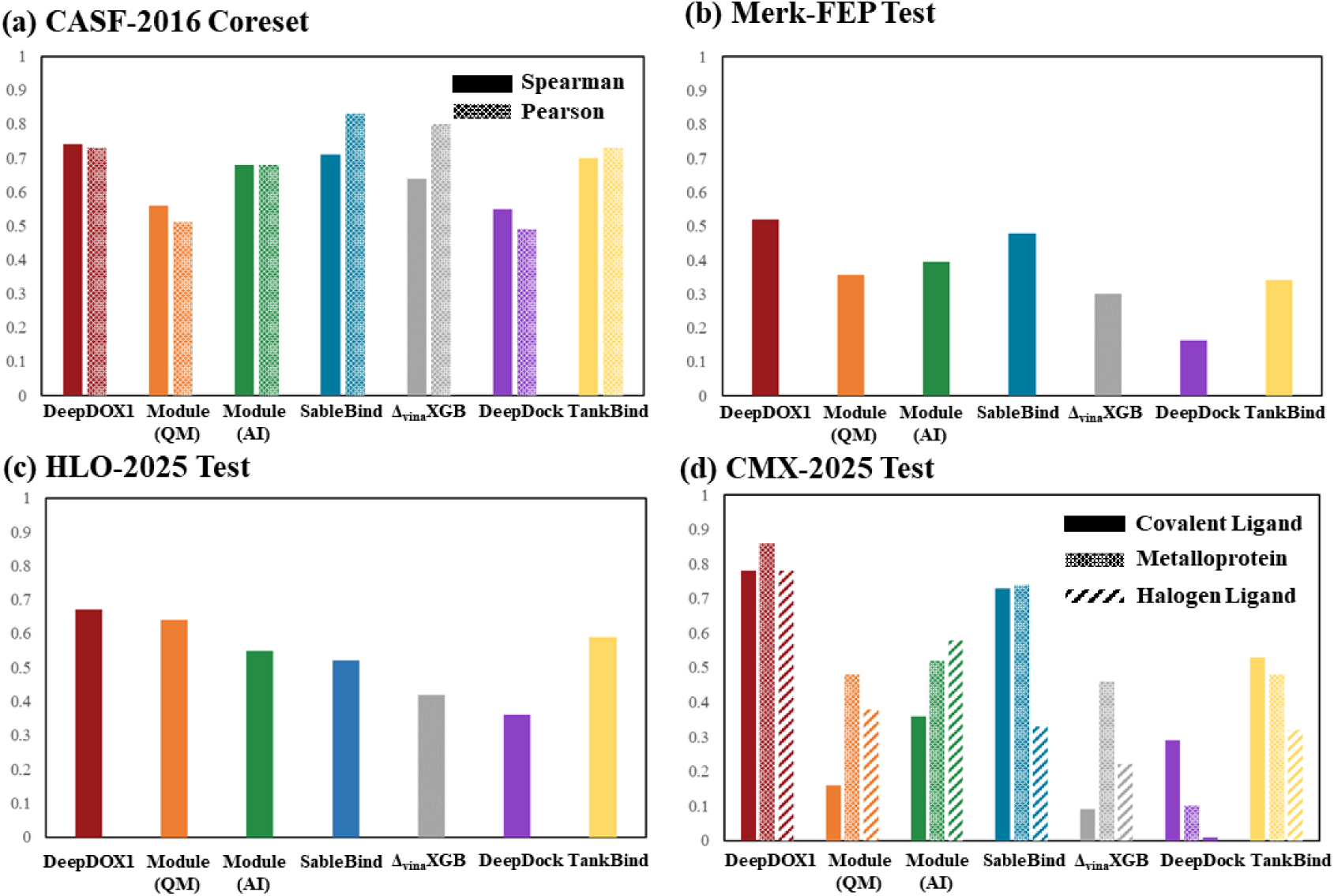
Overview of DeepDOX1 performance in comparison with AI powered DPA prediction representatives. (a) CASF 2016 coreset ranking and scoring test; ·(b) Weighted average of Pearson correlation obtained Merck-FEP Test set; (c) Weighted average of Pearson correlation obtained in HLO-2025 Tests; (d) Averaged Spearman correlation obtained in CMX-2025 Ranking Ability Test.

We also tested the performance of DeepDOX1 QM module and AI module separately for ablation study. Generally, the AI Module alone reaches an averaged performance of popular pure AI representatives studied in this work. The QM module, which means using the QM predicted binding energy alone, performs better in HLO test but poorly in other tests. However, their combination DeepDOX1 always outperform both of them, showcases our hybrid scheme’s capability of achieving synergy. See the following sections for more detailed discussion of each test. The raw data regarding CASF-2016, Merk-FEP, HLO-2025, and CMX-2025 could be found in **Supplementary Information Section 1-5.**

### CASF 2016 Coreset Ranking and Scoring Test

As the mostly cited validation system for DPA prediction, the famous CASF 2016 coreset test offers a standard to compare the ability of various DPA prediction methods. Figure 2a and **Extended Data Table 1** shows the obtained Spearman correlation for ranking power evaluation (ranking the five ligands of each target) and Pearson correlation for scoring power evaluation (plotting all 285 binding affinities together) of DeepDOX1, its QM module and AI module. In the original literature of CASF-2016^41^, Wang *et al.* suggested that the ranking power is applied only to the ligands of the same target protein. Therefore, the Spearman correlation should be calculated based on the 5 binders of the same target, then the 57 correlations were averaged to produce the overall Spearman correlation. We also tested SableBind, Δ_vina_XGB, DeepDock and TankBind using the code shared by the corresponding groups and the results are shown in **Extended Data Table 1** too. The obtained results are basically the same as the authors reported in their original literatures.

As shown in Figure 2a, DeepDOX1 outperform all pure AI powered methods in the ranking power evaluation with an averaged Spearman correlation of 0.74. It is closely followed by SableBind (0.71) and TankBind (0.70). Some other AI methods also reported similar target averaged CASF-2016 Spearman correlation, including GenScore (0.69) and IGModel (0.72). As for the scoring ability indicated by Pearson correlation of all 285 binders, DeepDOX1 falls behind SableBind, Δ_vina_XGB and TankBind, with a not so large difference. DeepDOX1 only include pocket residue information in the representation, likely affected the comparison between different proteins. The performance of isolated DeepDOX1 AI Module is comparable with other AI methods but significantly falls behind the integrated DeepDOX1.

Although CASF-2016 coreset are recommended for the baseline assessment since it is the most popular test in DPA prediction, its limited data size might not be sufficient to fully reflect the generalizability of the DPA method under evaluation. Also, all performance obtained on this dataset artificially benefited from the known exact binding structure, which is not available in typical drug design scenarios that requires the assistance of DPA prediction. Therefore, further tests are performed to fully assess the practical performance of DeepDOX1 in real-world drug design application, see below.

### Hit to Lead Optimization Scenario Tests

In recent years, literature has started to focus on the performance of DPA prediction methods in hit to lead optimization, which aims at transforming an initial promising compound (hit) into a more effective candidate (lead). Aiming at providing a unified framework for assessing the accuracy, efficiency, and robustness of DPA prediction method in hit to lead optimization, the Merck KGaA developed Merck-FEP Benchmark Dataset which contains 8 subsets and 264 protein-ligand binders divided by their targets^42^. The performance of many recently published AI approaches on Merck-FEP were validated in Gong’s work^20^, allowing us to evaluate the position of DeepDOX1 method among a range of AI-driven approaches. **Extended Data Table 2** shows the DeepDOX1 obtained Pearson correlation for all Merck-FEP subsets, together with its standalone AI module and QM module, and the AI methods validated in this work. DeepDOX1 achieved an averaged Pearson correlation of 0.52, surpassing all other approaches validated in this work, and all of the approaches discussed in Gong’s work^20^. However, none of the methods discussed achieved an averaged correlation larger than 0.6. Merck-FEP test were limited to comparison of similar ligands with no more than 10 atoms of modification since it is originally designed for evaluation of FEP methods. Therefore, the activity difference between the compounds is often minor, as evidenced by the fact that the subset averaged standard deviation of Merck-FEP set experimental data is merely 0.81 *pK*i units. This poses a significant challenge to AI based methods, which have a much lower computational load compared to FEP.

The HLO-2025 test set developed in this work was based on hit to lead studies that reported compounds with a larger experimental data variation (3∼4 *pK*_i_ units). However, that does not necessarily mean HLO-2025 is not challenging. Unlike CASF 2016 tests and Merck-FEP test whose protein-drug binding structures were known conditions, the unknown binding structure in the HLO-2025 tests significantly introduces higher challenge and reasonably reflects the demands of real-world drug design scenarios for DPA methods. Also, HLO-2025 dataset includes more drug targets, ligands, and diverse binding mechanism (covalent and non-covalent), spanning a wide range of chemical space. The Pearson correlations obtained in each HLO-2025 test sub-set were shown in Figure 3a and **Extended Data Table 3**, together with a weighted average by sample size of each sub-set. The averaged Pearson correlation of 0.68 indicated that DeepDOX1 significantly outperform all other models, attributed to its stability shown in the heatmap (Figure 3a). The pure AI models all have weaknesses which dragged down their overall performance, like Tyk2 and FKBP51 set for SableBind, JAK2, BRD4 and FKBP51 set for TankBind. When used alone, the AI Module of DeepDOX1 was able to obtain a similar overall performance as the pure AI representatives, while the QM Module ranked above all AI models but exhibited very poor performance on the Thrombin set, indicated by a correlation of −0.21. Interestingly, the AI component of DeepDOX1, which only obtained a 0.32 correlation on Thrombin set when used alone, was able to fix the severe mistake of the QM component and enabled the integrated DeepDOX1 to achieve a correlation of 0.72 on Thrombin set, indicating a remarkable synergy.

**Figure 3.**
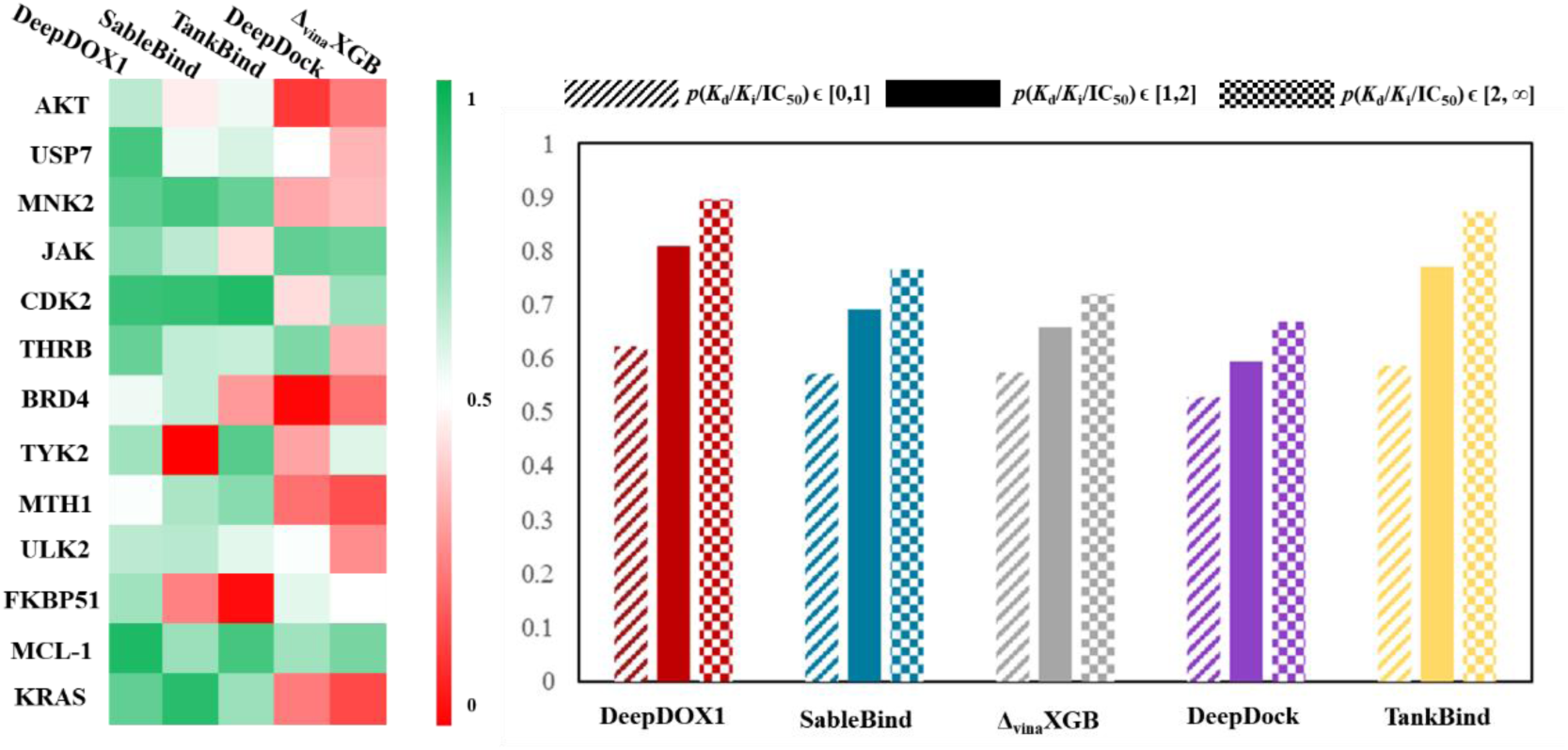
Performance of Different Methods on the HLO-2025 Test Set. (a) Heatmap of Pearson correlations obtained in HLO-2025 tests; (b) Comparison of prediction success rates, a successful prediction is defined as the compound predicted to be better is also the experimentally determined better compound when comparing two compounds bound to the same protein.

For the elucidation of the practical predictive power in drug screening from a different perspective, we also calculated prediction success rate for all compounds in the HLO-2025 test, while a successful prediction was defined as the drug predicted to be with stronger binding affinity is also the experimentally determined better drug when comparing two drugs of the same protein. As shown in Figure 3b, the success rates were calculated according to the difference of experimentally determined *pK*_d_, *pK*_i_, or *p*IC_50_. When the experimental data gap was less than one *pK*_d_/*K*_i_/IC_50_ unit, all models could only achieve a success rate a little better than throwing a coin. As the gap increased to 1∼2 *pK*_d_/*K*_i_/IC_50_ unit, DeepDOX1 achieved a success rate of 82%, outperforming all other models. Such performance indicated that DeepDOX1 fully met the requirements of hit to lead virtual optimization in real-world drug design scenario.

### CMX-2025: Performance of DeepDOX1 on some most challenging systems

Covalent bond^27^, halogen bond^43^ and coordination bond^44^ involved protein-drug binders were commonly seen in recent drug design studies, but they were quit uncommon in the PDBbind database which was usually used as training set. Among the 19443 complexes of PDBbind 2020 database, only 363 belongs to covalent drug, some 1122^45^ complexes contain halogen atoms in their drugs. Furthermore, electronic structure plays a crucial role in such interactions, leading to potential energy surface functions much more complicated than hydrogen bonds and vdW interactions. The aforementioned factors imply that covalent ligands, halogenated compounds and metalloproteins are likely to pose a significant challenge for DPA prediction methods. In this work, we have constructed a test set, dubbed as CMX-2025 augmented test, consists of 31 subsets related to covalent ligand, metalloprotein ligand and halogenated ligand for the validation of DPA methods on challenging protein-drug interactions.

As shown in Figure 4 and **Extended Data Table 4**, the CMX-2025 augmented test revealed that DeepDOX1 was the sole method evaluated in this study to demonstrate strong ranking ability (averaged Spearman correlation R_S_ > 0.7) across all types of challenging interactions. The Δ_vina_XGB, DeepDock and TankBind obtained averaged R_S_ was no larger than 0.6 for any type of interactions, indicating the challenges in characterizing these types of interactions. SableBind achieved similar performance as DeepDOX1 in covalent sub-test and metalloprotein sub-test. It is worthy of mention that 66% of the data included in the CMX-2025 metalloprotein sub-test was also found in the training data of SableBind, which likely strengthened its performance. SableBind also exhibited notable weaknesses, as indicated by the R_S_=0.33 obtained on halogenated ligands. In fact, all AI methods evaluated in this work exhibited poor performance on halogenated ligands, likely owing to the unique electrostatic and directional characteristics of halogen bond^46^. Incorporating information related to charge spatial distribution into molecular representations might help enhance the performance of AI based methods on halogenated drugs. Note that without such information in the representation, DeepDOX1 still reached a R_S_ of 0.78, demonstrates the value of this method for halogenated drugs.

**Figure 4.**
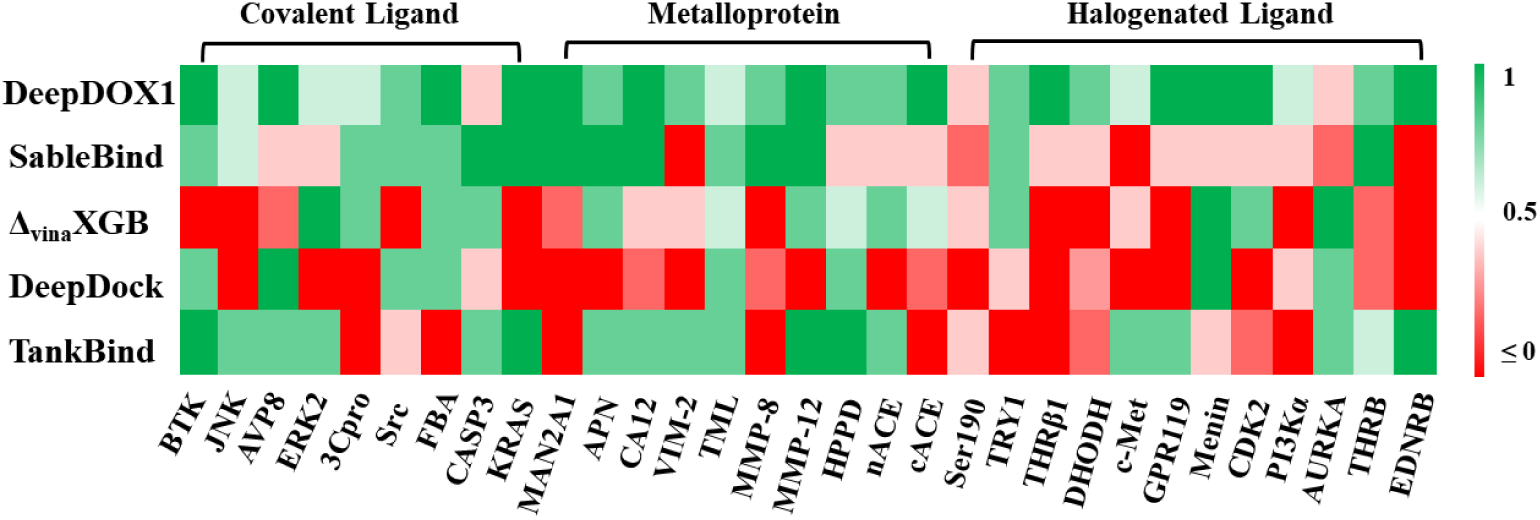
Heatmap of Spearman correlations obtained in CMX-2025 tests

### Application: *hu*-FBPase targeted covalent ligand design

Only real-world drug design practice could reveal the true value of a theoretical drug design approach. We have utilized DeepDOX1 whose training set does not contain any covalent ligand data to optimize a covalent inhibitor hit compound (compound 11) of *hu-*FBPase obtained in our previous work^47^. As shown in Figure 5, the compound 11 bind to *hu*-FBPase with a covalent bond between its bromoacetamide warhead and CYS179 residue, and a measured inhibition constant (*K*_i_) of 7.13 *μ*M. We employed Cov_DOX and DeepDOX1 for virtual screening of a batch of derivatives of compound 11, see **Supplementary Information Section 6** for a detailed list. Virtual designed derivatives with predicted bind affinity larger than compound 11(DeepDOX1 *pK*_i_ = 7.65) are shown in Figure 5a, considered as potential improvement over compound 11. For the moment, we have accomplished the synthesis and experimental activity assay for two of them, compound 11n (reported in our previous work) and **11t** (first reported in this work). The compound **11t** exhibited 5-fold decline of *K*_i_, which was considerably successful hit compound modification. We are working on the synthesis of the other compounds with predicted high binding affinity and we will address them in our future work.

**Figure 5.**
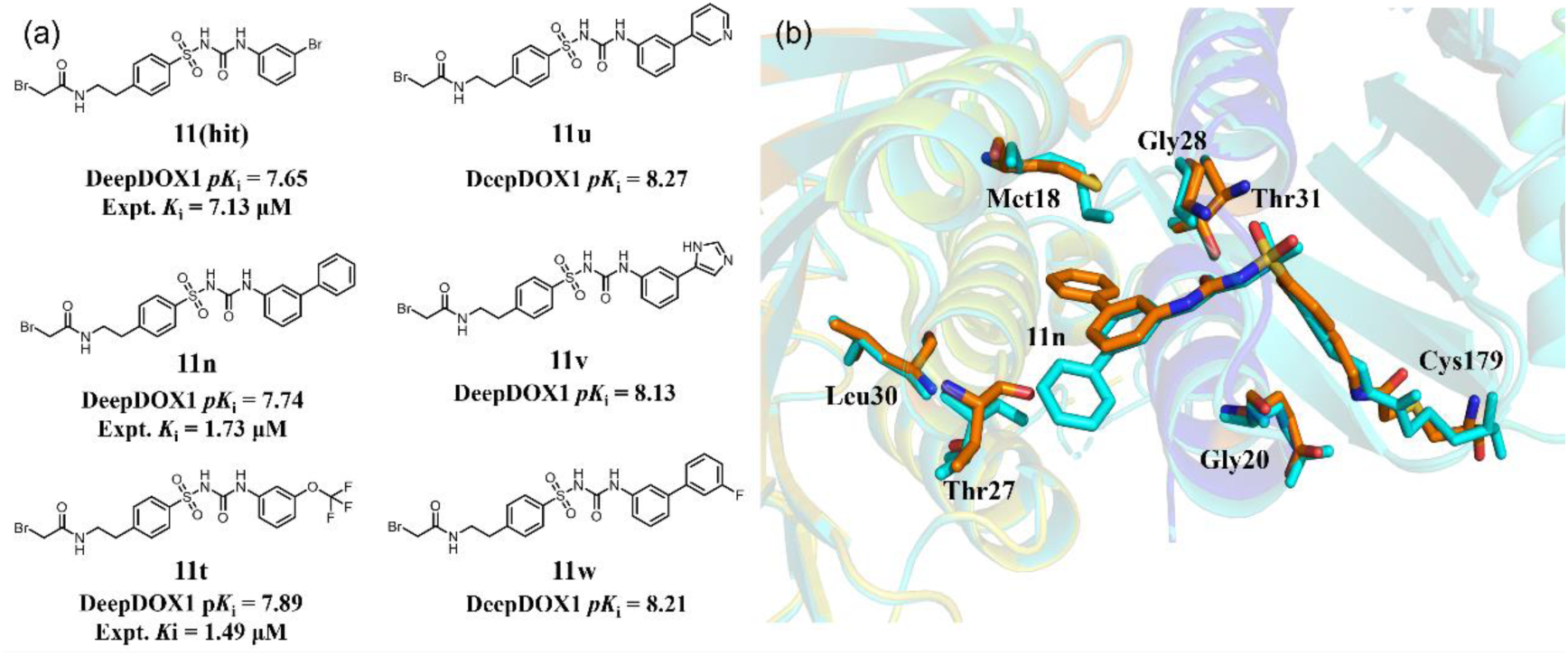
*hu*-FBPase targeted covalent ligand designed using DeepDOX1. (a) Hit compound 11 and its derivatives with DeepDOX1 predicted *pK*_i_ higher than itself, experimental data is also presented if available; (b) Crystal structure (orange) of 11n complexed with *hu*-FBPase obtained in this work, in comparison with previously predicted binding structure (cyan) of 11n employing a combination of Cov_DOX and DeepDOX1.

It is worthy of mention that the scoring of 11n and 11t was based on predicted binding structure employing a DeepDOX1 assisted Cov_DOX calculation (**see Supplementary Information Section 7**). In this work, we have obtained the crystal structure of compound 11n complexed with *hu*-FBPase (PDBID: 9XVI, attached in the SI material). As shown in Figure 5b, the superposition of crystal structure and Cov_DOX/DeepDOX1 predicted binding structure for compound 11n found that they were quite similar. The experimental and theoretical binding structure only differ with each other on the orientation of biphenyl group, likely owing to the rotation of Met18 residue side group. We suggest that both binding poses exist in solution and contribute to activity. We also found that the predicted binding affinity exhibits significant fluctuations(∼1*pK*_i_ unit) depending on the distinct binding poses, highlights the importance of binding structure and necessity to discuss both AI generated binding structure and AI employed binding structure.

## Discussion

In this work, we conducted first-principle quantum mechanics calculation for ∼10000 protein-ligand complexes and obtained their interaction energetics for representation. A streamlined CNN architecture was employed to dig the relationship between these first principal interaction energetics and experimental binding affinity, which resulted in the first AI and QM dual-drive framework DPA prediction method DeepDOX1. In essence, we are using AI to predict the dynamics correction to QM simulated binding affinity based on stationary structure. Therefore, a concise CNN architecture and a relatively small training set was enough to learn from this physically constrained representation. The DeepDOX1 approach demonstrates solid performance across various tests, including CASF-2016 test, Merck-FEP test, HLO-2025 test and CMX-2025 test, revealing its broad generality and good extrapolation performance. To evaluate its true value in real-world drug design applications, we designed novel covalent inhibitors targeting the *hu*-FBPase Cys179 site using DeepDOX1, even though its training set did not include any covalent ligand. Subsequent experimental validation revealed strengthened activity of newly design compounds. These comprehensive results demonstrate that by incorporating first-principles physics, we can build AI based DPA prediction models that are not only effective but also remarkably robust and generalizable across diverse interaction types and target classes, without relying on complex architectures or massive training datasets. The DeepDOX1 could be accessed by multiple ways. To allow convenient tryout of DeepDOX1, a web server is provided, which can be assessed at http://doxwebserver.ccnu.edu.cn/.

The DeepDOX1 main program, the data sets, the corresponding DOX/Cov_DOX program and a user’s guide could be found in the supplementary materials. A detailed illustration of DeepDOX1 architecture could be found in supplementary materials too. A typical DeepDOX1 calculation takes roughly 1 hour on a single CPU (AMD EPYC 7V12) server with 64 cores, which is commonly available in laboratories. If such computational cost is still considered too much, the DOX predicted QM binding energies and residue-ligand interaction features could be replaced with the results of cheaper QM theories like semi-empirical methods, re-train the model and get a customized version of DeepDOX1. We are working on a new version of DeepDOX1 with much less computational cost which will be released in near future.

## Methods

Figure 1 demonstrated the conceptual framework (Fig 1a), architecture (Fig 1b) and representation (Fig 1c-e) of DeepDOX1. Compared to mainstream AI based DPA prediction tools, DeepDOX1 is characterized by the integration of first-principle QM theory and divide & conquer computational strategy, which enables a compact neural network architecture, a straightforward representation and moderately high yet affordable computational cost. In the following contents, we will present a detailed introduction to the model architecture, representation, and data set.

### Representation

As indicated in **Table 1**, researchers have already exploited the power of the most advanced deep learning frameworks and reached the celling of protein-drug experimental data for training set. Therefore, the representation is likely the next breakthrough points of AI based DPA prediction. As shown in Figure 1b, six types of representation are fed into the model, including protein pocket residue feature, QM predicted residue-ligand interaction, ligand atomic feature, RDKit predicted ligand feature, QM predicted binding energy and ligand entropy.

The protein pocket residue feature is fed into a fixed-size 100×1×64 input tensor, where each one of the 100 channels correspond to a residue within 3.5Å from any atom of the drug molecule and unused channels tensor are padded with zeros. The characterization of each residue is performed independently. For each used channel, column 1-20 of the 1×64 matrix is the one-hot encoding of amino acid type, column 21-28 is the one-hot encoding of metal ion cofactor, column 29-37 is the one-hot encoding of secondary structure type predicted by DSSP^48^, column 38 is the charge of the residue. The rest columns are preserved for future work. Currently, they are padded with zeros.

For the ligand atomic feature, all ligand non-hydrogen atoms are divided into 64 atomic types based on the element type and the type and number of atoms bonded with it, see **supplemental materials Table S8** for a detailed description. Each row of the ligand atomic feature matrix shown in Figure 1b is the one-hot encoding of a ligand atom atomic type. The matrix is column-summed to produce a 1×64 feature vector representing the atomic type distribution.

Meanwhile, 12 selected molecular descriptors, including the topological polar surface area, the Wiener index, and the number of rotatable bonds (see **supplemental materials Table S9** for a complete list), are computed using RDKit to generate a 1×12 feature vector representing the physicochemical characteristics of the ligand.

The QM binding energy feature is predicted using a protocol similar to the DOX_P method proposed in our previous work, which is a combination of first principle density functional theory ωB97xD^49^, semiempirical PM7 and hybrid chemistry method eXtended ONIOM^50^. The ligand entropy term and residue-ligand interaction, which is the estimated interaction energy between each residue and the ligand, are also obtained in the QM binding energy calculation. The residue-ligand interaction energies are fed into a 100×1 vector. See **supplemental materials section 10** for a detailed illustration of QM binding energy calculation.

### Model Architecture

As illustrated in Figure 1b, the input features were categorized into four groups: protein pocket information, quantum mechanical (QM) properties, small-molecule one-hot encodings, and RDKit-based molecular descriptors. For the first three categories—pocket residues, QM terms, and one-hot encodings—each row in the feature matrix corresponds to a specific physical entity, such as an individual amino acid within the binding cavity. To process these structured representations, we applied a 2D convolutional neural network to each row, compressing the row-wise information into a unified 1 × 100 feature vector.

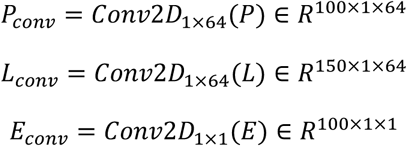

To hierarchically integrate information within the protein pocket (residue–residue), within the ligand (atom–atom), and between the protein and ligand, we employed a combination of self-attention and cross-attention mechanisms. This design endows the feature representation with biophysically meaningful interactions^51^.

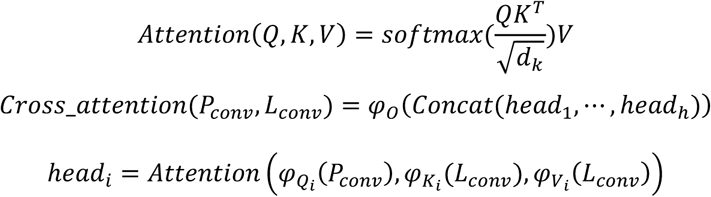

We leveraged a multi-head attention mechanism to fuse the QM-computed residue-wise energy decompositions with the geometric features of the binding pocket, generating a physically meaningful representation.

With the protein-ligand interactions comprehensively captured by the attention mechanism, the model’s remaining focus shifted to the intrinsic properties of the small molecule. These properties were processed through a simple fully connected layer to produce the final binding affinity prediction.

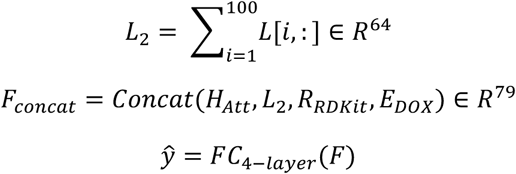

### Datasets

Similar to the approaches demonstrated in **Table 1**, all the training set data utilized in this study is sourced from the PDBbind database. The PDBbind v.2020 version contains 19,443 protein-drug binding complex structure and corresponding experimental *K*_d_, *K*_i_ or IC_50_ data. The training set of DeepDOX1 only included 9,938 complexes from the PDBbind v.2022 dataset, after screening based on Lipinski’s Rule of Five, RDKit processability and DOX processability.

To comprehensively and systematically evaluate the reliability and practicality of DeepDOX1 in multiple drug design scenarios, we employed a test set consisting of up to 1281 protein-ligand complexes, including the infamous CASF-2016 core set consists of 285 complexes, the public Merck-FEP Benchmark Dataset which contains 8 subsets and 264 protein-ligand binders divided by their targets, the HLO-2025 test set consists of 13 targets (12 non-covalent and 1 covalent) and 608 ligands for the mimic of hit to lead optimization scenario, and the CMX-2025 test set consists of 124 complexes related with challenging metalloprotein, covalent ligands and halogenated ligands. The HLO-2025 test set and CMX-2025 test set was the extension of the test sets employed in our previous work^34–36^. The HLO-2025 test set data was carefully collected from medicinal literatures which reported more than 20 compounds with similar backbone and diverse activity (≥1000-fold activity difference between worst and best compound) in the hit or lead optimization step. The CMX-2025 test set was organized focusing on three interaction types, including covalent bond, coordination bond (metalloprotein) and halogen bond. The covalent ligand set in CMX-2025 consists of 9 target based subsets and 36 ligands (4 ligands for each target, the same below). The metalloprotein set consists of 10 target based subsets and 40 ligands forming coordination bond with the metal cofactor of metal-dependent enzymes. The halogen set consists of 12 target based subsets and each subset contain four structurally conserved ligands that only differ by one halogen substitution.

There is no overlap between the training set and any of the test sets. All complexes were processed with an updated Cov_DOX software developed in our previous work to generate the representation features demonstrated above. For the training set and CASF-2016 test set complexes, the initial structures for calculations were based on the corresponding crystal structures without further binding pose sampling. For HLO-2025 tests and CMX-2025 tests, the binding structures were predicted using a simplified DOX protocol implemented in Cov_DOX, closely mimicking real-world drug design scenarios. For the Merck-FEP test set, the structures downloaded from the authors’ Github webpage (https://github.com/ORCAaAaA-ui/-Merk-FEP-benchmark-set) were used as initial structure for the calculation without further binding pose sampling. Note that for some cases in Merk-FEP test set, we found clash between the authors provided ligand structure with the corresponding protein structure. Simplified DOX protocol predicted binding pose were used for these individual cases.

### Deploy of Benchmarked Methods

All benchmarked models (e.g., TankBind (https://github.com/luwei0917/TankBind/), Δ_vina_XGB (https://github.com/jenniening/deltaVinaXGB), DeepDock (https://github.com/OptiMaL-PSE-Lab/DeepDock)), SableBind(https://github.com/MIALAB-RUC/SableBind) were evaluated in environments consistent with their official implementations. Specifically, we used the same hardware platform employed for training our own method comprising an NVIDIA RTX 4090 (48 GB) GPU and an AMD EPYC 9754 processor and strictly followed the authors’ provided code repositories for environment setup and testing. Note that the implementation of SableBind did not function on the NVIDIA RTX 4090 GPU during our setup. Pending resolution of this compatibility issue, all reported data for SableBind are based on a 1× NVIDIA V100 32GB GPU/Intel Xeon Gold 6130 CPU configuration to ensure a fair comparison.

### *hu-*FBPase covalent inhibitor design

Covalent protein inhibitor features the covalent bond between the ligand and the protein, offering significant advantages in improving potency, prolonging pharmacodynamic effects, and creating novel selectivity profiles. Human fructose-1,6-bisphosphatase (*hu*-FBPase) is the rate limiting enzyme in the gluconeogenesis pathway, considered as a potential target associated with cancer and type II diabetes. In our previous work^47^, we designed a series of non-covalent interaction/covalent-bond dual-driven bromoacetamide inhibitors targeting the AMP pocket and neighboring cysteine residue (CYS179) of *hu*-FBPase. In this work, we continued to optimize the bromoacetamide lead compounds based on DeepDOX1 predicted binding affinity. Note that DeepDOX1 naturally supports covalent ligand with the help of implemented QM theory. The experimental dissociation constant (*Kᵢ*) for selected high-scoring compounds was determined using the enzymatic activity assay established in our previous work.^47^ Also, we were able to obtain a new crystal structure for one of the high scored compounds, which is close to the binding structure predicted using DeepDOX1 assisted Cov_DOX protocol^36^. The details of experimental methods, including synthesis, enzymatical activity assay and protein X-ray crystallography, could be found in **Supplemental Materials Section 7**.

## Data availability

The protein-ligand complex structures and binding affinity data used in the training set and CASF-2016 test were obtained from the PDBbind database v.2020^22^, which is publicly accessible at http://www.pdbbind.org.cn. The structures and binding affinity data used in Merk-FEP test were obtained from Daniel Kuhn’s^42^ public repository (https://github.com/ORCAaAaA-ui/-Merk-FEP-benchmark-set). The binding affinity data and ligand chemical structure used in HLO-2025 test and CMX-2025 test were collected from various literatures. The corresponding protein-ligand complex structures were predicted using DOX, which are accessible at our public repository (https://github.com/DeepDOX/DeepDOX1, https://doi.org/10.5281/zenodo.17787765) and supplementary material of this article.

## Code availability

The source code of DeepDOX1, the training data saved in Pytorch format, and trained model weights for DeepDOX1 are freely available at our public repository (https://github.com/DeepDOX/DeepDOX1, https://doi.org/10.5281/zenodo.17787765). The Cov_DOX/DOX program could be found on our public repository too. We also provide webserver for Cov_DOX/DOX and DeepDOX1 calculation, which could be accessed at http://doxwebserver.ccnu.edu.cn/.

## Extended Data

**Extended Data Table 1.** The Spearman correlation and Pearson correlation obtained in the CASF 2016 Coreset test.

| Model | DeepDOX1 | Module (QM) | Module (AI) | SableBind | $\Delta_{vina}XGB$ | Deep-Dock | Tank-Bind |
| --- | --- | --- | --- | --- | --- | --- | --- |
| Spearman<sup>a</sup> | 0.74 | 0.56 | 0.68 | 0.71<sup>c</sup> | 0.65 | 0.55 | 0.70 |
| Pearson<sup>b</sup> | 0.72 | 0.51 | 0.68 | 0.83 | 0.80 | 0.49 | 0.73 |

- a. According to the original literature of CASF-2016<sup>41</sup>, the Spearman correlation was calculated based on the 5 binders of the same target in each subset, then the 57 correlations were averaged for the evaluation of ranking power. Note that SableBind<sup>20</sup> reported Spearman correlation based on the ranking of all 285 binders, in contradiction to the intent of CASF-2016 literature. In such case, we re-calculated the target averaged Spearman correlation based on the literature's original data. - b. The Pearson correlation was calculated according to the regression of all 285 binders. - c. Spearman correlation recomputed with the accepted methodology.

**Extended Data Table 2.** Performance of DeepDOX1 (Pearson correlation) on Merck-FEP Test in comparison with pure AI based DPA methods. The results of DeepDOX1 were obtained in this work.

| Protein (data size) | DeepDOX1 | Module (QM) | Module (AI) | SableBind<sup>a</sup> | $\Delta_{vina}XGB$ | Deep-Dock | Tank-Bind |
| --- | --- | --- | --- | --- | --- | --- | --- |
| HIF-2α(42) | 0.45 | 0.36 | 0.29 | 0.43 | 0.27 | -0.17 | 0.32 |
| PFKFB3(40) | 0.55 | 0.36 | 0.44 | 0.49 | 0.51 | 0.29 | 0.22 |
| Eg5(28) | 0.74 | 0.01 | 0.40 | 0.71 | -0.03 | 0.29 | 0.14 |
| CDK8(33) | 0.39 | 0.52 | 0.45 | 0.56 | 0.61 | 0.34 | 0.44 |
| SHP-2(26) | 0.39 | 0.74 | 0.52 | 0.44 | 0.71 | 0.24 | 0.47 |
| SYK(44) | 0.53 | 0.18 | 0.33 | 0.19 | -0.21 | 0.06 | 0.29 |
| c-MET(24) | 0.70 | 0.31 | 0.53 | 0.63 | 0.61 | 0.05 | 0.59 |
| TNKS2(27) | 0.48 | 0.49 | 0.40 | 0.39 | 0.18 | 0.35 | 0.41 |
| Average<sup>b</sup> | 0.52 | 0.36 | 0.40 | 0.46 | 0.30 | 0.16 | 0.34 |

- a. The results of SableBind were taken from its literature<sup>20</sup>. - b. Weighted average of Pearson correlation by sample size of each sub-set.

**Extended Data Table 3.** Detailed Results of HLO-2025 Test^a^.

| Protein (data size) | DeepDOX1 | Module (QM) | Module (AI) | SableBind | $\Delta_{vina}XGB$ | Deep-Dock | Tank-Bind |
| --- | --- | --- | --- | --- | --- | --- | --- |
| AKT1 (75) | 0.57 | 0.69 | 0.60 | 0.42 | 0.22 | 0.10 | 0.48 |
| USP7(27) | 0.78 | 0.68 | 0.72 | 0.48 | 0.32 | 0.45 | 0.52 |
| MNK2(29) | 0.74 | 0.44 | 0.66 | 0.78 | 0.33 | 0.30 | 0.72 |
| JAK2(50) | 0.66 | 0.83 | 0.42 | 0.57 | 0.71 | 0.73 | 0.39 |
| CDK2(72) | 0.80 | 0.86 | 0.72 | 0.81 | 0.63 | 0.39 | 0.84 |
| THRB(24) | 0.72 | -0.21 | 0.51 | 0.56 | 0.31 | 0.68 | 0.55 |
| BRD4(59) | 0.48 | 0.32 | 0.49 | 0.56 | 0.20 | 0.01 | 0.27 |
| Tyk2(75) | 0.62 | 0.69 | 0.66 | -0.09 | 0.51 | 0.29 | 0.75 |
| MTH1(32) | 0.46 | 0.78 | 0.43 | 0.60 | 0.14 | 0.20 | 0.66 |
| ULK2(26) | 0.57 | 0.54 | 0.52 | 0.58 | 0.25 | 0.46 | 0.50 |
| FKBP51(20) | 0.62 | 0.48 | -0.77 | 0.23 | 0.45 | 0.50 | 0.02 |
| MCL-1(81) | 0.85 | 0.74 | 0.70 | 0.63 | 0.69 | 0.62 | 0.78 |
| KRAS(38) | 0.73 | 0.70 | 0.48 | 0.83 | 0.13 | 0.22 | 0.63 |
| Average<sup>a</sup> | 0.67 | 0.64 | 0.55 | 0.52 | 0.42 | 0.36 | 0.59 |

a. Pearson correlation obtained for each sub-set and its weighted average by sample size of each sub-set.

**Extended Data Table 4.** Detailed Results of CMX-2025 Challenging Test^a^.

| Protein | DeepDOX1 | Module (QM) | Module (AI) | SableBind | $\Delta_{vina}XGB$ | Deep-Dock | Tank-Bind |
| --- | --- | --- | --- | --- | --- | --- | --- |
| Covalent Ligand |  |  |  |  |  |  |  |
| BTK | 1.00 | 1.00 | 1.00 | 0.80 | -0.60 | 0.80 | 1.00 |
| JNK | 0.60 | -0.80 | -1.00 | 0.60 | -0.80 | -0.60 | 0.80 |
| AVP8 | 1.00 | 0.40 | 1.00 | 0.40 | 0.20 | 1.00 | 0.80 |
| ERK2 | 0.60 | -0.60 | -0.40 | 0.40 | 1.00 | -0.40 | 0.80 |
| 3Cpro | 0.60 | 0.60 | 0.20 | 0.80 | 0.80 | -0.20 | 0.00 |
| Src | 0.80 | -0.80 | 0.80 | 0.80 | -1.00 | 0.80 | 0.40 |
| FBA | 1.00 | 0.40 | 1.00 | 0.80 | 0.80 | 0.80 | -0.80 |
| CASP3 | 0.40 | 0.40 | 0.20 | 1.00 | 0.80 | 0.40 | 0.80 |
| KRAS | 1.00 | 0.80 | 0.40 | 1.00 | -0.40 | 0.00 | 1.00 |
| Average | 0.78 | 0.16 | 0.36 | 0.73 | 0.09 | 0.29 | 0.53 |

| Metalloprotein |  |  |  |  |  |  |  |
| --- | --- | --- | --- | --- | --- | --- | --- |
| MAN2A1 | 1.00 | 1.00 | 0.80 | 1.00 | 0.20 | -0.40 | 0.00 |
| APN | 0.80 | 0.80 | 0.80 | 1.00 | 0.80 | 0.00 | 0.80 |
| CA12 | 1.00 | 0.00 | 1.00 | 1.00 | 0.40 | 0.20 | 0.80 |
| VIM-2 | 0.80 | 0.80 | 0.80 | 0.00 | 0.40 | -0.40 | 0.80 |
| TML | 0.60 | 0.60 | 0.60 | 0.80 | 0.60 | 0.80 | 0.80 |
| MMP-8 | 0.80 | 0.80 | 0.60 | 1.00 | -0.40 | 0.20 | 0.00 |
| MMP-12 | 1.00 | 1.00 | 0.80 | 1.00 | 0.80 | 0.00 | 1.00 |
| HPPD | 0.80 | 0.80 | 0.80 | 0.40 | 0.60 | 0.80 | 1.00 |
| nACE | 0.80 | 0.40 | -0.80 | 0.40 | 0.80 | -0.40 | 0.80 |
| cACE | 1.00 | 1.00 | -0.20 | 0.40 | 0.60 | 0.20 | -0.40 |
| Average | 0.86 | 0.48 | 0.52 | 0.74 | 0.46 | 0.10 | 0.48 |
| Halogenated Ligand |  |  |  |  |  |  |  |
| Ser190 | 0.40 | 0.20 | 0.20 | 0.20 | 0.40 | 0.00 | 0.40 |
| TRY1 | 0.80 | -0.80 | -0.40 | 0.80 | 0.80 | 0.40 | 0.00 |
| THRβ1 | 1.00 | -0.80 | 0.40 | 0.40 | -0.40 | -0.40 | -0.80 |
| DHODH | 0.80 | 0.80 | 0.80 | 0.40 | 0.00 | 0.30 | 0.20 |
| c-Met | 0.60 | 0.00 | 0.80 | -0.20 | 0.40 | -0.40 | 0.80 |
| GPR119 | 1.00 | -0.40 | 0.00 | 0.40 | 0.00 | -0.20 | 0.80 |
| Menin | 1.00 | 0.80 | 1.00 | 0.40 | 1.00 | 1.00 | 0.40 |
| CDK2 | 1.00 | 0.40 | 1.00 | 0.40 | 0.80 | -1.00 | 0.20 |
| PI3Kα | 0.60 | 0.80 | 0.60 | 0.40 | -0.80 | 0.40 | -0.60 |
| AURKA | 0.40 | 1.00 | 1.00 | 0.20 | 1.00 | 0.80 | 0.80 |
| THRB | 0.80 | -0.80 | 0.80 | 1.00 | 0.20 | 0.20 | 0.60 |
| EDNRB | 1.00 | -0.30 | 0.80 | -0.40 | -0.80 | -0.20 | 1.00 |
| Average | 0.78 | 0.38 | 0.58 | 0.33 | 0.22 | 0.01 | 0.32 |

- a. Spearman correlation obtained for each target by ranking the corresponding four ligands of the same target.

## Acknowledgements

This study was financed by the National Key Research and Development Program of China (2023YFD1700501), National Natural Science Foundation of China (22373039, 22177036, 22377030).

## Author contributions

L. R., Z. L. and H. S., conceived the DeepDOX1 architecture and representation. Z. L., Y. W., X. Z. and M. L. carried out the calculations. Y. R., Z. H. and H. C. designed and carried out the synthesis, biological assay and crystal experiments. L. R. and J. W. supervised the project. Z. L., H. S., Y. W. and Y. R. contributed equally to the article.

## Competing interests

All authors declare no competing interests.

## Notes

https://github.com/DeepDOX/DeepDOX1

### Competing Interest Statement

The authors have declared no competing interest.

## Reference

1. Tropsha, A., Isayev, O., Varnek, A., Schneider, G. & Cherkasov, A. Integrating QSAR modelling and deep learning in drug discovery: the emergence of deep QSAR. Nature Reviews Drug Discovery 23, 141–155 (2023).

2. Kitchen, D.B., Decornez, H., Furr, J.R. & Bajorath, J. Docking and scoring in virtual screening for drug discovery: methods and applications. Nature Reviews Drug Discovery 3, 935–949 (2004).

3. Genheden, S. & Ryde, U. The MM/PBSA and MM/GBSA methods to estimate ligand-binding affinities. Expert Opinion on Drug Discovery 10, 449–461 (2015).

4. Steinbrecher, T. & Labahn, A. Towards Accurate Free Energy Calculations in Ligand Protein-Binding Studies. Current Medicinal Chemistry 17, 767–785 (2010).

5. Wang, C. & Zhang, Y. Improving scoring-docking-screening powers of protein-ligand scoring functions using random forest. J Comput Chem 38, 169–177 (2017).

6. Lu, J., Hou, X., Wang, C. & Zhang, Y. Incorporating Explicit Water Molecules and Ligand Conformation Stability in Machine-Learning Scoring Functions. J Chem Inf Model 59, 4540–4549 (2019).

7. Nascimento, A.C., Prudencio, R.B. & Costa, I.G. A multiple kernel learning algorithm for drug-target interaction prediction. BMC Bioinformatics 17, 46 (2016).

8. Ragoza, M., Hochuli, J., Idrobo, E., Sunseri, J. & Koes, D.R. Protein-Ligand Scoring with Convolutional Neural Networks. J Chem Inf Model 57, 942–957 (2017).

9. Wang, K., Zhou, R., Li, Y. & Li, M. DeepDTAF: a deep learning method to predict protein-ligand binding affinity. Brief Bioinform 22(2021).

10. Liao, Z. et al. DeepDock: Enhancing Ligand-protein Interaction Prediction by a Combination of Ligand and Structure Information. in 2019 IEEE International Conference on Bioinformatics and Biomedicine (BIBM) 311–317 (2019).

11. Moon, S., Zhung, W., Yang, S., Lim, J. & Kim, W.Y. PIGNet: a physics-informed deep learning model toward generalized drug-target interaction predictions. Chem Sci 13, 3661–3673 (2022).

12. Li, Q. et al. PLA-MoRe: A Protein-Ligand Binding Affinity Prediction Model via Comprehensive Molecular Representations. J Chem Inf Model 62, 4380–4390 (2022).

13. Lu, W. et al. TANKBind: Trigonometry-Aware Neural NetworKs for Drug-Protein Binding Structure Prediction. in Advances in Neural Information Processing Systems Vol. 35 (eds Koyejo, S. et al.) 7236–7249-7236–7249 (Curran Associates, Inc.).

14. Shen, C. et al. A generalized protein-ligand scoring framework with balanced scoring, docking, ranking and screening powers. Chem Sci 14, 8129–8146 (2023).

15. Yan, J. et al. Multi-task bioassay pre-training for protein-ligand binding affinity prediction. Brief Bioinform 25(2023).

16. Zhang, X. et al. PLANET: A Multi-objective Graph Neural Network Model for Protein–Ligand Binding Affinity Prediction. Journal of Chemical Information and Modeling 64, 2205–2220 (2023).

17. Kyro, G.W., Brent, R.I. & Batista, V.S. HAC-Net: A Hybrid Attention-Based Convolutional Neural Network for Highly Accurate Protein-Ligand Binding Affinity Prediction. J Chem Inf Model 63, 1947–1960 (2023).

18. Wang, W. et al. AdptDilatedGCN: Protein-ligand binding affinity prediction based on multi-scale interaction fusion mechanism and dilated GCN. Int J Biol Macromol 311, 143751 (2025).

19. Lu, Z. et al. DTIAM: a unified framework for predicting drug-target interactions, binding affinities and drug mechanisms. Nat Commun 16, 2548 (2025).

20. Li, J. & Gong, X. Harnessing pre-trained models for accurate prediction of protein-ligand binding affinity. BMC Bioinformatics 26, 55 (2025).

21. Passaro, S. et al. Boltz-2: Towards Accurate and Efficient Binding Affinity Prediction. *bioRxiv* (2025).

22. Liu, Z. et al. Forging the Basis for Developing Protein-Ligand Interaction Scoring Functions. Acc Chem Res 50, 302–309 (2017).

23. Liu, Z. et al. PDB-wide collection of binding data: current status of the PDBbind database. Bioinformatics 31, 405–12 (2015).

24. Davies, M. et al. ChEMBL web services: streamlining access to drug discovery data and utilities. Nucleic Acids Res 43, W612–20 (2015).

25. Xu, Z., Zhang, Q., Shi, J. & Zhu, W. Underestimated Noncovalent Interactions in Protein Data Bank. J Chem Inf Model 59, 3389–3399 (2019).

26. Rummel, L. & Schreiner, P.R. Advances and Prospects in Understanding London Dispersion Interactions in Molecular Chemistry. Angew Chem Int Ed Engl 63, e202316364 (2024).

27. Tamura, T., Kawano, M. & Hamachi, I. Targeted Covalent Modification Strategies for Drugging the Undruggable Targets. Chem Rev 125, 1191–1253 (2025).

28. Li, J. et al. Impact of Halogen Bonds on Protein-Peptide Binding and Protein Structural Stability Revealed by Computational Approaches. J Med Chem 67, 4782–4792 (2024).

29. de Almeida, L.G.N. et al. Matrix Metalloproteinases: From Molecular Mechanisms to Physiology, Pathophysiology, and Pharmacology. Pharmacol Rev 74, 712–768 (2022).

30. Raha, K. et al. The role of quantum mechanics in structure-based drug design. Drug Discov Today 12, 725–31 (2007).

31. Manathunga, M., Gotz, A.W. & Merz, K.M., Jr. Computer-aided drug design, quantum-mechanical methods for biological problems. Curr Opin Struct Biol 75, 102417 (2022).

32. Pantano, S. et al. Computational Chemistry in the Global South: A Latin American Perspective. J Chem Inf Model 65, 1677–1678 (2025).

33. Jin, H. & Merz, K.M., Jr. Integrating Machine Learning and Quantum Circuits for Proton Affinity Predictions. J Chem Theory Comput 21, 2235–2243 (2025).

34. Liu, J. et al. DOX_BDW: Incorporating Solvation and Desolvation Effects of Cavity Water into Nonfitting Protein-Ligand Binding Affinity Prediction. J Chem Inf Model 63, 4850–4863 (2023).

35. Rao, L. et al. DOX: A new computational protocol for accurate prediction of the protein-ligand binding structures. J Comput Chem 37, 336–44 (2016).

36. Wei, L. et al. Cov_DOX: A Method for Structure Prediction of Covalent Protein-Ligand Bindings. J Med Chem 65, 5528–5538 (2022).

37. Wu, J. & Xu, X. The X1 method for accurate and efficient prediction of heats of formation. J Chem Phys 127, 214105 (2007).

38. Hu, L., Wang, X., Wong, L. & Chen, G. Combined first-principles calculation and neural-network correction approach for heat of formation. The Journal of Chemical Physics 119, 11501–11507 (2003).

39. Yan, W. & Xu, X. Accurate Prediction of Nuclear Magnetic Resonance Parameters via the XYG3 Type of Doubly Hybrid Density Functionals. J Chem Theory Comput 18, 2931–2946 (2022).

40. Wang, Z. et al. X2-GNN: A Physical Message Passing Neural Network with Natural Generalization Ability to Large and Complex Molecules. J Phys Chem Lett 15, 12501–12512 (2024).

41. Su, M. et al. Comparative Assessment of Scoring Functions: The CASF-2016 Update. Journal of Chemical Information and Modeling 59, 895–913 (2018).

42. Schindler, C.E.M. et al. Large-Scale Assessment of Binding Free Energy Calculations in Active Drug Discovery Projects. J Chem Inf Model 60, 5457–5474 (2020).

43. Li, J. et al. Impact of Halogen Bonds on Protein–Peptide Binding and Protein Structural Stability Revealed by Computational Approaches. Journal of Medicinal Chemistry 67, 4782–4792 (2024).

44. Yu, J.L. et al. MeDBA: the Metalloenzyme Data Bank and Analysis platform. Nucleic Acids Res 51, D593–D602 (2023).

45. Zhang, Q., Xu, Z. & Zhu, W. The Underestimated Halogen Bonds Forming with Protein Side Chains in Drug Discovery and Design. J Chem Inf Model 57, 22–26 (2017).

46. Zhu, Z. et al. Halogen bonding in differently charged complexes: basic profile, essential interaction terms and intrinsic sigma-hole. Phys Chem Chem Phys 21, 15106–15119 (2019).

47. Cao, H. et al. Structure-Guided Design of Affinity/Covalent-Bond Dual-Driven Inhibitors Targeting the AMP Site of FBPase. Journal of Medicinal Chemistry 67, 20421–20437 (2024).

48. Hekkelman, M.L., Salmoral, D.Á., Perrakis, A. & Joosten, R.P. DSSP 4: FAIR annotation of protein secondary structure. Protein Science 34(2025).

49. Chai, J.D. & Head-Gordon, M. Systematic optimization of long-range corrected hybrid density functionals. J Chem Phys 128, 084106 (2008).

50. Guo, W., Wu, A., Zhang, I.Y. & Xu, X. XO: an extended ONIOM method for accurate and efficient modeling of large systems. J Comput Chem 33, 2142–60 (2012).

51. Vaswani, A. et al. Attention is all you need. in Proceedings of the 31st International Conference on Neural Information Processing Systems 6000–6010 (Curran Associates Inc., Long Beach, California, USA, 2017).

