## Supplementary Information for "DeepDOX1: A Dual-Drive Framework Integrating Deep Learning and First-Principles Physics for Drug-Protein Affinity Prediction"

Section 1. Detailed results of the CASF-2016 Test^1^

**Table S1.** Predicted binding affinity against the CASF-2016 core set.

| Target | PDBID | DeepDOX1 | Expt. |
| --- | --- | --- | --- |
| 1 | 4llx | 4.47 | 2.89 |
| 1 | 5c28 | 4.97 | 5.66 |
| 1 | 3uuo | 6.35 | 7.96 |
| 1 | 3ui7 | 6.73 | 9.00 |
| 1 | 5c2h | 7.25 | 11.09 |
| 2 | 2v00 | 5.08 | 3.66 |
| 2 | 3wz8 | 6.50 | 5.82 |
| 2 | 3pww | 8.18 | 7.32 |
| 2 | 3prs | 8.90 | 7.82 |
| 2 | 3uri | 7.72 | 9.00 |
| 3 | 4m0z | 6.85 | 5.19 |
| 3 | 4m0y | 5.71 | 6.46 |
| 3 | 3qgy | 6.95 | 7.80 |
| 3 | 4qd6 | 7.05 | 8.64 |
| 3 | 4rfm | 8.15 | 10.05 |
| 4 | 4cr9 | 4.74 | 4.10 |
| 4 | 4cra | 6.73 | 7.22 |
| 4 | 4x6p | 7.96 | 8.30 |
| 4 | 4crc | 7.24 | 8.72 |
| 4 | 4ty7 | 7.55 | 9.52 |
| 5 | 5aba | 5.67 | 2.98 |
| 5 | 5a7b | 5.81 | 3.57 |
| 5 | 4agn | 5.67 | 3.97 |
| 5 | 4agp | 6.77 | 4.69 |
| 5 | 4agq | 6.11 | 5.01 |
| 6 | 3bgz | 6.49 | 6.26 |
| 6 | 3jya | 5.52 | 6.89 |
| 6 | 2c3i | 6.63 | 7.60 |
| 6 | 4k18 | 6.76 | 8.96 |
| 6 | 5dwr | 7.95 | 11.22 |
| 7 | 3mss | 5.48 | 4.66 |
| 7 | 3k5v | 6.65 | 6.30 |
| 7 | 3pyy | 6.28 | 6.86 |
| 7 | 2v7a | 6.88 | 8.30 |
| 7 | 4twp | 7.20 | 10.00 |
| 8 | 3wtj | 5.58 | 6.53 |
| 8 | 3zdg | 5.11 | 7.10 |
| 8 | 3u8k | 5.79 | 8.66 |
| 8 | 4qac | 6.44 | 9.40 |
| 8 | 3u8n | 6.23 | 10.17 |
| 9 | 1a30 | 4.60 | 4.30 |
| 9 | 2qnq | 8.46 | 6.11 |
| 9 | 1g2k | 8.09 | 7.96 |
| 9 | 1eby | 8.58 | 9.70 |
| 9 | 3o9i | 9.47 | 11.82 |
| 10 | 4lzs | 5.09 | 4.80 |
| 10 | 3u5j | 5.88 | 5.61 |
| 10 | 4wiv | 6.94 | 6.26 |
| 10 | 4ogj | 7.29 | 6.79 |
| 10 | 3p5o | 7.22 | 7.30 |
| 11 | 1ps3 | 6.07 | 2.28 |
| 11 | 3dx1 | 5.49 | 3.58 |
| 11 | 3d4z | 5.49 | 4.89 |
| 11 | 3dx2 | 5.94 | 6.82 |
| 11 | 3ejr | 6.88 | 8.57 |
| 12 | 3l7b | 5.36 | 2.40 |
| 12 | 4eky | 5.20 | 3.52 |
| 12 | 3g2n | 4.75 | 4.09 |
| 12 | 3syr | 5.38 | 5.10 |
| 12 | 3ebp | 5.97 | 5.91 |
| 13 | 2w66 | 5.11 | 4.05 |
| 13 | 2w4x | 5.01 | 4.85 |
| 13 | 2wca | 6.17 | 5.60 |
| 13 | 2xj7 | 5.66 | 6.66 |
| 13 | 2vvn | 5.50 | 7.30 |
| 14 | 3aru | 5.05 | 3.22 |
| 14 | 3arv | 5.83 | 5.64 |
| 14 | 3ary | 4.91 | 6.00 |
| 14 | 3arq | 6.56 | 6.40 |
| 14 | 3arp | 7.51 | 7.15 |
| 15 | 4ih5 | 5.52 | 4.11 |
| 15 | 4ih7 | 5.56 | 5.24 |
| 15 | 3cj4 | 6.65 | 6.51 |
| 15 | 4eo8 | 6.97 | 8.15 |
| 15 | 3gnw | 8.79 | 9.10 |
| 16 | 1gpk | 6.31 | 5.37 |
| 16 | 1gpn | 6.46 | 6.48 |
| 16 | 1h23 | 7.57 | 8.35 |
| 16 | 1h22 | 6.80 | 9.10 |
| 16 | 1e66 | 6.82 | 9.89 |
| 17 | 3f3a | 5.31 | 4.19 |
| 17 | 3f3c | 5.07 | 6.02 |
| 17 | 4mme | 6.38 | 6.50 |
| 17 | 3f3d | 5.55 | 7.16 |
| 17 | 3f3e | 6.12 | 7.70 |
| 18 | 2wbg | 5.50 | 4.45 |
| 18 | 2cbv | 5.53 | 5.48 |
| 18 | 2j78 | 5.80 | 6.42 |
| 18 | 2j7h | 5.74 | 7.19 |
| 18 | 2cet | 6.31 | 8.02 |
| 19 | 3udh | 5.26 | 2.85 |
| 19 | 3rsx | 5.18 | 4.41 |
| 19 | 4djv | 7.03 | 6.72 |
| 19 | 2vkm | 9.01 | 8.74 |
| 19 | 4gid | 8.20 | 10.77 |
| 20 | 4jfs | 5.73 | 5.27 |
| 20 | 4j28 | 5.53 | 5.70 |
| 20 | 2wvt | 5.85 | 6.12 |
| 20 | 2xii | 6.29 | 7.20 |
| 20 | 4pcs | 5.92 | 7.85 |
| 21 | 3rr4 | 5.26 | 4.55 |
| 21 | 1s38 | 5.07 | 5.15 |
| 21 | 1r5y | 5.12 | 6.46 |
| 21 | 3gc5 | 6.26 | 7.26 |
| 21 | 3ge7 | 6.7 | 8.70 |
| 22 | 4dli | 6.38 | 5.62 |
| 22 | 2zb1 | 6.49 | 6.32 |
| 22 | 4f9w | 6.86 | 6.94 |
| 22 | 3e92 | 6.55 | 8.00 |
| 22 | 3e93 | 7.50 | 8.85 |
| 23 | 4owm | 4.31 | 2.96 |
| 23 | 3twp | 4.12 | 3.92 |
| 23 | 3r88 | 4.62 | 4.82 |
| 23 | 4gkm | 5.65 | 5.17 |
| 23 | 3qqs | 5.25 | 5.82 |
| 24 | 3gv9 | 4.61 | 2.12 |
| 24 | 3gr2 | 4.73 | 2.52 |
| 24 | 4kz6 | 4.98 | 3.10 |
| 24 | 4jxs | 5.32 | 4.74 |
| 24 | 2r9w | 5.98 | 5.10 |
| 25 | 2hb1 | 5.45 | 3.80 |
| 25 | 1bzc | 6.44 | 4.92 |
| 25 | 2qbr | 7.07 | 6.33 |
| 25 | 2qbq | 7.52 | 7.44 |
| 25 | 2qbp | 7.35 | 8.40 |
| 26 | 1q8t | 5.64 | 4.76 |
| 26 | 1ydr | 6.41 | 5.52 |
| 26 | 1q8u | 7.03 | 5.96 |
| 26 | 1ydt | 6.29 | 7.32 |
| 26 | 3ag9 | 5.99 | 8.05 |
| 27 | 3fcq | 5.09 | 2.77 |
| 27 | 1z9g | 5.32 | 5.64 |
| 27 | 1qf1 | 6.55 | 7.32 |
| 27 | 5tmn | 8.18 | 8.04 |
| 27 | 4tmn | 6.34 | 10.17 |
| 28 | 4ddk | 4.32 | 2.29 |
| 28 | 4ddh | 5.03 | 3.32 |
| 28 | 3ivg | 6.47 | 4.30 |
| 28 | 3coz | 5.47 | 5.57 |
| 28 | 3coy | 6.08 | 6.02 |
| 29 | 3pxf | 5.40 | 4.43 |
| 29 | 4eor | 7.49 | 6.30 |
| 29 | 2xnb | 6.76 | 6.83 |
| 29 | 1pxn | 6.43 | 7.15 |
| 29 | 2fvd | 6.99 | 8.52 |
| 30 | 4k77 | 5.88 | 6.63 |
| 30 | 4e5w | 7.36 | 7.66 |
| 30 | 4ivb | 7.88 | 8.72 |
| 30 | 4ivd | 7.12 | 9.52 |
| 30 | 4ivc | 7.41 | 10.00 |
| 31 | 4f09 | 6.36 | 6.70 |
| 31 | 4gfm | 5.72 | 7.22 |
| 31 | 4hge | 6.84 | 7.92 |
| 31 | 4e6q | 6.54 | 8.36 |
| 31 | 4jia | 6.26 | 9.22 |
| 32 | 2brb | 5.98 | 4.86 |
| 32 | 2br1 | 6.58 | 5.14 |
| 32 | 3jvr | 6.28 | 5.72 |
| 32 | 3jvs | 6.90 | 6.54 |
| 32 | 1nvq | 7.68 | 8.25 |
| 33 | 3acw | 6.09 | 4.76 |
| 33 | 4ea2 | 7.36 | 6.44 |
| 33 | 2zcr | 7.60 | 6.87 |
| 33 | 2zy1 | 6.54 | 7.40 |
| 33 | 2zcq | 7.72 | 8.82 |
| 34 | 1bcu | 5.35 | 3.28 |
| 34 | 3bv9 | 6.32 | 5.36 |
| 34 | 1oyt | 6.63 | 7.24 |
| 34 | 2zda | 6.13 | 8.40 |
| 34 | 3utu | 8.46 | 10.92 |
| 35 | 3u9q | 5.02 | 4.38 |
| 35 | 2yfe | 5.95 | 6.63 |
| 35 | 3fur | 8.07 | 8.00 |
| 35 | 3b1m | 7.50 | 8.48 |
| 35 | 2p4y | 8.17 | 9.00 |
| 36 | 3uo4 | 6.96 | 6.52 |
| 36 | 3up2 | 6.53 | 7.40 |
| 36 | 3e5a | 7.03 | 8.23 |
| 36 | 2wtv | 7.81 | 8.74 |
| 36 | 3myg | 7.43 | 10.70 |
| 37 | 3kgp | 4.32 | 2.57 |
| 37 | 1c5z | 4.48 | 4.01 |
| 37 | 1o5b | 6.17 | 5.77 |
| 37 | 1owh | 6.12 | 7.40 |
| 37 | 1sqa | 6.92 | 9.21 |
| 38 | 4jsz | 4.89 | 2.30 |
| 38 | 3kwa | 4.64 | 4.08 |
| 38 | 2weg | 5.22 | 6.50 |
| 38 | 3ryj | 5.67 | 7.80 |
| 38 | 3dd0 | 6.20 | 9.00 |
| 39 | 2xdl | 5.03 | 3.10 |
| 39 | 3b27 | 5.76 | 5.16 |
| 39 | 1yc1 | 6.49 | 6.17 |
| 39 | 3rlr | 6.90 | 7.52 |
| 39 | 2yki | 8.36 | 9.46 |
| 40 | 1z95 | 7.62 | 7.12 |
| 40 | 3b68 | 7.09 | 8.40 |
| 40 | 3b5r | 7.54 | 8.77 |
| 40 | 3b65 | 7.82 | 9.27 |
| 40 | 3g0w | 7.23 | 9.52 |
| 41 | 4u4s | 5.60 | 2.92 |
| 41 | 1p1q | 6.55 | 4.89 |
| 41 | 1syi | 6.37 | 5.44 |
| 41 | 1p1n | 6.13 | 6.80 |
| 41 | 2al5 | 6.55 | 8.40 |
| 42 | 3g2z | 4.65 | 2.36 |
| 42 | 3g31 | 4.79 | 2.89 |
| 42 | 4de2 | 5.85 | 4.12 |
| 42 | 4de3 | 6.4 | 5.52 |
| 42 | 4de1 | 6.15 | 5.96 |
| 43 | 1vso | 5.51 | 4.72 |
| 43 | 4dld | 6.46 | 5.82 |
| 43 | 3gbb | 6.85 | 6.90 |
| 43 | 3fv2 | 6.67 | 8.11 |
| 43 | 3fv1 | 7.46 | 9.30 |
| 44 | 4mgd | 6.35 | 4.69 |
| 44 | 2qe4 | 7.29 | 7.96 |
| 44 | 1qkt | 7.26 | 9.04 |
| 44 | 2pog | 6.83 | 9.54 |
| 44 | 2p15 | 8.05 | 10.30 |
| 45 | 2y5h | 7.10 | 5.79 |
| 45 | 1lpg | 7.53 | 7.09 |
| 45 | 2xbv | 8.08 | 8.43 |
| 45 | 1z6e | 8.52 | 9.72 |
| 45 | 1mq6 | 8.11 | 11.15 |
| 46 | 1nc3 | 6.02 | 5.00 |
| 46 | 1nc1 | 5.22 | 6.12 |
| 46 | 1y6r | 6.10 | 10.11 |
| 46 | 4f2w | 6.09 | 11.30 |
| 46 | 4f3c | 6.19 | 11.82 |
| 47 | 1uto | 4.10 | 2.27 |
| 47 | 4abg | 5.34 | 3.57 |
| 47 | 3gy4 | 4.81 | 5.10 |
| 47 | 1k1i | 7.59 | 6.58 |
| 47 | 1o3f | 6.55 | 7.96 |
| 48 | 2yge | 7.20 | 5.06 |
| 48 | 2fxs | 6.32 | 6.06 |
| 48 | 2iwx | 6.70 | 6.68 |
| 48 | 2wer | 7.23 | 7.05 |
| 48 | 2vw5 | 7.85 | 8.52 |
| 49 | 4kzq | 5.49 | 6.10 |
| 49 | 4kzu | 5.73 | 6.50 |
| 49 | 4j21 | 5.93 | 7.41 |
| 49 | 4j3l | 6.68 | 7.80 |
| 49 | 3kr8 | 6.33 | 8.10 |
| 50 | 2ymd | 5.24 | 3.16 |
| 50 | 2wnc | 5.65 | 6.32 |
| 50 | 2xys | 6.21 | 7.42 |
| 50 | 2wn9 | 5.91 | 8.52 |
| 50 | 2x00 | 7.64 | 11.33 |
| 51 | 3ozt | 6.19 | 4.13 |
| 51 | 3ozs | 6.93 | 5.33 |
| 51 | 3oe5 | 6.58 | 6.88 |
| 51 | 3oe4 | 6.63 | 7.47 |
| 51 | 3nw9 | 7.75 | 9.00 |
| 52 | 3ao4 | 5.01 | 2.07 |
| 52 | 3zt2 | 5.72 | 2.84 |
| 52 | 3zsx | 5.91 | 3.28 |
| 52 | 4cig | 6.32 | 3.67 |
| 52 | 3zso | 6.67 | 5.12 |
| 53 | 3n7a | 5.2 | 3.70 |
| 53 | 4ciw | 6.15 | 4.82 |
| 53 | 3n86 | 6.27 | 5.64 |
| 53 | 3n76 | 6.51 | 6.85 |
| 53 | 2xb8 | 5.94 | 7.59 |
| 54 | 4bkt | 4.93 | 3.62 |
| 54 | 4w9c | 5.78 | 4.65 |
| 54 | 4w9l | 6.58 | 5.02 |
| 54 | 4w9i | 5.85 | 5.96 |
| 54 | 4w9h | 6.86 | 6.73 |
| 55 | 3nq9 | 4.35 | 4.03 |
| 55 | 3ueu | 4.77 | 5.24 |
| 55 | 3uev | 5.29 | 5.89 |
| 55 | 3uew | 5.72 | 6.31 |
| 55 | 3uex | 5.69 | 6.92 |
| 56 | 3lka | 4.98 | 2.82 |
| 56 | 3ehy | 4.91 | 5.85 |
| 56 | 3tsk | 6.77 | 7.17 |
| 56 | 3nx7 | 5.61 | 8.10 |
| 56 | 4gr0 | 8.83 | 9.55 |
| 57 | 3dxg | 4.47 | 2.40 |
| 57 | 3d6q | 4.29 | 3.76 |
| 57 | 1w4o | 4.69 | 5.22 |
| 57 | 1o0h | 4.74 | 5.92 |
| 57 | 1u1b | 6.59 | 7.80 |

Section 2. Results of the DeepDOX1 ablation experiment on CASF-2016

**Table S2.** Pearson(R_P_) and Spearman(R_S_) correction on CASF-2016 obtained by ablation experiment.

|  | R_P_Pearson | Spearman |
| --- | --- | --- |
| DeepDOX1-dRDkitFeature | 0.65 | 0.64 |
| DeepDOX1-dInteraction | 0.69 | 0.70 |
| DeepDOX1-dAtomicFeature | 0.59 | 0.60 |
| DeepDOX1-dBindingEnergy | 0.65 | 0.67 |
| DeepDOX1-dPocketResidue | 0.60 | 0.66 |

Section 3. Detailed results of the Merk_FEP Test^2^

**Table S3.** Predicted binding affinity against the Merk_FEP set.

| Target | Ligand ID | DeepDOX1 |  |  |  | Expt. |
| --- | --- | --- | --- | --- | --- | --- |
| HIF-2α | HIF-2α_1 | 7.14 | 7.76 | 7.23 | 82.14 | 8.29 |
| HIF-2α | HIF-2α_2 | 7.55 | 7.68 | 7.97 | 80.62 | 8.08 |
| HIF-2α | HIF-2α_3 | 7.06 | 7.60 | 7.31 | 96.76 | 5.17 |
| HIF-2α | HIF-2α_4 | 7.42 | 7.19 | 6.87 | 70.40 | 7.52 |
| HIF-2α | HIF-2α_5 | 7.08 | 7.09 | 7.78 | 91.80 | 7.73 |
| HIF-2α | HIF-2α_6 | 6.60 | 6.79 | 6.68 | 82.27 | 6.47 |
| HIF-2α | HIF-2α_7 | 6.91 | 7.22 | 7.25 | 72.15 | 5.23 |
| HIF-2α | HIF-2α_8 | 6.75 | 6.65 | 7.17 | 75.14 | 7.16 |
| HIF-2α | HIF-2α_9 | 6.78 | 6.45 | 7.23 | 71.98 | 6.54 |
| HIF-2α | HIF-2α_10 | 6.80 | 6.45 | 7.11 | 80.35 | 6.36 |
| HIF-2α | HIF-2α_11 | 6.60 | 7.06 | 7.17 | 63.60 | 7.09 |
| HIF-2α | HIF-2α_12 | 6.88 | 6.79 | 7.09 | 74.56 | 6.81 |
| HIF-2α | HIF-2α_13 | 7.31 | 6.57 | 7.95 | 70.22 | 7.81 |
| HIF-2α | HIF-2α_14 | 6.98 | 7.50 | 6.94 | 75.15 | 7.40 |
| HIF-2α | HIF-2α_15 | 6.77 | 7.94 | 6.26 | 69.99 | 7.44 |
| HIF-2α | HIF-2α_16 | 6.73 | 7.30 | 6.44 | 69.08 | 5.54 |
| HIF-2α | HIF-2α_17 | 6.71 | 6.27 | 7.65 | 70.43 | 7.41 |
| HIF-2α | HIF-2α_18 | 7.33 | 7.28 | 7.60 | 78.86 | 7.69 |
| HIF-2α | HIF-2α_19 | 7.14 | 7.03 | 7.20 | 87.92 | 7.12 |
| HIF-2α | HIF-2α_20 | 7.21 | 7.62 | 7.82 | 65.14 | 8.06 |
| HIF-2α | HIF-2α_21 | 6.74 | 6.60 | 7.66 | 74.55 | 4.95 |
| HIF-2α | HIF-2α_22 | 7.32 | 7.16 | 7.61 | 77.09 | 6.76 |
| HIF-2α | HIF-2α_23 | 7.50 | 7.44 | 7.62 | 81.43 | 7.81 |
| HIF-2α | HIF-2α_24 | 7.37 | 7.08 | 7.22 | 83.58 | 7.41 |
| HIF-2α | HIF-2α_25 | 7.24 | 6.77 | 8.04 | 72.22 | 7.76 |
| HIF-2α | HIF-2α_26 | 7.56 | 6.95 | 6.66 | 85.91 | 7.07 |
| HIF-2α | HIF-2α_27 | 7.11 | 7.74 | 7.75 | 55.86 | 7.87 |
| HIF-2α | HIF-2α_28 | 7.03 | 7.12 | 7.60 | 70.20 | 7.33 |
| HIF-2α | HIF-2α_29 | 7.35 | 6.99 | 7.79 | 63.98 | 7.57 |
| HIF-2α | HIF-2α_30 | 7.56 | 6.88 | 6.74 | 71.94 | 7.41 |
| HIF-2α | HIF-2α_31 | 7.72 | 7.39 | 7.43 | 76.61 | 6.99 |
| HIF-2α | HIF-2α_32 | 7.09 | 7.66 | 7.20 | 89.28 | 7.50 |
| HIF-2α | HIF-2α_33 | 7.35 | 7.28 | 7.82 | 57.88 | 7.11 |
| HIF-2α | HIF-2α_34 | 7.31 | 6.50 | 7.54 | 78.79 | 6.99 |
| HIF-2α | HIF-2α_35 | 7.37 | 8.13 | 7.23 | 70.30 | 7.81 |
| HIF-2α | HIF-2α_36 | 7.87 | 7.28 | 7.42 | 86.95 | 7.71 |
| HIF-2α | HIF-2α_37 | 7.05 | 7.09 | 7.62 | 67.61 | 7.99 |
| HIF-2α | HIF-2α_38 | 7.69 | 6.82 | 7.41 | 72.75 | 7.73 |
| HIF-2α | HIF-2α_39 | 7.20 | 6.88 | 7.64 | 56.85 | 7.19 |
| HIF-2α | HIF-2α_40 | 7.50 | 6.24 | 7.49 | 80.12 | 7.87 |
| HIF-2α | HIF-2α_41 | 7.25 | 6.63 | 7.06 | 83.09 | 7.31 |
| HIF-2α | HIF-2α_42 | 7.19 | 6.32 | 7.08 | 83.38 | 5.41 |
| PFKFB3 | PFKFB3_1 | 7.15 | 7.54 | 6.53 | 85.07 | 5.52 |
| PFKFB3 | PFKFB3_2 | 7.51 | 6.95 | 6.29 | 76.54 | 6.23 |
| PFKFB3 | PFKFB3_3 | 7.43 | 7.45 | 6.16 | 80.77 | 5.79 |
| PFKFB3 | PFKFB3_4 | 7.60 | 8.23 | 6.24 | 86.71 | 6.49 |
| PFKFB3 | PFKFB3_5 | 7.57 | 8.25 | 6.27 | 97.65 | 6.71 |
| PFKFB3 | PFKFB3_6 | 7.87 | 8.08 | 7.08 | 88.79 | 7.06 |
| PFKFB3 | PFKFB3_7 | 7.36 | 7.59 | 6.58 | 84.82 | 6.73 |
| PFKFB3 | PFKFB3_8 | 7.64 | 7.89 | 6.33 | 72.51 | 6.79 |
| PFKFB3 | PFKFB3_9 | 8.01 | 8.02 | 6.56 | 72.48 | 6.30 |
| PFKFB3 | PFKFB3_10 | 7.87 | 8.09 | 5.92 | 77.79 | 7.30 |
| PFKFB3 | PFKFB3_11 | 8.12 | 8.26 | 6.96 | 84.08 | 7.42 |
| PFKFB3 | PFKFB3_12 | 7.86 | 7.37 | 6.59 | 84.87 | 5.44 |
| PFKFB3 | PFKFB3_13 | 8.35 | 8.51 | 6.72 | 83.68 | 7.31 |
| PFKFB3 | PFKFB3_14 | 7.89 | 7.91 | 6.49 | 90.61 | 6.41 |
| PFKFB3 | PFKFB3_15 | 7.51 | 7.38 | 6.32 | 76.84 | 5.26 |
| PFKFB3 | PFKFB3_16 | 7.74 | 3.38 | 6.87 | 68.82 | 5.40 |
| PFKFB3 | PFKFB3_17 | 7.56 | 6.94 | 6.29 | 70.58 | 6.19 |
| PFKFB3 | PFKFB3_18 | 7.98 | 7.80 | 5.95 | 82.24 | 5.85 |
| PFKFB3 | PFKFB3_19 | 8.07 | 7.54 | 5.90 | 89.06 | 7.28 |
| PFKFB3 | PFKFB3_20 | 7.34 | 8.40 | 6.65 | 100.60 | 5.75 |
| PFKFB3 | PFKFB3_21 | 7.79 | 7.25 | 6.47 | 85.42 | 6.10 |
| PFKFB3 | PFKFB3_22 | 7.47 | 7.48 | 6.05 | 94.20 | 5.86 |
| PFKFB3 | PFKFB3_23 | 7.54 | 5.25 | 6.12 | 82.84 | 5.80 |
| PFKFB3 | PFKFB3_24 | 8.12 | 8.25 | 6.74 | 72.10 | 7.61 |
| PFKFB3 | PFKFB3_25 | 8.25 | 7.98 | 7.36 | 73.99 | 5.12 |
| PFKFB3 | PFKFB3_26 | 8.25 | 8.10 | 7.07 | 75.76 | 5.98 |
| PFKFB3 | PFKFB3_27 | 8.36 | 8.69 | 6.69 | 110.13 | 7.78 |
| PFKFB3 | PFKFB3_28 | 8.36 | 8.55 | 7.12 | 91.08 | 7.73 |
| PFKFB3 | PFKFB3_29 | 8.34 | 8.78 | 7.42 | 98.01 | 7.66 |
| PFKFB3 | PFKFB3_30 | 7.95 | 8.17 | 6.34 | 77.56 | 6.68 |
| PFKFB3 | PFKFB3_31 | 7.84 | 7.21 | 6.57 | 97.27 | 5.94 |
| PFKFB3 | PFKFB3_32 | 7.98 | 8.67 | 6.95 | 102.98 | 5.96 |
| PFKFB3 | PFKFB3_33 | 7.73 | 8.34 | 6.52 | 63.95 | 5.63 |
| PFKFB3 | PFKFB3_34 | 7.95 | 8.15 | 6.92 | 66.02 | 6.26 |
| PFKFB3 | PFKFB3_35 | 8.04 | 8.30 | 6.63 | 74.50 | 6.89 |
| PFKFB3 | PFKFB3_36 | 8.21 | 8.01 | 7.10 | 67.59 | 6.94 |
| PFKFB3 | PFKFB3_37 | 7.71 | 8.22 | 6.75 | 81.83 | 7.50 |
| PFKFB3 | PFKFB3_38 | 7.95 | 8.49 | 7.05 | 99.94 | 7.15 |
| PFKFB3 | PFKFB3_39 | 8.23 | 8.43 | 6.77 | 95.71 | 7.84 |
| PFKFB3 | PFKFB3_40 | 8.17 | 8.64 | 6.75 | 87.71 | 7.54 |
| Eg5 | Eg5_1 | 7.20 | 6.72 | 6.16 | 67.08 | 8.08 |
| Eg5 | Eg5_2 | 7.18 | 6.51 | 6.40 | 44.19 | 8.29 |
| Eg5 | Eg5_3 | 6.93 | 7.34 | 7.13 | 72.16 | 7.46 |
| Eg5 | Eg5_4 | 7.09 | 7.49 | 7.08 | 69.01 | 7.76 |
| Eg5 | Eg5_5 | 6.98 | 6.33 | 6.87 | 43.70 | 7.59 |
| Eg5 | Eg5_6 | 7.09 | 6.93 | 6.61 | 54.32 | 8.51 |
| Eg5 | Eg5_7 | 7.27 | 7.71 | 7.19 | 80.69 | 8.21 |
| Eg5 | Eg5_8 | 6.87 | 6.88 | 6.39 | 98.15 | 8.38 |
| Eg5 | Eg5_9 | 6.91 | 7.06 | 6.39 | 73.08 | 7.94 |
| Eg5 | Eg5_10 | 6.86 | 6.83 | 7.00 | 69.18 | 6.91 |
| Eg5 | Eg5_11 | 7.42 | 6.31 | 6.33 | 81.05 | 8.68 |
| Eg5 | Eg5_12 | 7.17 | 7.24 | 6.99 | 76.90 | 7.17 |
| Eg5 | Eg5_13 | 7.22 | 6.87 | 6.36 | 73.39 | 7.84 |
| Eg5 | Eg5_14 | 7.52 | 6.93 | 7.13 | 69.39 | 8.38 |
| Eg5 | Eg5_15 | 7.47 | 6.26 | 6.50 | 61.48 | 8.14 |
| Eg5 | Eg5_16 | 7.21 | 7.20 | 6.54 | 65.98 | 7.71 |
| Eg5 | Eg5_17 | 6.81 | 7.42 | 6.40 | 67.84 | 7.69 |
| Eg5 | Eg5_18 | 7.51 | 6.57 | 6.45 | 77.99 | 8.29 |
| Eg5 | Eg5_19 | 7.35 | 6.53 | 6.56 | 61.57 | 7.94 |
| Eg5 | Eg5_20 | 7.14 | 7.55 | 6.16 | 67.90 | 7.22 |
| Eg5 | Eg5_21 | 7.07 | 7.75 | 6.53 | 67.26 | 8.38 |
| Eg5 | Eg5_22 | 6.34 | 6.63 | 6.22 | 63.87 | 6.36 |
| Eg5 | Eg5_23 | 6.71 | 6.63 | 6.89 | 44.33 | 6.91 |
| Eg5 | Eg5_24 | 6.34 | 6.13 | 6.13 | 56.43 | 6.13 |
| Eg5 | Eg5_25 | 7.27 | 7.02 | 6.65 | 70.32 | 7.71 |
| Eg5 | Eg5_26 | 7.04 | 6.21 | 6.66 | 62.93 | 8.21 |
| Eg5 | Eg5_27 | 6.98 | 6.86 | 6.49 | 68.57 | 8.14 |
| Eg5 | Eg5_28 | 7.21 | 5.90 | 7.29 | 71.16 | 8.68 |
| CDK8 | CDK8_1 | 7.57 | 6.96 | 6.96 | 102.79 | 8.69 |
| CDK8 | CDK8_2 | 6.53 | 4.14 | 7.16 | 93.10 | 6.54 |
| CDK8 | CDK8_3 | 6.42 | 6.08 | 7.32 | 68.36 | 4.51 |
| CDK8 | CDK8_4 | 7.70 | 6.21 | 7.35 | 56.89 | 7.48 |
| CDK8 | CDK8_5 | 6.09 | 5.12 | 6.84 | 56.87 | 6.76 |
| CDK8 | CDK8_6 | 7.14 | 7.06 | 7.95 | 73.69 | 8.51 |
| CDK8 | CDK8_7 | 7.57 | 5.20 | 7.47 | 71.93 | 7.10 |
| CDK8 | CDK8_8 | 6.61 | 6.09 | 7.27 | 77.25 | 7.52 |
| CDK8 | CDK8_9 | 7.54 | 7.10 | 7.58 | 82.12 | 8.28 |
| CDK8 | CDK8_10 | 7.00 | 7.12 | 7.93 | 78.79 | 8.69 |
| CDK8 | CDK8_11 | 7.79 | 6.41 | 7.68 | 81.42 | 8.21 |
| CDK8 | CDK8_12 | 7.47 | 6.52 | 7.30 | 76.27 | 6.73 |
| CDK8 | CDK8_13 | 7.62 | 6.48 | 7.62 | 69.73 | 7.43 |
| CDK8 | CDK8_14 | 7.25 | 6.31 | 7.80 | 65.40 | 7.79 |
| CDK8 | CDK8_15 | 6.64 | 6.77 | 7.05 | 94.28 | 8.39 |
| CDK8 | CDK8_16 | 6.81 | 6.81 | 7.86 | 70.13 | 8.39 |
| CDK8 | CDK8_17 | 6.81 | 6.21 | 7.27 | 93.56 | 8.39 |
| CDK8 | CDK8_18 | 7.05 | 5.87 | 7.22 | 83.90 | 7.98 |
| CDK8 | CDK8_19 | 7.30 | 6.19 | 7.77 | 75.86 | 8.14 |
| CDK8 | CDK8_20 | 6.21 | 6.08 | 7.12 | 58.86 | 7.81 |
| CDK8 | CDK8_21 | 7.68 | 6.09 | 7.44 | 79.31 | 8.09 |
| CDK8 | CDK8_22 | 7.22 | 5.73 | 7.32 | 79.57 | 6.15 |
| CDK8 | CDK8_23 | 6.21 | 6.33 | 7.14 | 45.47 | 8.21 |
| CDK8 | CDK8_24 | 7.64 | 6.42 | 7.48 | 82.43 | 8.39 |
| CDK8 | CDK8_25 | 6.17 | 5.70 | 7.18 | 61.19 | 6.94 |
| CDK8 | CDK8_26 | 7.09 | 7.02 | 7.46 | 73.85 | 8.39 |
| CDK8 | CDK8_27 | 7.89 | 5.27 | 7.59 | 73.20 | 6.56 |
| CDK8 | CDK8_28 | 6.19 | 5.70 | 6.94 | 62.18 | 5.59 |
| CDK8 | CDK8_29 | 7.61 | 6.44 | 7.60 | 83.81 | 8.14 |
| CDK8 | CDK8_30 | 6.56 | 5.96 | 7.47 | 72.91 | 7.13 |
| CDK8 | CDK8_31 | 5.70 | 5.03 | 6.31 | 71.82 | 6.06 |
| CDK8 | CDK8_32 | 6.89 | 5.31 | 7.49 | 99.02 | 8.14 |
| CDK8 | CDK8_33 | 7.59 | 6.48 | 7.39 | 85.62 | 8.21 |
| SHP-2 | SHP-2_1 | 6.98 | 7.34 | 6.31 | 60.57 | 7.14 |
| SHP-2 | SHP-2_2 | 7.65 | 7.23 | 6.74 | 64.64 | 6.76 |
| SHP-2 | SHP-2_3 | 6.82 | 6.75 | 5.82 | 46.62 | 5.58 |
| SHP-2 | SHP-2_4 | 6.98 | 4.19 | 6.36 | 60.55 | 4.65 |
| SHP-2 | SHP-2_5 | 6.98 | 0.10 | 5.95 | 50.71 | 4.26 |
| SHP-2 | SHP-2_6 | 7.10 | 7.02 | 6.16 | 53.62 | 6.05 |
| SHP-2 | SHP-2_7 | 6.85 | 6.54 | 6.12 | 65.09 | 7.15 |
| SHP-2 | SHP-2_8 | 7.18 | 7.60 | 6.76 | 60.80 | 6.55 |
| SHP-2 | SHP-2_9 | 7.03 | 0.45 | 6.08 | 61.75 | 4.91 |
| SHP-2 | SHP-2_10 | 6.98 | 7.67 | 6.74 | 60.54 | 6.64 |
| SHP-2 | SHP-2_11 | 6.90 | 7.48 | 6.46 | 60.06 | 6.58 |
| SHP-2 | SHP-2_12 | 6.67 | 0.06 | 5.66 | 54.09 | 3.99 |
| SHP-2 | SHP-2_13 | 7.00 | 6.49 | 6.39 | 61.66 | 6.43 |
| SHP-2 | SHP-2_14 | 7.15 | 6.64 | 6.28 | 56.09 | 7.11 |
| SHP-2 | SHP-2_15 | 6.76 | 3.93 | 6.30 | 61.63 | 5.18 |
| SHP-2 | SHP-2_16 | 7.27 | 7.04 | 6.04 | 66.81 | 6.64 |
| SHP-2 | SHP-2_17 | 6.92 | 3.54 | 6.14 | 26.55 | 6.19 |
| SHP-2 | SHP-2_18 | 7.02 | 7.30 | 6.07 | 60.29 | 5.23 |
| SHP-2 | SHP-2_19 | 7.24 | 7.58 | 6.79 | 67.93 | 6.49 |
| SHP-2 | SHP-2_20 | 7.04 | 0.46 | 6.19 | 67.93 | 6.23 |
| SHP-2 | SHP-2_21 | 6.79 | 7.34 | 6.04 | 53.88 | 5.87 |
| SHP-2 | SHP-2_22 | 7.09 | 7.60 | 6.36 | 62.17 | 6.70 |
| SHP-2 | SHP-2_23 | 6.71 | 4.88 | 6.60 | 58.26 | 5.53 |
| SHP-2 | SHP-2_24 | 6.49 | 6.28 | 5.72 | 44.47 | 6.00 |
| SHP-2 | SHP-2_25 | 7.15 | 7.35 | 6.80 | 65.78 | 6.39 |
| SHP-2 | SHP-2_26 | 6.57 | 6.45 | 5.90 | 53.05 | 6.01 |
| SYK | SYK_1 | 7.46 | 6.27 | 8.19 | 248.89 | 8.36 |
| SYK | SYK_2 | 6.93 | 5.30 | 7.59 | 235.44 | 7.61 |
| SYK | SYK_3 | 7.55 | 6.44 | 7.42 | 203.06 | 8.41 |
| SYK | SYK_4 | 7.05 | 6.29 | 7.81 | 233.04 | 6.94 |
| SYK | SYK_5 | 7.09 | 6.49 | 7.57 | 219.64 | 7.63 |
| SYK | SYK_6 | 7.16 | 6.52 | 7.37 | 210.03 | 8.06 |
| SYK | SYK_7 | 7.20 | 5.72 | 7.97 | 226.33 | 7.67 |
| SYK | SYK_8 | 8.03 | 6.45 | 7.90 | 251.85 | 7.77 |
| SYK | SYK_9 | 7.33 | 6.64 | 7.93 | 211.41 | 8.67 |
| SYK | SYK_10 | 6.42 | 5.69 | 7.07 | 201.42 | 7.56 |
| SYK | SYK_11 | 7.03 | 6.73 | 7.69 | 217.88 | 8.06 |
| SYK | SYK_12 | 7.36 | 6.42 | 8.27 | 244.97 | 8.04 |
| SYK | SYK_13 | 7.50 | 6.63 | 7.71 | 247.11 | 7.81 |
| SYK | SYK_14 | 7.14 | 5.19 | 8.32 | 220.30 | 7.18 |
| SYK | SYK_15 | 7.39 | 5.75 | 8.18 | 243.64 | 9.06 |
| SYK | SYK_16 | 7.36 | 6.92 | 7.45 | 206.48 | 8.10 |
| SYK | SYK_17 | 7.76 | 6.58 | 7.96 | 207.48 | 8.84 |
| SYK | SYK_18 | 7.75 | 5.41 | 8.21 | 211.86 | 9.34 |
| SYK | SYK_19 | 7.08 | 6.86 | 7.43 | 242.73 | 8.32 |
| SYK | SYK_20 | 7.04 | 5.88 | 7.86 | 242.62 | 7.72 |
| SYK | SYK_21 | 7.98 | 6.49 | 7.90 | 252.24 | 7.92 |
| SYK | SYK_22 | 7.48 | 6.67 | 8.23 | 242.47 | 8.58 |
| SYK | SYK_23 | 7.01 | 6.41 | 7.69 | 209.46 | 8.14 |
| SYK | SYK_24 | 7.24 | 6.25 | 8.08 | 216.44 | 8.11 |
| SYK | SYK_25 | 7.15 | 6.86 | 7.81 | 231.15 | 8.63 |
| SYK | SYK_26 | 7.03 | 5.54 | 7.87 | 221.34 | 8.19 |
| SYK | SYK_27 | 6.91 | 6.83 | 7.86 | 244.17 | 7.16 |
| SYK | SYK_28 | 7.13 | 1.61 | 7.68 | 209.75 | 8.56 |
| SYK | SYK_29 | 6.57 | 6.45 | 7.60 | 243.50 | 7.47 |
| SYK | SYK_30 | 6.61 | 6.27 | 7.82 | 233.11 | 7.18 |
| SYK | SYK_31 | 6.92 | 5.93 | 7.13 | 66.39 | 7.74 |
| SYK | SYK_32 | 7.81 | 6.85 | 8.08 | 254.04 | 8.81 |
| SYK | SYK_33 | 6.46 | 7.08 | 7.60 | 240.83 | 7.29 |
| SYK | SYK_34 | 7.06 | 6.78 | 7.83 | 202.39 | 8.39 |
| SYK | SYK_35 | 7.22 | 7.08 | 7.82 | 215.55 | 7.90 |
| SYK | SYK_36 | 7.11 | 5.79 | 7.81 | 217.16 | 7.89 |
| SYK | SYK_37 | 8.10 | 6.09 | 7.91 | 247.52 | 7.95 |
| SYK | SYK_38 | 6.68 | 6.93 | 7.87 | 246.83 | 7.54 |
| SYK | SYK_39 | 7.34 | 6.59 | 7.88 | 222.44 | 7.97 |
| SYK | SYK_40 | 6.97 | 6.95 | 7.81 | 229.51 | 6.32 |
| SYK | SYK_41 | 6.43 | 6.83 | 7.83 | 238.13 | 7.56 |
| SYK | SYK_42 | 7.33 | 6.69 | 7.86 | 221.12 | 8.41 |
| SYK | SYK_43 | 7.36 | 3.38 | 8.17 | 241.96 | 8.65 |
| SYK | SYK_44 | 7.00 | 6.04 | 7.68 | 240.14 | 7.42 |
| c-MET | c-MET_1 | 6.71 | 8.11 | 7.23 | 68.78 | 6.69 |
| c-MET | c-MET_2 | 6.88 | 7.62 | 7.28 | 69.13 | 8.23 |
| c-MET | c-MET_3 | 6.40 | 7.52 | 7.22 | 88.72 | 5.46 |
| c-MET | c-MET_4 | 6.64 | 7.08 | 7.46 | 145.92 | 6.51 |
| c-MET | c-MET_5 | 6.95 | 7.32 | 7.64 | 133.19 | 6.69 |
| c-MET | c-MET_6 | 7.61 | 8.33 | 7.91 | 80.88 | 8.99 |
| c-MET | c-MET_7 | 6.54 | 6.88 | 7.90 | 158.15 | 6.39 |
| c-MET | c-MET_8 | 6.80 | 7.87 | 7.55 | 140.79 | 7.36 |
| c-MET | c-MET_9 | 6.55 | 6.73 | 7.05 | 79.90 | 4.51 |
| c-MET | c-MET_10 | 7.12 | 7.79 | 7.37 | 69.46 | 5.37 |
| c-MET | c-MET_11 | 6.93 | 7.26 | 7.07 | 63.57 | 7.98 |
| c-MET | c-MET_12 | 6.81 | 7.17 | 7.60 | 87.25 | 6.29 |
| c-MET | c-MET_13 | 7.62 | 7.51 | 8.07 | 60.25 | 8.17 |
| c-MET | c-MET_14 | 7.26 | 7.86 | 8.03 | 145.32 | 7.95 |
| c-MET | c-MET_15 | 7.48 | 8.09 | 7.63 | 78.20 | 8.99 |
| c-MET | c-MET_16 | 7.33 | 7.48 | 7.63 | 136.08 | 7.62 |
| c-MET | c-MET_17 | 7.15 | 7.72 | 7.61 | 89.98 | 7.38 |
| c-MET | c-MET_18 | 6.62 | 6.81 | 7.32 | 64.49 | 5.27 |
| c-MET | c-MET_19 | 7.36 | 7.56 | 7.48 | 96.74 | 6.39 |
| c-MET | c-MET_20 | 7.41 | 6.97 | 7.49 | 153.31 | 7.03 |
| c-MET | c-MET_21 | 6.83 | 7.03 | 7.00 | 132.93 | 7.03 |
| c-MET | c-MET_22 | 7.18 | 7.31 | 7.33 | 69.48 | 5.67 |
| c-MET | c-MET_23 | 7.54 | 7.57 | 8.25 | 68.63 | 8.55 |
| c-MET | c-MET_24 | 7.64 | 7.91 | 8.06 | 73.11 | 8.99 |
| TNKS2 | TNKS2_1 | 6.28 | 6.80 | 7.61 | 229.27 | 8.05 |
| TNKS2 | TNKS2_2 | 6.55 | 6.78 | 8.01 | 252.59 | 7.51 |
| TNKS2 | TNKS2_3 | 6.95 | 7.94 | 8.17 | 224.89 | 7.49 |
| TNKS2 | TNKS2_4 | 6.62 | 7.30 | 8.16 | 249.98 | 7.38 |
| TNKS2 | TNKS2_5 | 6.55 | 8.00 | 8.70 | 240.87 | 8.73 |
| TNKS2 | TNKS2_6 | 6.55 | 6.72 | 7.94 | 231.45 | 7.91 |
| TNKS2 | TNKS2_7 | 5.79 | 6.03 | 7.61 | 226.12 | 7.27 |
| TNKS2 | TNKS2_8 | 6.62 | 6.35 | 7.86 | 227.11 | 7.40 |
| TNKS2 | TNKS2_9 | 6.93 | 6.44 | 7.98 | 177.33 | 8.84 |
| TNKS2 | TNKS2_10 | 6.53 | 8.26 | 8.78 | 249.10 | 9.14 |
| TNKS2 | TNKS2_11 | 6.43 | 6.28 | 8.01 | 246.31 | 7.66 |
| TNKS2 | TNKS2_12 | 6.51 | 6.99 | 7.80 | 224.32 | 7.35 |
| TNKS2 | TNKS2_13 | 6.97 | -1.03 | 7.97 | 233.51 | 7.40 |
| TNKS2 | TNKS2_14 | 6.20 | 6.34 | 7.74 | 238.05 | 7.87 |
| TNKS2 | TNKS2_15 | 7.11 | 7.24 | 7.67 | 215.37 | 8.02 |
| TNKS2 | TNKS2_16 | 6.36 | 6.48 | 8.02 | 157.88 | 6.15 |
| TNKS2 | TNKS2_17 | 6.68 | 6.71 | 7.44 | 222.10 | 7.68 |
| TNKS2 | TNKS2_18 | 5.46 | 5.85 | 6.39 | 221.14 | 6.26 |
| TNKS2 | TNKS2_19 | 6.70 | 7.05 | 7.73 | 238.35 | 8.11 |
| TNKS2 | TNKS2_20 | 6.68 | 8.22 | 8.82 | 238.30 | 8.64 |
| TNKS2 | TNKS2_21 | 6.41 | 7.00 | 7.59 | 231.68 | 8.43 |
| TNKS2 | TNKS2_22 | 6.14 | 7.76 | 8.11 | 236.41 | 7.91 |
| TNKS2 | TNKS2_23 | 6.93 | 2.84 | 7.83 | 235.96 | 8.81 |
| TNKS2 | TNKS2_24 | 6.84 | 6.79 | 7.98 | 217.72 | 7.97 |
| TNKS2 | TNKS2_25 | 7.05 | 6.81 | 7.43 | 240.11 | 9.29 |
| TNKS2 | TNKS2_26 | 6.75 | 6.85 | 7.74 | 226.09 | 7.84 |
| TNKS2 | TNKS2_27 | 6.82 | 5.74 | 8.11 | 222.33 | 7.29 |

Section 4. Establishment of the HLO-2025 set and detailed results

To address potential limitations in the generalizability assessment using existing benchmarks such as Merk_FEP and CASF-2016, we constructed a novel binding affinity dataset comprising both covalent and non-covalent targets. The dataset includes one covalent target and twelve non-covalent targets, each containing at least 20 small molecules with activity values spanning over four orders of magnitude. DeepDOX1 results and activity data are provided in the table below.

**Table S4.** Predicted binding affinity against the HLO-2025 test set

| Target | Ligand ID | DeepDOX1 | SableBind | ∆_vina_XGB | TankBind | DeepDOCK | Expt. |
| --- | --- | --- | --- | --- | --- | --- | --- |
| AKT1^3^ | AKT1_1 | 6.03 | 6.36 | 5.59 | 4.54 | 53.50 | 6.13 |
| AKT1 | AKT1_2 | 8.44 | 7.55 | 5.60 | 6.88 | 63.23 | 7.04 |
| AKT1 | AKT1_3 | 7.11 | 8.08 | 5.05 | 5.97 | 66.86 | 7.04 |
| AKT1 | AKT1_4 | 8.81 | 7.69 | 5.97 | 6.32 | 61.05 | 5.37 |
| AKT1 | AKT1_5 | 7.06 | 6.85 | 5.81 | 5.31 | 74.14 | 5.60 |
| AKT1 | AKT1_6 | 8.07 | 8.04 | 4.99 | 6.28 | 77.36 | 4.92 |
| AKT1 | AKT1_7 | 6.41 | 7.23 | 4.64 | 3.07 | 56.14 | 5.88 |
| AKT1 | AKT1_8 | 7.04 | 8.26 | 5.61 | 5.35 | 83.76 | 6.38 |
| AKT1 | AKT1_9 | 7.09 | 7.86 | 0.56 | 6.63 | 86.59 | 6.76 |
| AKT1 | AKT1_10 | 6.56 | 6.30 | 4.70 | 4.68 | 34.31 | 5.05 |
| AKT1 | AKT1_11 | 8.35 | 7.38 | 7.87 | 5.36 | 66.80 | 7.08 |
| AKT1 | AKT1_12 | 8.38 | 7.94 | 7.37 | 5.00 | 55.84 | 8.77 |
| AKT1 | AKT1_13 | 7.37 | 8.15 | 3.59 | 6.12 | 49.21 | 6.08 |
| AKT1 | AKT1_14 | 6.04 | 7.96 | 4.98 | 5.62 | 40.08 | 3.52 |
| AKT1 | AKT1_15 | 6.22 | 7.61 | 4.98 | 4.89 | 53.41 | 6.03 |
| AKT1 | AKT1_16 | 7.16 | 7.92 | 3.66 | 5.98 | 68.74 | 7.29 |
| AKT1 | AKT1_17 | 7.41 | 7.91 | 4.09 | 6.64 | 61.38 | 7.48 |
| AKT1 | AKT1_18 | 7.99 | 7.90 | 5.33 | 6.71 | 57.28 | 8.32 |
| AKT1 | AKT1_19 | 7.79 | 7.35 | 5.60 | 6.53 | 47.16 | 8.20 |
| AKT1 | AKT1_20 | 7.59 | 7.58 | 7.04 | 6.24 | 62.02 | 8.85 |
| AKT1 | AKT1_21 | 7.99 | 7.77 | 1.72 | 5.52 | 42.02 | 8.84 |
| AKT1 | AKT1_22 | 7.86 | 8.27 | 6.59 | 6.55 | 77.65 | 8.61 |
| AKT1 | AKT1_23 | 6.42 | 7.75 | 7.80 | 6.40 | 60.21 | 8.95 |
| AKT1 | AKT1_24 | 8.19 | 7.82 | 6.24 | 6.77 | 45.04 | 8.42 |
| AKT1 | AKT1_25 | 6.06 | 7.51 | 5.37 | 5.62 | 36.90 | 4.88 |
| AKT1 | AKT1_26 | 8.01 | 7.96 | 5.13 | 6.03 | 60.68 | 7.32 |
| AKT1 | AKT1_27 | 6.15 | 7.35 | 5.07 | 3.72 | 31.85 | 5.40 |
| AKT1 | AKT1_28 | 8.92 | 8.31 | 6.93 | 6.21 | 57.88 | 8.51 |
| AKT1 | AKT1_29 | 8.66 | 7.88 | 4.71 | 6.48 | 60.19 | 9.15 |
| AKT1 | AKT1_30 | 8.58 | 8.25 | 8.36 | 6.87 | 77.76 | 8.92 |
| AKT1 | AKT1_31 | 8.96 | 7.63 | 6.50 | 6.44 | 66.51 | 9.69 |
| AKT1 | AKT1_32 | 8.87 | 8.53 | 6.54 | 5.18 | 45.39 | 8.13 |
| AKT1 | AKT1_33 | 6.28 | 7.89 | 6.37 | 5.95 | 35.12 | 6.05 |
| AKT1 | AKT1_34 | 7.88 | 7.94 | 4.91 | 7.05 | 63.47 | 8.30 |
| AKT1 | AKT1_35 | 7.25 | 7.72 | 5.20 | 7.16 | 67.16 | 7.79 |
| AKT1 | AKT1_36 | 7.88 | 8.00 | 7.91 | 6.19 | 57.18 | 8.50 |
| AKT1 | AKT1_37 | 9.30 | 7.75 | 8.73 | 7.27 | 55.19 | 9.09 |
| AKT1 | AKT1_38 | 7.59 | 7.80 | 5.95 | 6.41 | 40.47 | 8.36 |
| AKT1 | AKT1_39 | 7.62 | 8.03 | 6.21 | 5.94 | 50.90 | 8.24 |
| AKT1 | AKT1_40 | 8.00 | 8.09 | 5.18 | 6.32 | 38.08 | 7.44 |
| AKT1 | AKT1_41 | 7.74 | 8.22 | 7.18 | 6.18 | 35.12 | 6.96 |
| AKT1 | AKT1_42 | 8.13 | 8.44 | 6.57 | 6.17 | 53.63 | 8.95 |
| AKT1 | AKT1_43 | 8.83 | 8.12 | 7.47 | 5.96 | 60.36 | 9.39 |
| AKT1 | AKT1_44 | 7.97 | 8.17 | 4.40 | 5.31 | 29.94 | 8.05 |
| AKT1 | AKT1_45 | 8.59 | 8.22 | 5.95 | 5.34 | 36.78 | 8.14 |
| AKT1 | AKT1_46 | 6.80 | 7.90 | 4.27 | 4.22 | 53.91 | 7.63 |
| AKT1 | AKT1_47 | 9.38 | 8.68 | 5.78 | 6.52 | 67.70 | 8.76 |
| AKT1 | AKT1_48 | 6.68 | 7.76 | 5.54 | 5.17 | 59.73 | 5.88 |
| AKT1 | AKT1_49 | 8.54 | 8.15 | 5.01 | 5.84 | 45.86 | 9.69 |
| AKT1 | AKT1_50 | 7.44 | 7.97 | 5.50 | 5.84 | 40.93 | 6.18 |
| AKT1 | AKT1_51 | 6.80 | 8.07 | 6.39 | 6.35 | 38.06 | 8.11 |
| AKT1 | AKT1_52 | 7.41 | 8.05 | 6.64 | 6.51 | 42.47 | 5.27 |
| AKT1 | AKT1_53 | 6.97 | 8.05 | 6.00 | 5.84 | 45.92 | 6.60 |
| AKT1 | AKT1_54 | 7.19 | 7.82 | 5.87 | 5.48 | 56.59 | 7.45 |
| AKT1 | AKT1_55 | 6.77 | 7.53 | 5.84 | 5.78 | 51.07 | 6.58 |
| AKT1 | AKT1_56 | 8.13 | 8.02 | 4.78 | 5.29 | 48.95 | 5.93 |
| AKT1 | AKT1_57 | 6.75 | 7.27 | 4.87 | 3.33 | 67.14 | 3.52 |
| AKT1 | AKT1_58 | 7.46 | 7.82 | 7.73 | 5.38 | 58.22 | 8.85 |
| AKT1 | AKT1_59 | 6.53 | 7.65 | 5.62 | 4.91 | 52.03 | 3.52 |
| AKT1 | AKT1_60 | 6.51 | 7.64 | 4.98 | 5.62 | 40.13 | 3.73 |
| AKT1 | AKT1_61 | 6.28 | 6.47 | 4.72 | 4.56 | 50.98 | 6.14 |
| AKT1 | AKT1_62 | 8.00 | 7.82 | 6.41 | 4.64 | 58.66 | 7.45 |
| AKT1 | AKT1_63 | 7.07 | 8.05 | 5.30 | 6.46 | 56.51 | 7.89 |
| AKT1 | AKT1_64 | 7.93 | 8.28 | 5.83 | 5.26 | 47.51 | 8.26 |
| AKT1 | AKT1_65 | 7.43 | 8.02 | 5.63 | 7.14 | 42.92 | 8.24 |
| AKT1 | AKT1_66 | 6.31 | 7.99 | 6.19 | 6.38 | 38.27 | 8.36 |
| AKT1 | AKT1_67 | 7.56 | 8.05 | 5.39 | 6.47 | 55.48 | 8.23 |
| AKT1 | AKT1_68 | 6.81 | 8.96 | 2.95 | 6.16 | 72.58 | 9.69 |
| AKT1 | AKT1_69 | 7.17 | 7.18 | 5.46 | 6.18 | 45.79 | 8.32 |
| AKT1 | AKT1_70 | 7.08 | 8.10 | 6.22 | 6.47 | 45.66 | 7.93 |
| AKT1 | AKT1_71 | 7.72 | 7.87 | 4.78 | 6.49 | 45.85 | 8.38 |
| AKT1 | AKT1_72 | 6.92 | 7.92 | 6.08 | 6.84 | 44.43 | 8.35 |
| AKT1 | AKT1_73 | 7.77 | 7.95 | 5.95 | 6.35 | 39.58 | 8.22 |
| AKT1 | AKT1_74 | 6.99 | 8.51 | 6.07 | 5.83 | 61.06 | 6.26 |
| AKT1 | AKT1_75 | 8.17 | 8.03 | 4.13 | 5.42 | 52.33 | 9.25 |
| USP7^4^ | USP7_1 | 6.51 | 6.17 | 7.41 | 4.36 | 36.95 | 5.25 |
| USP7 | USP7_2 | 6.77 | 6.07 | -6.26 | 4.62 | 25.41 | 6.03 |
| USP7 | USP7_3 | 7.11 | 6.90 | 7.08 | 4.31 | 25.53 | 6.88 |
| USP7 | USP7_4 | 7.32 | 6.80 | 6.00 | 4.52 | 31.98 | 7.36 |
| USP7 | USP7_5 | 7.47 | 6.32 | -16.15 | 5.10 | 23.62 | 7.13 |
| USP7 | USP7_6 | 7.53 | 6.94 | 3.75 | 5.07 | 30.85 | 8.04 |
| USP7 | USP7_7 | 7.79 | 6.93 | -2.98 | 6.65 | 35.29 | 7.56 |
| USP7 | USP7_8 | 7.59 | 7.29 | -0.02 | 4.99 | 29.25 | 7.60 |
| USP7 | USP7_9 | 7.74 | 7.07 | 7.32 | 5.78 | 38.45 | 7.58 |
| USP7 | USP7_10 | 7.67 | 7.05 | 7.51 | 4.99 | 33.38 | 9.74 |
| USP7 | USP7_11 | 7.26 | 6.71 | 3.90 | 5.10 | 36.27 | 6.68 |
| USP7 | USP7_12 | 7.91 | 6.31 | 6.57 | 5.33 | 28.22 | 8.09 |
| USP7 | USP7_13 | 7.44 | 6.96 | 6.45 | 5.71 | 33.82 | 8.82 |
| USP7 | USP7_14 | 7.53 | 6.67 | 7.33 | 5.28 | 34.52 | 8.46 |
| USP7 | USP7_15 | 7.67 | 6.70 | 7.35 | 5.72 | 28.39 | 8.67 |
| USP7 | USP7_16 | 8.14 | 6.33 | 8.81 | 5.31 | 29.87 | 8.95 |
| USP7 | USP7_17 | 8.35 | 6.54 | 7.20 | 6.04 | 42.04 | 9.74 |
| USP7 | USP7_18 | 8.31 | 7.22 | 6.43 | 5.84 | 36.44 | 8.61 |
| USP7 | USP7_19 | 8.43 | 7.18 | 6.71 | 6.16 | 37.34 | 9.10 |
| USP7 | USP7_20 | 8.35 | 6.84 | 5.80 | 6.01 | 39.73 | 8.88 |
| USP7 | USP7_21 | 7.58 | 6.35 | 6.95 | 5.06 | 38.99 | 7.49 |
| USP7 | USP7_22 | 7.64 | 6.81 | 6.13 | 5.41 | 41.33 | 8.18 |
| USP7 | USP7_23 | 7.83 | 6.64 | 5.71 | 5.50 | 38.33 | 9.49 |
| USP7 | USP7_24 | 7.76 | 7.21 | 8.18 | 5.06 | 43.52 | 9.79 |
| USP7 | USP7_25 | 7.97 | 7.33 | 7.93 | 5.16 | 46.09 | 8.82 |
| USP7 | USP7_26 | 7.96 | 7.41 | 8.21 | 5.61 | 49.87 | 9.40 |
| USP7 | USP7_27 | 7.63 | 6.72 | -2.87 | 5.43 | 31.75 | 9.35 |
| MNK2^5^ | MNK2_1 | 4.94 | 5.83 | 4.65 | 5.40 | 59.70 | 5.44 |
| MNK2 | MNK2_2 | 5.07 | 6.21 | 4.40 | 5.75 | 57.29 | 5.52 |
| MNK2 | MNK2_3 | 5.28 | 5.99 | 4.32 | 6.25 | 55.56 | 5.16 |
| MNK2 | MNK2_4 | 5.14 | 6.01 | 4.40 | 6.16 | 61.33 | 4.88 |
| MNK2 | MNK2_5 | 5.24 | 6.55 | 2.38 | 5.91 | 50.21 | 6.00 |
| MNK2 | MNK2_6 | 5.22 | 6.54 | 4.61 | 5.94 | 59.79 | 5.14 |
| MNK2 | MNK2_7 | 5.69 | 6.80 | -0.44 | 5.93 | 55.97 | 6.09 |
| MNK2 | MNK2_8 | 5.54 | 6.79 | -2.73 | 6.15 | 40.52 | 5.29 |
| MNK2 | MNK2_9 | 5.41 | 6.64 | 1.98 | 6.51 | 51.16 | 6.46 |
| MNK2 | MNK2_10 | 5.56 | 7.02 | 4.89 | 6.82 | 51.99 | 6.13 |
| MNK2 | MNK2_11 | 5.76 | 6.74 | -0.75 | 6.76 | 67.81 | 6.52 |
| MNK2 | MNK2_12 | 5.27 | 6.51 | -0.88 | 6.38 | 47.97 | 6.33 |
| MNK2 | MNK2_13 | 5.78 | 6.77 | 2.77 | 6.58 | 59.38 | 5.74 |
| MNK2 | MNK2_14 | 6.15 | 6.97 | 4.63 | 6.80 | 52.24 | 5.92 |
| MNK2 | MNK2_15 | 5.24 | 6.33 | 4.72 | 5.91 | 52.11 | 5.82 |
| MNK2 | MNK2_16 | 5.37 | 6.29 | 4.81 | 6.10 | 42.29 | 5.19 |
| MNK2 | MNK2_17 | 5.31 | 6.24 | 5.06 | 6.21 | 44.06 | 5.85 |
| MNK2 | MNK2_18 | 5.08 | 5.77 | 3.53 | 5.91 | 50.42 | 3.96 |
| MNK2 | MNK2_19 | 4.66 | 6.18 | 4.68 | 6.21 | 50.09 | 4.36 |
| MNK2 | MNK2_20 | 5.32 | 6.37 | 4.81 | 6.11 | 53.26 | 5.48 |
| MNK2 | MNK2_21 | 5.32 | 6.37 | 4.81 | 6.11 | 53.26 | 6.38 |
| MNK2 | MNK2_22 | 6.08 | 6.90 | -22.97 | 7.14 | 29.63 | 6.50 |
| MNK2 | MNK2_23 | 5.92 | 6.99 | -22.26 | 7.03 | 26.73 | 6.45 |
| MNK2 | MNK2_24 | 6.11 | 7.53 | 1.75 | 7.60 | 48.21 | 6.79 |
| MNK2 | MNK2_25 | 6.87 | 7.77 | 1.94 | 7.74 | 48.55 | 7.09 |
| MNK2 | MNK2_26 | 5.99 | 6.67 | 2.86 | 7.22 | 41.55 | 6.30 |
| MNK2 | MNK2_27 | 5.82 | 7.44 | -0.33 | 7.00 | 36.35 | 6.40 |
| MNK2 | MNK2_28 | 4.99 | 7.67 | -1.73 | 6.88 | 41.40 | 6.20 |
| MNK2 | MNK2_29 | 6.99 | 7.89 | 2.01 | 7.67 | 43.94 | 7.37 |
| JAK^6^ | JAK_1 | 7.17 | 8.66 | 2.07 | 8.08 | 26.09 | 6.29 |
| JAK | JAK_2 | 7.04 | 7.55 | 1.74 | 7.21 | 20.92 | 5.84 |
| JAK | JAK_3 | 5.77 | 7.26 | 2.73 | 6.98 | 25.28 | 6.31 |
| JAK | JAK_4 | 5.98 | 6.83 | 2.70 | 6.90 | 18.94 | 6.49 |
| JAK | JAK_5 | 5.95 | 7.42 | 1.62 | 7.34 | 24.67 | 6.11 |
| JAK | JAK_6 | 8.04 | 8.01 | 8.71 | 8.39 | 96.58 | 9.52 |
| JAK | JAK_7 | 7.76 | 7.67 | 7.43 | 7.54 | 91.93 | 8.26 |
| JAK | JAK_8 | 6.39 | 6.89 | 5.87 | 7.35 | 94.17 | 6.99 |
| JAK | JAK_9 | 5.99 | 7.65 | 3.11 | 7.00 | 26.73 | 5.00 |
| JAK | JAK_10 | 5.95 | 7.11 | 2.66 | 7.07 | 20.62 | 5.00 |
| JAK | JAK_11 | 7.93 | 7.14 | 6.33 | 7.84 | 65.93 | 8.19 |
| JAK | JAK_12 | 7.62 | 8.02 | 6.27 | 8.33 | 78.71 | 8.25 |
| JAK | JAK_13 | 6.51 | 6.93 | 4.65 | 6.99 | 52.98 | 5.00 |
| JAK | JAK_14 | 5.98 | 6.63 | 3.04 | 6.76 | 22.49 | 5.00 |
| JAK | JAK_15 | 7.18 | 7.45 | 4.91 | 6.65 | 54.46 | 7.38 |
| JAK | JAK_16 | 5.90 | 7.59 | 2.60 | 7.35 | 27.59 | 5.69 |
| JAK | JAK_17 | 7.45 | 8.08 | 6.05 | 7.20 | 80.97 | 9.09 |
| JAK | JAK_18 | 7.67 | 8.33 | 6.41 | 7.48 | 70.54 | 7.09 |
| JAK | JAK_19 | 7.55 | 8.99 | 2.15 | 7.98 | 28.59 | 7.67 |
| JAK | JAK_20 | 6.07 | 6.49 | 2.96 | 7.21 | 31.39 | 5.00 |
| JAK | JAK_21 | 7.38 | 7.17 | 7.41 | 7.83 | 97.42 | 8.48 |
| JAK | JAK_22 | 5.67 | 6.47 | 3.22 | 7.17 | 35.79 | 5.00 |
| JAK | JAK_23 | 6.98 | 7.63 | 4.81 | 6.67 | 35.30 | 8.37 |
| JAK | JAK_24 | 7.99 | 7.78 | 5.60 | 8.51 | 58.22 | 7.79 |
| JAK | JAK_25 | 7.88 | 7.57 | 5.31 | 7.57 | 65.29 | 7.23 |
| JAK | JAK_26 | 7.99 | 8.49 | 6.56 | 7.59 | 43.31 | 8.10 |
| JAK | JAK_27 | 7.87 | 8.25 | 6.64 | 7.35 | 71.74 | 7.76 |
| JAK | JAK_28 | 6.01 | 6.58 | 2.74 | 7.20 | 16.96 | 4.56 |
| JAK | JAK_29 | 5.86 | 6.54 | 2.81 | 6.76 | 23.15 | 5.72 |
| JAK | JAK_30 | 5.46 | 7.04 | 2.41 | 7.07 | 30.55 | 6.25 |
| JAK | JAK_31 | 6.04 | 7.06 | 2.51 | 7.28 | 18.90 | 5.36 |
| JAK | JAK_32 | 7.06 | 7.31 | 1.09 | 7.11 | 27.16 | 5.92 |
| JAK | JAK_33 | 7.03 | 7.74 | 2.28 | 7.67 | 35.73 | 5.88 |
| JAK | JAK_34 | 5.87 | 7.54 | 1.52 | 7.28 | 30.58 | 6.56 |
| JAK | JAK_35 | 6.53 | 7.25 | 2.53 | 7.29 | 23.43 | 6.37 |
| JAK | JAK_36 | 5.78 | 6.61 | 3.30 | 7.34 | 31.81 | 4.00 |
| JAK | JAK_37 | 7.44 | 7.73 | 3.51 | 7.44 | 29.44 | 5.35 |
| JAK | JAK_38 | 7.29 | 7.35 | 3.68 | 7.61 | 24.80 | 5.88 |
| JAK | JAK_39 | 5.82 | 7.29 | 3.27 | 6.94 | 35.85 | 4.52 |
| JAK | JAK_40 | 6.65 | 7.20 | 2.81 | 7.49 | 23.37 | 6.79 |
| JAK | JAK_41 | 7.58 | 7.34 | 3.55 | 7.87 | 20.85 | 5.88 |
| JAK | JAK_42 | 6.58 | 7.18 | 3.31 | 7.61 | 20.67 | 5.18 |
| JAK | JAK_43 | 7.24 | 7.60 | 3.25 | 7.53 | 21.53 | 5.32 |
| JAK | JAK_44 | 7.56 | 7.12 | 3.76 | 7.67 | 39.85 | 5.04 |
| JAK | JAK_45 | 6.76 | 7.45 | 3.08 | 7.36 | 22.22 | 5.55 |
| JAK | JAK_46 | 6.40 | 7.11 | 3.15 | 7.65 | 18.31 | 4.82 |
| JAK | JAK_47 | 6.18 | 7.30 | 3.38 | 7.44 | 21.91 | 5.20 |
| JAK | JAK_48 | 6.12 | 7.20 | 3.41 | 7.36 | 34.22 | 6.37 |
| JAK | JAK_49 | 6.71 | 7.16 | 3.36 | 7.46 | 25.19 | 5.55 |
| JAK | JAK_50 | 6.87 | 7.26 | 3.53 | 7.43 | 19.50 | 6.00 |
| CDK2^7^ | CDK2_1 | 4.87 | 3.78 | 3.26 | 3.48 | 26.35 | 4.17 |
| CDK2 | CDK2_2 | 5.16 | 5.29 | 3.96 | 5.18 | 36.95 | 4.32 |
| CDK2 | CDK2_3 | 5.53 | 5.19 | 3.11 | 4.20 | 41.50 | 4.16 |
| CDK2 | CDK2_4 | 4.87 | 4.01 | 3.42 | 2.91 | 23.55 | 4.10 |
| CDK2 | CDK2_5 | 4.66 | 4.27 | 3.56 | 3.58 | 25.34 | 4.45 |
| CDK2 | CDK2_6 | 4.75 | 4.64 | 3.46 | 4.33 | 24.45 | 4.67 |
| CDK2 | CDK2_7 | 5.09 | 4.75 | 3.77 | 4.16 | 23.71 | 4.79 |
| CDK2 | CDK2_8 | 5.31 | 5.85 | 3.85 | 4.79 | 25.72 | 4.46 |
| CDK2 | CDK2_9 | 5.89 | 4.56 | 4.26 | 4.46 | 27.47 | 4.67 |
| CDK2 | CDK2_10 | 5.48 | 4.44 | 4.09 | 3.90 | 27.31 | 4.51 |
| CDK2 | CDK2_11 | 5.85 | 4.54 | 4.36 | 4.25 | 30.07 | 4.65 |
| CDK2 | CDK2_12 | 4.56 | 3.90 | 3.23 | 4.07 | 37.44 | 4.31 |
| CDK2 | CDK2_13 | 5.38 | 4.33 | 4.04 | 3.45 | 33.25 | 4.79 |
| CDK2 | CDK2_14 | 6.21 | 5.48 | 4.05 | 4.71 | 40.93 | 4.35 |
| CDK2 | CDK2_15 | 4.80 | 4.70 | 3.80 | 3.79 | 33.05 | 4.45 |
| CDK2 | CDK2_16 | 5.63 | 5.36 | 3.57 | 4.38 | 23.23 | 4.18 |
| CDK2 | CDK2_17 | 5.66 | 4.56 | 4.13 | 3.79 | 50.24 | 4.47 |
| CDK2 | CDK2_18 | 6.04 | 4.37 | 4.07 | 4.38 | 50.24 | 4.77 |
| CDK2 | CDK2_19 | 6.57 | 5.65 | 2.36 | 5.56 | 49.09 | 6.01 |
| CDK2 | CDK2_20 | 7.40 | 8.09 | -1.90 | 7.76 | 123.24 | 8.26 |
| CDK2 | CDK2_21 | 5.26 | 4.51 | 2.81 | 4.50 | 39.99 | 4.31 |
| CDK2 | CDK2_22 | 6.38 | 6.42 | 4.49 | 5.66 | 31.77 | 5.30 |
| CDK2 | CDK2_23 | 6.15 | 5.67 | 4.95 | 5.46 | 27.68 | 5.55 |
| CDK2 | CDK2_24 | 6.26 | 5.17 | 5.41 | 6.62 | 32.37 | 5.92 |
| CDK2 | CDK2_25 | 6.20 | 4.85 | 4.71 | 4.55 | 29.81 | 5.55 |
| CDK2 | CDK2_26 | 6.84 | 5.85 | 5.50 | 5.85 | 34.12 | 5.63 |
| CDK2 | CDK2_27 | 7.20 | 6.79 | 4.64 | 5.57 | 35.00 | 5.16 |
| CDK2 | CDK2_28 | 7.21 | 6.23 | 5.18 | 5.46 | 32.20 | 4.92 |
| CDK2 | CDK2_29 | 6.84 | 7.72 | -5.16 | 5.57 | 55.38 | 6.39 |
| CDK2 | CDK2_30 | 6.69 | 5.59 | 2.88 | 6.15 | 35.97 | 5.74 |
| CDK2 | CDK2_31 | 6.99 | 5.20 | 4.78 | 6.54 | 50.72 | 5.76 |
| CDK2 | CDK2_32 | 4.76 | 3.95 | 3.35 | 3.37 | 33.14 | 4.12 |
| CDK2 | CDK2_33 | 6.29 | 7.48 | -4.02 | 5.33 | 35.92 | 6.18 |
| CDK2 | CDK2_34 | 7.13 | 6.31 | -1.28 | 6.27 | 42.73 | 8.15 |
| CDK2 | CDK2_35 | 7.27 | 6.29 | -6.81 | 6.22 | 34.21 | 7.25 |
| CDK2 | CDK2_36 | 6.95 | 6.12 | -0.97 | 5.88 | 58.13 | 7.19 |
| CDK2 | CDK2_37 | 7.46 | 7.02 | -2.04 | 6.02 | 32.23 | 7.15 |
| CDK2 | CDK2_38 | 7.24 | 7.02 | -9.23 | 6.39 | 30.06 | 6.67 |
| CDK2 | CDK2_39 | 6.74 | 6.22 | 1.23 | 5.97 | 51.34 | 7.19 |
| CDK2 | CDK2_40 | 6.35 | 7.39 | -8.81 | 5.75 | 67.05 | 6.69 |
| CDK2 | CDK2_41 | 6.95 | 6.91 | -10.98 | 6.45 | 32.34 | 6.69 |
| CDK2 | CDK2_42 | 6.92 | 7.07 | -3.60 | 6.18 | 33.81 | 6.52 |
| CDK2 | CDK2_43 | 6.70 | 7.04 | -6.94 | 6.00 | 63.18 | 6.88 |
| CDK2 | CDK2_44 | 6.48 | 6.24 | 0.53 | 6.34 | 42.10 | 6.52 |
| CDK2 | CDK2_45 | 6.43 | 5.99 | 5.19 | 6.00 | 35.72 | 5.76 |
| CDK2 | CDK2_46 | 7.11 | 4.92 | 4.45 | 4.48 | 40.55 | 5.30 |
| CDK2 | CDK2_47 | 7.34 | 7.54 | -6.39 | 7.13 | 36.91 | 5.36 |
| CDK2 | CDK2_48 | 7.49 | 6.14 | -4.34 | 6.57 | 40.08 | 6.79 |
| CDK2 | CDK2_49 | 7.47 | 4.00 | 3.46 | 6.38 | 40.33 | 6.92 |
| CDK2 | CDK2_50 | 5.08 | 3.87 | 3.70 | 3.65 | 38.45 | 4.37 |
| CDK2 | CDK2_51 | 7.47 | 7.20 | -1.02 | 5.96 | 37.08 | 6.46 |
| CDK2 | CDK2_52 | 7.37 | 5.71 | 5.44 | 5.44 | 42.77 | 6.34 |
| CDK2 | CDK2_53 | 7.77 | 7.20 | -4.36 | 5.34 | 28.56 | 6.34 |
| CDK2 | CDK2_54 | 7.29 | 7.73 | -12.12 | 5.53 | 35.22 | 7.34 |
| CDK2 | CDK2_55 | 7.03 | 7.12 | -2.72 | 5.75 | 35.50 | 7.32 |
| CDK2 | CDK2_56 | 7.68 | 7.45 | -10.90 | 6.40 | 39.42 | 7.63 |
| CDK2 | CDK2_57 | 7.49 | 5.09 | 4.84 | 4.34 | 44.71 | 5.43 |
| CDK2 | CDK2_58 | 6.91 | 6.97 | -5.23 | 6.32 | 34.49 | 7.10 |
| CDK2 | CDK2_59 | 8.16 | 7.17 | -5.98 | 6.64 | 53.21 | 6.46 |
| CDK2 | CDK2_60 | 7.38 | 7.17 | -10.32 | 5.89 | 44.30 | 6.58 |
| CDK2 | CDK2_61 | 7.67 | 6.99 | -5.96 | 6.50 | 37.23 | 6.62 |
| CDK2 | CDK2_62 | 7.54 | 6.13 | 3.20 | 5.71 | 43.99 | 6.50 |
| CDK2 | CDK2_63 | 7.57 | 7.02 | -2.88 | 5.70 | 39.01 | 6.58 |
| CDK2 | CDK2_64 | 7.23 | 6.21 | 1.96 | 6.26 | 36.73 | 6.63 |
| CDK2 | CDK2_65 | 7.64 | 7.71 | -3.33 | 6.24 | 48.82 | 6.60 |
| CDK2 | CDK2_66 | 7.49 | 7.25 | -17.35 | 3.99 | 39.66 | 5.74 |
| CDK2 | CDK2_67 | 7.68 | 6.86 | -4.65 | 6.40 | 38.85 | 6.53 |
| CDK2 | CDK2_68 | 7.44 | 7.31 | -9.36 | 6.29 | 34.67 | 6.74 |
| CDK2 | CDK2_69 | 7.13 | 7.01 | -7.59 | 6.13 | 37.08 | 6.95 |
| CDK2 | CDK2_70 | 5.03 | 4.40 | 3.41 | 4.17 | 18.37 | 4.58 |
| CDK2 | CDK2_71 | 5.44 | 6.37 | 4.80 | 5.77 | 46.10 | 4.88 |
| CDK2 | CDK2_72 | 4.70 | 4.07 | 3.46 | 4.44 | 67.23 | 4.04 |
| THRB^7^ | THRB_1 | 5.79 | 5.79 | 2.14 | 6.13 | 92.14 | 5.39 |
| THRB | THRB_2 | 6.50 | 5.41 | 2.64 | 6.74 | 147.58 | 6.74 |
| THRB | THRB_3 | 6.80 | 5.86 | 2.48 | 6.50 | 111.40 | 6.25 |
| THRB | THRB_4 | 6.22 | 6.47 | 1.34 | 6.44 | 100.39 | 6.26 |
| THRB | THRB_5 | 6.04 | 5.02 | 3.91 | 6.28 | 86.97 | 4.82 |
| THRB | THRB_6 | 5.81 | 5.72 | -0.70 | 5.89 | 83.88 | 5.31 |
| THRB | THRB_7 | 5.74 | 5.92 | 2.90 | 6.77 | 94.28 | 5.30 |
| THRB | THRB_8 | 5.94 | 5.21 | 2.47 | 6.29 | 94.32 | 4.83 |
| THRB | THRB_9 | 6.00 | 5.18 | 4.50 | 5.83 | 95.32 | 5.91 |
| THRB | THRB_10 | 6.19 | 5.97 | -1.44 | 5.73 | 81.48 | 5.28 |
| THRB | THRB_11 | 5.72 | 5.33 | -1.63 | 6.02 | 75.53 | 3.73 |
| THRB | THRB_12 | 6.67 | 7.55 | 3.14 | 6.91 | 122.30 | 8.39 |
| THRB | THRB_13 | 6.16 | 4.55 | 3.33 | 5.20 | 73.34 | 4.95 |
| THRB | THRB_14 | 6.64 | 5.67 | 3.31 | 7.56 | 135.41 | 7.22 |
| THRB | THRB_15 | 6.92 | 5.81 | 3.18 | 6.46 | 130.92 | 6.58 |
| THRB | THRB_16 | 6.04 | 6.01 | 3.50 | 6.84 | 144.36 | 6.56 |
| THRB | THRB_17 | 6.20 | 5.75 | 3.20 | 8.01 | 124.61 | 6.36 |
| THRB | THRB_18 | 6.89 | 6.31 | 3.36 | 7.54 | 137.18 | 6.30 |
| THRB | THRB_19 | 6.49 | 5.86 | 3.05 | 6.76 | 94.17 | 5.55 |
| THRB | THRB_20 | 6.17 | 5.38 | 3.45 | 6.91 | 76.15 | 5.39 |
| THRB | THRB_21 | 6.27 | 5.33 | 2.75 | 6.39 | 82.75 | 5.83 |
| THRB | THRB_22 | 5.71 | 5.42 | 3.74 | 5.56 | 49.54 | 5.11 |
| THRB | THRB_23 | 5.49 | 6.23 | 2.40 | 6.18 | 96.66 | 4.12 |
| THRB | THRB_24 | 6.04 | 5.56 | 3.95 | 6.38 | 107.45 | 4.46 |
| BRD4^8^ | BRD4_1 | 6.62 | 5.75 | 4.97 | 5.55 | 99.91 | 6.14 |
| BRD4 | BRD4_2 | 6.45 | 5.65 | 4.93 | 5.38 | 85.55 | 5.42 |
| BRD4 | BRD4_3 | 6.73 | 5.96 | 4.87 | 5.62 | 97.23 | 5.61 |
| BRD4 | BRD4_4 | 7.13 | 6.67 | 4.77 | 6.00 | 98.81 | 6.48 |
| BRD4 | BRD4_5 | 6.84 | 5.78 | 4.26 | 5.77 | 82.56 | 6.77 |
| BRD4 | BRD4_6 | 6.35 | 5.97 | 4.15 | 5.34 | 29.80 | 6.11 |
| BRD4 | BRD4_7 | 7.29 | 5.81 | 4.62 | 6.18 | 81.75 | 6.62 |
| BRD4 | BRD4_8 | 6.91 | 5.61 | 4.47 | 5.64 | 79.51 | 6.26 |
| BRD4 | BRD4_9 | 6.88 | 5.91 | 4.45 | 5.69 | 78.62 | 5.81 |
| BRD4 | BRD4_10 | 6.84 | 5.98 | 4.62 | 5.68 | 84.78 | 6.14 |
| BRD4 | BRD4_11 | 6.65 | 6.26 | 4.59 | 5.43 | 77.89 | 7.30 |
| BRD4 | BRD4_12 | 7.21 | 5.87 | 4.73 | 5.58 | 82.18 | 7.04 |
| BRD4 | BRD4_13 | 6.84 | 6.55 | 4.75 | 5.92 | 75.61 | 5.41 |
| BRD4 | BRD4_14 | 6.66 | 6.08 | 4.73 | 5.74 | 80.61 | 6.76 |
| BRD4 | BRD4_15 | 7.19 | 6.13 | 4.81 | 5.85 | 84.03 | 7.04 |
| BRD4 | BRD4_16 | 6.89 | 5.84 | 4.84 | 5.75 | 90.83 | 5.04 |
| BRD4 | BRD4_17 | 7.11 | 6.35 | 4.87 | 5.91 | 89.16 | 5.77 |
| BRD4 | BRD4_18 | 7.09 | 5.67 | 4.87 | 5.90 | 94.34 | 6.09 |
| BRD4 | BRD4_19 | 7.15 | 5.90 | 4.73 | 6.01 | 81.67 | 7.03 |
| BRD4 | BRD4_20 | 7.11 | 5.98 | 4.48 | 5.67 | 74.23 | 6.52 |
| BRD4 | BRD4_21 | 6.96 | 6.17 | 4.38 | 5.78 | 39.87 | 7.05 |
| BRD4 | BRD4_22 | 6.83 | 5.56 | 4.15 | 5.60 | 44.45 | 5.46 |
| BRD4 | BRD4_23 | 6.70 | 5.53 | 4.27 | 5.56 | 41.95 | 5.80 |
| BRD4 | BRD4_24 | 6.74 | 6.09 | 4.26 | 5.62 | 34.27 | 6.68 |
| BRD4 | BRD4_25 | 7.40 | 5.57 | 5.21 | 5.72 | 95.34 | 5.58 |
| BRD4 | BRD4_26 | 7.08 | 5.89 | 5.11 | 5.81 | 82.50 | 6.18 |
| BRD4 | BRD4_27 | 7.44 | 6.39 | 4.84 | 5.82 | 95.13 | 6.24 |
| BRD4 | BRD4_28 | 7.01 | 6.48 | 4.70 | 5.65 | 98.89 | 6.56 |
| BRD4 | BRD4_29 | 7.67 | 6.81 | 4.58 | 6.11 | 100.49 | 6.94 |
| BRD4 | BRD4_30 | 7.55 | 6.88 | 4.94 | 6.18 | 89.82 | 6.29 |
| BRD4 | BRD4_31 | 7.73 | 6.12 | 4.59 | 6.13 | 101.63 | 6.48 |
| BRD4 | BRD4_32 | 7.11 | 5.71 | 4.38 | 6.16 | 95.14 | 6.65 |
| BRD4 | BRD4_33 | 7.74 | 6.77 | 4.31 | 6.08 | 103.09 | 5.65 |
| BRD4 | BRD4_34 | 6.94 | 6.22 | 4.98 | 5.92 | 91.90 | 7.02 |
| BRD4 | BRD4_35 | 7.32 | 6.31 | 4.78 | 6.29 | 92.34 | 6.66 |
| BRD4 | BRD4_36 | 7.49 | 6.18 | 4.40 | 5.86 | 77.74 | 7.0 |
| BRD4 | BRD4_37 | 7.50 | 5.92 | 4.57 | 6.06 | 91.90 | 6.40 |
| BRD4 | BRD4_38 | 7.68 | 6.45 | 4.17 | 5.97 | 79.36 | 6.77 |
| BRD4 | BRD4_39 | 7.51 | 6.19 | 4.13 | 6.27 | 80.43 | 6.14 |
| BRD4 | BRD4_40 | 7.11 | 5.54 | 4.37 | 6.43 | 76.16 | 6.47 |
| BRD4 | BRD4_41 | 7.38 | 6.53 | 5.24 | 5.95 | 85.17 | 7.45 |
| BRD4 | BRD4_42 | 7.43 | 6.41 | 4.86 | 5.86 | 98.43 | 7.43 |
| BRD4 | BRD4_43 | 7.48 | 6.13 | 4.74 | 5.88 | 90.67 | 6.05 |
| BRD4 | BRD4_44 | 7.38 | 6.51 | 4.87 | 5.97 | 80.46 | 7.05 |
| BRD4 | BRD4_45 | 7.36 | 7.18 | 4.89 | 5.77 | 81.77 | 7.89 |
| BRD4 | BRD4_46 | 7.87 | 7.19 | 4.79 | 5.90 | 75.68 | 7.77 |
| BRD4 | BRD4_47 | 7.48 | 6.29 | 5.23 | 5.79 | 76.35 | 7.75 |
| BRD4 | BRD4_48 | 7.56 | 6.32 | 5.01 | 6.07 | 82.57 | 6.45 |
| BRD4 | BRD4_49 | 7.61 | 6.19 | 5.03 | 6.05 | 65.80 | 7.26 |
| BRD4 | BRD4_50 | 6.86 | 6.51 | 5.10 | 5.87 | 71.45 | 6.78 |
| BRD4 | BRD4_51 | 7.93 | 6.82 | 4.85 | 6.23 | 87.10 | 7.35 |
| BRD4 | BRD4_52 | 8.20 | 6.79 | 4.50 | 6.47 | 87.38 | 6.97 |
| BRD4 | BRD4_53 | 8.25 | 6.83 | 4.73 | 5.90 | 83.96 | 7.84 |
| BRD4 | BRD4_54 | 8.08 | 6.33 | 4.97 | 6.07 | 86.81 | 6.17 |
| BRD4 | BRD4_55 | 8.50 | 6.81 | 5.07 | 6.09 | 90.51 | 7.42 |
| BRD4 | BRD4_56 | 8.29 | 6.97 | 4.87 | 6.23 | 80.13 | 7.18 |
| BRD4 | BRD4_57 | 8.26 | 6.93 | 4.86 | 6.17 | 84.27 | 7.18 |
| BRD4 | BRD4_58 | 8.41 | 7.03 | 4.67 | 6.15 | 80.48 | 7.20 |
| BRD4 | BRD4_59 | 7.34 | 5.86 | 5.19 | 6.34 | 93.08 | 6.68 |
| TYK2^7^ | TYK2_1 | 6.04 | 9.28 | 0.27 | 6.24 | 26.43 | 6.63 |
| TYK2 | TYK2_2 | 5.14 | 6.87 | 0.44 | 5.12 | 31.99 | 5.50 |
| TYK2 | TYK2_3 | 5.48 | 9.79 | 0.97 | 5.75 | 64.50 | 5.95 |
| TYK2 | TYK2_4 | 5.56 | 7.03 | 0.31 | 5.81 | 73.42 | 5.61 |
| TYK2 | TYK2_5 | 5.71 | 7.30 | 0.17 | 6.39 | 65.75 | 6.63 |
| TYK2 | TYK2_6 | 5.68 | 9.63 | 0.81 | 5.94 | 30.09 | 6.26 |
| TYK2 | TYK2_7 | 5.88 | 9.32 | 0.30 | 6.65 | 21.80 | 5.55 |
| TYK2 | TYK2_8 | 6.22 | 9.22 | 0.78 | 6.53 | 25.92 | 5.72 |
| TYK2 | TYK2_9 | 5.71 | 9.10 | 0.23 | 6.09 | 59.60 | 6.10 |
| TYK2 | TYK2_10 | 6.14 | 8.90 | 1.03 | 6.37 | 66.96 | 6.04 |
| TYK2 | TYK2_11 | 6.10 | 8.93 | 0.85 | 6.70 | 70.16 | 5.55 |
| TYK2 | TYK2_12 | 5.62 | 9.63 | 0.19 | 7.28 | 40.98 | 5.92 |
| TYK2 | TYK2_13 | 5.85 | 9.21 | 0.94 | 6.26 | 84.47 | 6.42 |
| TYK2 | TYK2_14 | 5.64 | 9.03 | -1.41 | 6.30 | 80.97 | 5.49 |
| TYK2 | TYK2_15 | 6.27 | 7.27 | 0.05 | 6.70 | 29.10 | 5.95 |
| TYK2 | TYK2_16 | 6.15 | 8.91 | 0.13 | 6.43 | 29.67 | 6.03 |
| TYK2 | TYK2_17 | 6.49 | 9.70 | 1.26 | 8.35 | 37.94 | 5.82 |
| TYK2 | TYK2_18 | 6.75 | 8.79 | 0.07 | 7.75 | 38.48 | 7.01 |
| TYK2 | TYK2_19 | 6.35 | 8.43 | 1.20 | 7.73 | 48.64 | 6.85 |
| TYK2 | TYK2_20 | 7.14 | 8.86 | 1.14 | 7.95 | 26.91 | 7.53 |
| TYK2 | TYK2_21 | 6.86 | 9.62 | 0.30 | 8.21 | 46.30 | 5.92 |
| TYK2 | TYK2_22 | 6.98 | 9.40 | 0.34 | 8.30 | 34.93 | 7.85 |
| TYK2 | TYK2_23 | 7.07 | 9.27 | 0.35 | 8.27 | 47.99 | 8.30 |
| TYK2 | TYK2_24 | 7.05 | 9.27 | 1.25 | 7.84 | 39.65 | 8.0 |
| TYK2 | TYK2_25 | 6.74 | 9.13 | 0.46 | 7.80 | 39.96 | 7.72 |
| TYK2 | TYK2_26 | 6.22 | 8.22 | 0.17 | 7.07 | 43.28 | 7.0 |
| TYK2 | TYK2_27 | 6.49 | 8.71 | 0.29 | 8.62 | 39.85 | 6.61 |
| TYK2 | TYK2_28 | 6.13 | 8.93 | 1.69 | 7.57 | 41.56 | 7.19 |
| TYK2 | TYK2_29 | 6.41 | 9.33 | 0.35 | 7.79 | 44.89 | 6.07 |
| TYK2 | TYK2_30 | 6.91 | 9.13 | 0.20 | 7.81 | 39.34 | 7.02 |
| TYK2 | TYK2_31 | 7.20 | 8.91 | 0.29 | 7.82 | 36.45 | 8.31 |
| TYK2 | TYK2_32 | 6.92 | 9.26 | 0.32 | 8.01 | 34.00 | 7.13 |
| TYK2 | TYK2_33 | 7.13 | 7.74 | 0.52 | 6.63 | 51.62 | 6.61 |
| TYK2 | TYK2_34 | 7.06 | 8.47 | 0.18 | 7.63 | 43.75 | 5.69 |
| TYK2 | TYK2_35 | 5.96 | 8.29 | -0.01 | 7.25 | 34.82 | 6.60 |
| TYK2 | TYK2_36 | 6.64 | 9.15 | 0.03 | 6.08 | 25.93 | 5.92 |
| TYK2 | TYK2_37 | 6.73 | 9.06 | 0.10 | 7.28 | 56.39 | 5.82 |
| TYK2 | TYK2_38 | 6.75 | 8.24 | 0.19 | 7.24 | 35.06 | 7.74 |
| TYK2 | TYK2_39 | 6.34 | 8.50 | 0.31 | 6.95 | 45.51 | 6.76 |
| TYK2 | TYK2_40 | 6.86 | 9.32 | 1.03 | 7.88 | 48.07 | 9.30 |
| TYK2 | TYK2_41 | 7.20 | 8.83 | 1.74 | 7.78 | 35.45 | 8.31 |
| TYK2 | TYK2_42 | 6.68 | 8.86 | 0.31 | 8.18 | 40.58 | 9.0 |
| TYK2 | TYK2_43 | 6.41 | 8.92 | 0.24 | 8.47 | 37.16 | 8.52 |
| TYK2 | TYK2_44 | 6.69 | 7.10 | 1.61 | 8.18 | 45.92 | 7.86 |
| TYK2 | TYK2_45 | 6.46 | 6.88 | 1.65 | 8.18 | 39.64 | 8.50 |
| TYK2 | TYK2_46 | 6.54 | 7.83 | 1.42 | 8.20 | 26.97 | 8.10 |
| TYK2 | TYK2_47 | 6.75 | 9.11 | 1.23 | 8.22 | 29.07 | 8.26 |
| TYK2 | TYK2_48 | 6.36 | 8.13 | 1.48 | 8.72 | 35.41 | 8.11 |
| TYK2 | TYK2_49 | 6.60 | 9.74 | 1.78 | 8.52 | 36.17 | 7.66 |
| TYK2 | TYK2_50 | 6.80 | 9.28 | 0.10 | 7.95 | 31.44 | 7.98 |
| TYK2 | TYK2_51 | 6.49 | 8.94 | 1.39 | 8.33 | 51.88 | 8.35 |
| TYK2 | TYK2_52 | 6.48 | 7.56 | 1.49 | 8.64 | 42.30 | 8.48 |
| TYK2 | TYK2_53 | 6.81 | 8.37 | 1.56 | 8.32 | 39.90 | 7.21 |
| TYK2 | TYK2_54 | 7.10 | 8.41 | 1.68 | 8.26 | 30.26 | 7.45 |
| TYK2 | TYK2_55 | 6.68 | 7.23 | 1.70 | 8.58 | 44.59 | 8.10 |
| TYK2 | TYK2_56 | 6.51 | 9.35 | 1.74 | 8.32 | 42.22 | 7.52 |
| TYK2 | TYK2_57 | 7.26 | 8.15 | 1.48 | 8.59 | 42.49 | 8.74 |
| TYK2 | TYK2_58 | 7.49 | 8.27 | 1.70 | 8.45 | 30.02 | 8.02 |
| TYK2 | TYK2_59 | 7.18 | 6.42 | 0.05 | 8.44 | 43.94 | 8.46 |
| TYK2 | TYK2_60 | 6.71 | 9.71 | 0.09 | 8.31 | 43.75 | 7.96 |
| TYK2 | TYK2_61 | 7.08 | 9.21 | 1.35 | 7.99 | 40.23 | 8.60 |
| TYK2 | TYK2_62 | 6.65 | 9.21 | 1.35 | 8.00 | 39.52 | 8.14 |
| TYK2 | TYK2_63 | 6.65 | 9.21 | 1.35 | 8.00 | 39.52 | 7.94 |
| TYK2 | TYK2_64 | 6.88 | 9.07 | 1.31 | 7.99 | 36.75 | 7.65 |
| TYK2 | TYK2_65 | 6.66 | 9.06 | 1.78 | 8.05 | 36.25 | 8.29 |
| TYK2 | TYK2_66 | 6.68 | 9.02 | 1.69 | 7.80 | 32.45 | 8.07 |
| TYK2 | TYK2_67 | 6.53 | 7.78 | 1.38 | 8.38 | 34.05 | 7.83 |
| TYK2 | TYK2_68 | 7.49 | 8.23 | 1.45 | 8.05 | 35.92 | 8.04 |
| TYK2 | TYK2_69 | 6.75 | 8.89 | 1.34 | 8.14 | 42.80 | 7.95 |
| TYK2 | TYK2_70 | 6.35 | 9.06 | 1.61 | 8.19 | 36.75 | 8.49 |
| TYK2 | TYK2_71 | 7.08 | 8.30 | 0.74 | 8.88 | 41.28 | 8.88 |
| TYK2 | TYK2_72 | 6.55 | 8.52 | 1.20 | 9.01 | 38.22 | 7.82 |
| TYK2 | TYK2_73 | 7.53 | 8.98 | 1.30 | 8.70 | 40.63 | 8.85 |
| TYK2 | TYK2_74 | 6.63 | 8.98 | 1.21 | 8.72 | 33.87 | 8.79 |
| TYK2 | TYK2_75 | 6.74 | 8.60 | 1.21 | 7.87 | 39.77 | 8.79 |
| MTH1^9^ | MTH1_1 | 4.86 | 5.31 | 1.69 | 5.05 | 70.17 | 5.00 |
| MTH1 | MTH1_2 | 6.04 | 7.28 | 4.84 | 6.35 | 45.73 | 5.60 |
| MTH1 | MTH1_3 | 5.46 | 6.01 | 4.12 | 5.95 | 58.04 | 5.43 |
| MTH1 | MTH1_4 | 5.56 | 7.10 | 2.99 | 6.92 | 49.42 | 5.72 |
| MTH1 | MTH1_5 | 5.59 | 6.93 | 1.75 | 6.78 | 36.41 | 6.69 |
| MTH1 | MTH1_6 | 5.67 | 6.76 | 0.29 | 7.15 | 80.74 | 7.92 |
| MTH1 | MTH1_7 | 5.77 | 7.01 | -0.41 | 6.92 | 82.45 | 7.69 |
| MTH1 | MTH1_8 | 6.35 | 6.90 | 4.47 | 6.96 | 42.63 | 5.56 |
| MTH1 | MTH1_9 | 5.40 | 7.21 | 0.46 | 6.68 | 91.78 | 5.23 |
| MTH1 | MTH1_10 | 6.40 | 7.35 | 4.93 | 6.92 | 69.83 | 8.0 |
| MTH1 | MTH1_11 | 5.61 | 7.39 | 3.16 | 6.75 | 55.61 | 5.0 |
| MTH1 | MTH1_12 | 5.77 | 7.06 | 3.13 | 7.12 | 54.21 | 5.25 |
| MTH1 | MTH1_13 | 6.22 | 8.19 | -2.85 | 7.39 | 65.35 | 5.0 |
| MTH1 | MTH1_14 | 5.61 | 7.57 | 1.10 | 6.95 | 72.76 | 5.85 |
| MTH1 | MTH1_15 | 5.46 | 7.32 | 3.91 | 6.69 | 40.05 | 7.39 |
| MTH1 | MTH1_16 | 5.80 | 7.27 | 5.67 | 7.37 | 58.22 | 8.30 |
| MTH1 | MTH1_17 | 5.50 | 6.32 | 3.62 | 6.89 | 73.56 | 6.69 |
| MTH1 | MTH1_18 | 5.63 | 8.00 | 4.73 | 6.83 | 52.98 | 6.15 |
| MTH1 | MTH1_19 | 5.65 | 7.11 | 4.80 | 6.36 | 39.11 | 6.52 |
| MTH1 | MTH1_20 | 6.28 | 8.23 | -0.02 | 7.59 | 75.43 | 10.0 |
| MTH1 | MTH1_21 | 6.63 | 7.83 | -0.01 | 7.53 | 70.92 | 10.0 |
| MTH1 | MTH1_22 | 6.14 | 7.88 | 0.09 | 7.59 | 70.52 | 10.0 |
| MTH1 | MTH1_23 | 5.94 | 8.01 | -2.85 | 7.49 | 47.49 | 10.0 |
| MTH1 | MTH1_24 | 5.64 | 6.91 | 3.22 | 6.77 | 47.15 | 5.52 |
| MTH1 | MTH1_25 | 6.10 | 7.98 | 2.07 | 7.21 | 47.22 | 9.0 |
| MTH1 | MTH1_26 | 6.40 | 7.94 | 5.45 | 6.99 | 49.05 | 7.52 |
| MTH1 | MTH1_27 | 5.81 | 7.52 | 5.38 | 7.16 | 57.89 | 8.69 |
| MTH1 | MTH1_28 | 6.23 | 8.81 | 5.89 | 7.04 | 53.29 | 8.30 |
| MTH1 | MTH1_29 | 5.64 | 8.11 | 2.66 | 7.17 | 62.34 | 10.0 |
| MTH1 | MTH1_30 | 5.42 | 7.50 | 2.08 | 7.30 | 65.13 | 9.09 |
| MTH1 | MTH1_31 | 6.13 | 8.48 | 2.64 | 7.63 | 77.74 | 10.0 |
| MTH1 | MTH1_32 | 6.08 | 8.57 | 4.49 | 7.25 | 64.94 | 10.0 |
| ULK2^10^ | ULK2_1 | 4.63 | 4.69 | 3.82 | 4.30 | 45.97 | 6.88 |
| ULK2 | ULK2_2 | 6.05 | 7.18 | 5.98 | 6.74 | 103.22 | 6.63 |
| ULK2 | ULK2_3 | 6.27 | 7.38 | 7.18 | 7.78 | 114.79 | 7.67 |
| ULK2 | ULK2_4 | 6.76 | 7.70 | 6.78 | 7.52 | 111.26 | 7.37 |
| ULK2 | ULK2_5 | 6.66 | 7.79 | 6.68 | 7.08 | 120.12 | 7.85 |
| ULK2 | ULK2_6 | 6.99 | 7.31 | 4.84 | 7.76 | 111.38 | 8.15 |
| ULK2 | ULK2_7 | 6.63 | 7.99 | 6.60 | 7.84 | 97.34 | 8.30 |
| ULK2 | ULK2_8 | 6.62 | 7.09 | 6.96 | 7.67 | 108.94 | 7.63 |
| ULK2 | ULK2_9 | 6.95 | 7.13 | 6.75 | 7.81 | 114.10 | 7.85 |
| ULK2 | ULK2_10 | 6.85 | 8.13 | 6.37 | 7.62 | 113.14 | 8.22 |
| ULK2 | ULK2_11 | 6.75 | 7.37 | 7.04 | 7.97 | 112.88 | 7.53 |
| ULK2 | ULK2_12 | 7.19 | 7.71 | 7.64 | 7.81 | 116.07 | 7.76 |
| ULK2 | ULK2_13 | 6.50 | 7.27 | 6.13 | 7.64 | 110.47 | 7.46 |
| ULK2 | ULK2_14 | 6.94 | 8.15 | 6.43 | 8.10 | 114.88 | 7.50 |
| ULK2 | ULK2_15 | 6.69 | 8.07 | 6.77 | 7.39 | 116.15 | 7.72 |
| ULK2 | ULK2_16 | 6.57 | 7.70 | 6.75 | 7.50 | 123.46 | 8.30 |
| ULK2 | ULK2_17 | 6.53 | 8.16 | 6.12 | 7.72 | 108.75 | 7.79 |
| ULK2 | ULK2_18 | 6.70 | 7.82 | 6.20 | 7.59 | 88.29 | 7.88 |
| ULK2 | ULK2_19 | 6.97 | 8.28 | 6.48 | 7.62 | 117.95 | 8.52 |
| ULK2 | ULK2_20 | 6.31 | 7.42 | 6.03 | 7.11 | 100.09 | 7.39 |
| ULK2 | ULK2_21 | 6.67 | 8.09 | 6.34 | 7.48 | 116.54 | 7.85 |
| ULK2 | ULK2_22 | 6.29 | 7.35 | 6.91 | 6.73 | 103.48 | 7.30 |
| ULK2 | ULK2_23 | 6.07 | 6.90 | 6.49 | 7.04 | 106.91 | 7.69 |
| ULK2 | ULK2_24 | 6.39 | 7.16 | 6.28 | 7.35 | 96.33 | 7.88 |
| ULK2 | ULK2_25 | 6.95 | 7.18 | 6.69 | 8.26 | 109.27 | 7.52 |
| ULK2 | ULK2_26 | 6.25 | 7.88 | 6.23 | 6.93 | 112.85 | 7.88 |
| FKBP51^11^ | FKBP51_1 | 7.83 | 6.02 | 6.19 | 6.97 | 85.60 | 8.39 |
| FKBP51 | FKBP51_2 | 8.17 | 6.51 | 5.99 | 7.58 | 88.72 | 8.22 |
| FKBP51 | FKBP51_3 | 7.74 | 6.60 | 6.20 | 6.92 | 86.02 | 7.36 |
| FKBP51 | FKBP51_4 | 7.56 | 5.97 | 6.50 | 6.74 | 83.21 | 5.92 |
| FKBP51 | FKBP51_5 | 7.64 | 5.79 | 6.80 | 7.14 | 80.76 | 5.61 |
| FKBP51 | FKBP51_6 | 7.35 | 6.00 | 6.43 | 7.15 | 72.30 | 5.10 |
| FKBP51 | FKBP51_7 | 7.57 | 6.20 | 7.31 | 7.39 | 88.49 | 5.58 |
| FKBP51 | FKBP51_8 | 7.73 | 5.69 | 6.78 | 7.27 | 80.42 | 5.69 |
| FKBP51 | FKBP51_9 | 7.56 | 6.49 | 7.14 | 7.39 | 86.17 | 5.72 |
| FKBP51 | FKBP51_10 | 6.89 | 5.99 | 6.55 | 7.27 | 76.07 | 5.58 |
| FKBP51 | FKBP51_11 | 7.66 | 5.88 | 6.77 | 7.29 | 84.54 | 6.09 |
| FKBP51 | FKBP51_12 | 7.80 | 5.79 | 6.81 | 7.34 | 81.78 | 6.30 |
| FKBP51 | FKBP51_13 | 6.88 | 6.00 | 6.59 | 7.10 | 79.81 | 4.55 |
| FKBP51 | FKBP51_14 | 7.38 | 5.99 | 6.56 | 7.19 | 76.76 | 5.50 |
| FKBP51 | FKBP51_15 | 7.40 | 6.38 | 6.55 | 7.05 | 83.89 | 5.88 |
| FKBP51 | FKBP51_16 | 7.49 | 6.62 | 6.85 | 7.22 | 77.58 | 6.63 |
| FKBP51 | FKBP51_17 | 7.47 | 6.39 | 6.98 | 7.52 | 78.05 | 6.48 |
| FKBP51 | FKBP51_18 | 7.47 | 6.65 | 6.96 | 7.19 | 83.13 | 6.18 |
| FKBP51 | FKBP51_19 | 7.90 | 6.83 | 6.47 | 7.51 | 74.73 | 5.08 |
| FKBP51 | FKBP51_20 | 7.78 | 6.29 | 6.75 | 7.30 | 74.44 | 6.55 |
| MCL-1^7,12^ | MCL-1_2_1 | 6.98 | 7.03 | 5.77 | 5.48 | 57.99 | 5.82 |
| MCL-1 | MCL-1_2_2 | 6.56 | 5.40 | 5.60 | 5.53 | 46.44 | 5.58 |
| MCL-1 | MCL-1_2_3 | 6.91 | 6.46 | 5.99 | 5.51 | 57.56 | 4.15 |
| MCL-1 | MCL-1_2_4 | 6.81 | 6.64 | 6.08 | 5.53 | 56.75 | 5.21 |
| MCL-1 | MCL-1_2_5 | 6.81 | 6.86 | 5.84 | 5.72 | 62.60 | 5.56 |
| MCL-1 | MCL-1_2_6 | 7.23 | 7.21 | 5.56 | 6.09 | 59.88 | 5.76 |
| MCL-1 | MCL-1_2_7 | 7.78 | 7.06 | 5.97 | 6.34 | 56.82 | 7.06 |
| MCL-1 | MCL-1_2_8 | 7.70 | 7.65 | 5.75 | 6.19 | 57.66 | 6.69 |
| MCL-1 | MCL-1_2_9 | 7.60 | 6.71 | 5.98 | 5.91 | 55.83 | 6.48 |
| MCL-1 | MCL-1_2_10 | 7.39 | 6.99 | 5.81 | 5.77 | 57.58 | 6.24 |
| MCL-1 | MCL-1_2_11 | 7.39 | 6.18 | 5.70 | 5.94 | 53.40 | 6.07 |
| MCL-1 | MCL-1_2_12 | 7.57 | 6.67 | 5.69 | 5.97 | 54.30 | 6.11 |
| MCL-1 | MCL-1_2_13 | 7.96 | 6.73 | 5.65 | 6.72 | 58.32 | 7.13 |
| MCL-1 | MCL-1_2_14 | 7.90 | 6.48 | 5.27 | 6.73 | 51.86 | 7.02 |
| MCL-1 | MCL-1_2_15 | 7.95 | 6.45 | 5.66 | 6.91 | 50.48 | 6.53 |
| MCL-1 | MCL-1_2_16 | 8.11 | 6.24 | 5.03 | 6.63 | 54.07 | 7.13 |
| MCL-1 | MCL-1_1_1 | 5.03 | 4.90 | 3.33 | 3.19 | 25.48 | 3.66 |
| MCL-1 | MCL-1_1_2 | 4.87 | 5.63 | 3.26 | 3.68 | 44.76 | 4.04 |
| MCL-1 | MCL-1_1_3 | 4.49 | 3.81 | 2.55 | 2.98 | 42.60 | 3.79 |
| MCL-1 | MCL-1_1_4 | 4.99 | 4.18 | 2.06 | 4.17 | 32.86 | 4.09 |
| MCL-1 | MCL-1_1_5 | 5.35 | 5.81 | 3.69 | 4.20 | 17.35 | 3.86 |
| MCL-1 | MCL-1_1_6 | 4.54 | 4.30 | 2.74 | 4.18 | 40.27 | 3.98 |
| MCL-1 | MCL-1_1_7 | 5.02 | 4.10 | 3.60 | 3.92 | 44.31 | 4.22 |
| MCL-1 | MCL-1_1_8 | 5.06 | 4.51 | 3.01 | 4.18 | 32.05 | 4.06 |
| MCL-1 | MCL-1_1_9 | 4.67 | 3.95 | 3.31 | 3.31 | 44.88 | 3.88 |
| MCL-1 | MCL-1_1_10 | 5.76 | 6.24 | 4.15 | 7.02 | 40.79 | 4.90 |
| MCL-1 | MCL-1_1_11 | 6.22 | 6.42 | 4.65 | 6.35 | 60.37 | 5.01 |
| MCL-1 | MCL-1_1_12 | 6.06 | 5.81 | 5.58 | 6.66 | 101.50 | 6.42 |
| MCL-1 | MCL-1_1_13 | 5.45 | 6.33 | 5.82 | 5.99 | 61.90 | 5.02 |
| MCL-1 | MCL-1_1_14 | 5.91 | 6.89 | 4.76 | 6.49 | 58.75 | 4.83 |
| MCL-1 | MCL-1_1_15 | 5.81 | 5.83 | 5.37 | 6.31 | 103.76 | 6.0 |
| MCL-1 | MCL-1_1_16 | 5.37 | 5.41 | 4.00 | 4.94 | 58.16 | 4.45 |
| MCL-1 | MCL-1_1_17 | 5.69 | 5.82 | 4.71 | 6.19 | 64.90 | 4.82 |
| MCL-1 | MCL-1_1_18 | 5.78 | 6.11 | 5.09 | 5.94 | 77.70 | 5.05 |
| MCL-1 | MCL-1_1_19 | 4.98 | 4.35 | 2.37 | 4.46 | 23.60 | 4.65 |
| MCL-1 | MCL-1_1_20 | 5.95 | 5.86 | 4.65 | 5.47 | 88.33 | 5.72 |
| MCL-1 | MCL-1_1_21 | 6.11 | 5.29 | 5.21 | 6.57 | 91.95 | 5.76 |
| MCL-1 | MCL-1_1_22 | 5.63 | 5.37 | 4.83 | 5.61 | 93.12 | 4.79 |
| MCL-1 | MCL-1_1_23 | 5.70 | 5.78 | 4.67 | 5.63 | 72.66 | 5.00 |
| MCL-1 | MCL-1_1_24 | 6.02 | 5.57 | 5.68 | 5.81 | 90.73 | 5.00 |
| MCL-1 | MCL-1_1_25 | 6.32 | 6.04 | 4.96 | 6.37 | 95.66 | 6.42 |
| MCL-1 | MCL-1_1_26 | 6.40 | 5.55 | 5.03 | 6.06 | 84.54 | 5.95 |
| MCL-1 | MCL-1_1_27 | 6.65 | 6.05 | 5.69 | 6.87 | 99.56 | 6.52 |
| MCL-1 | MCL-1_1_28 | 5.81 | 5.79 | 3.79 | 6.92 | 92.33 | 5.11 |
| MCL-1 | MCL-1_1_29 | 5.92 | 5.27 | 1.12 | 7.02 | 74.92 | 5.11 |
| MCL-1 | MCL-1_1_30 | 5.02 | 5.05 | 0.69 | 3.75 | 51.60 | 4.22 |
| MCL-1 | MCL-1_1_31 | 6.25 | 5.71 | 0.38 | 6.64 | 59.03 | 5.28 |
| MCL-1 | MCL-1_1_32 | 5.89 | 5.62 | 5.35 | 6.70 | 119.53 | 6.52 |
| MCL-1 | MCL-1_1_33 | 5.97 | 5.96 | 5.49 | 6.66 | 111.51 | 5.12 |
| MCL-1 | MCL-1_1_34 | 6.59 | 5.64 | 5.74 | 6.29 | 105.04 | 6.31 |
| MCL-1 | MCL-1_1_35 | 6.28 | 5.84 | 5.59 | 6.19 | 63.56 | 6.52 |
| MCL-1 | MCL-1_1_36 | 5.96 | 5.25 | 5.33 | 6.25 | 66.02 | 5.53 |
| MCL-1 | MCL-1_1_37 | 6.15 | 4.98 | 5.19 | 5.72 | 33.20 | 4.20 |
| MCL-1 | MCL-1_1_38 | 6.24 | 6.06 | 5.20 | 5.77 | 76.57 | 4.85 |
| MCL-1 | MCL-1_1_39 | 6.32 | 6.12 | 5.43 | 6.72 | 93.14 | 6.09 |
| MCL-1 | MCL-1_1_40 | 5.01 | 4.06 | 2.76 | 3.99 | 28.15 | 4.39 |
| MCL-1 | MCL-1_1_41 | 6.70 | 6.17 | 5.57 | 7.21 | 96.35 | 6.79 |
| MCL-1 | MCL-1_1_42 | 6.42 | 5.78 | 5.68 | 7.30 | 99.08 | 6.15 |
| MCL-1 | MCL-1_1_43 | 6.54 | 6.13 | 5.54 | 6.72 | 91.49 | 6.72 |
| MCL-1 | MCL-1_1_44 | 6.68 | 6.27 | 6.29 | 7.03 | 97.67 | 7.25 |
| MCL-1 | MCL-1_1_45 | 6.25 | 5.71 | 5.81 | 6.12 | 102.65 | 7.12 |
| MCL-1 | MCL-1_1_46 | 6.69 | 5.84 | 5.36 | 6.37 | 68.78 | 7.18 |
| MCL-1 | MCL-1_1_47 | 6.37 | 6.45 | 5.58 | 6.81 | 95.85 | 6.74 |
| MCL-1 | MCL-1_1_48 | 6.19 | 5.77 | 5.61 | 6.76 | 111.56 | 6.58 |
| MCL-1 | MCL-1_1_49 | 6.69 | 6.40 | 6.33 | 7.41 | 94.99 | 6.85 |
| MCL-1 | MCL-1_1_50 | 6.60 | 6.63 | 6.44 | 7.44 | 106.97 | 6.53 |
| MCL-1 | MCL-1_1_51 | 7.25 | 6.35 | 5.59 | 6.86 | 72.70 | 6.49 |
| MCL-1 | MCL-1_1_52 | 6.68 | 6.64 | 5.61 | 7.14 | 73.42 | 5.88 |
| MCL-1 | MCL-1_1_53 | 6.20 | 5.93 | 5.26 | 7.13 | 67.84 | 5.79 |
| MCL-1 | MCL-1_1_54 | 6.90 | 6.67 | 6.09 | 7.08 | 101.39 | 6.60 |
| MCL-1 | MCL-1_1_55 | 6.26 | 5.97 | 6.24 | 7.24 | 113.06 | 6.92 |
| MCL-1 | MCL-1_1_56 | 6.87 | 6.60 | 5.67 | 7.00 | 87.61 | 6.12 |
| MCL-1 | MCL-1_1_57 | 6.30 | 6.18 | 5.57 | 6.94 | 97.81 | 6.12 |
| MCL-1 | MCL-1_1_58 | 6.23 | 6.77 | 5.38 | 6.65 | 103.57 | 5.52 |
| MCL-1 | MCL-1_1_59 | 6.21 | 6.14 | 5.58 | 6.76 | 94.78 | 5.60 |
| MCL-1 | MCL-1_1_60 | 6.02 | 5.62 | 5.54 | 5.98 | 83.50 | 5.92 |
| MCL-1 | MCL-1_1_61 | 4.89 | 4.11 | 2.85 | 4.21 | 38.94 | 4.28 |
| MCL-1 | MCL-1_1_62 | 6.38 | 5.88 | 5.51 | 6.48 | 99.18 | 6.38 |
| MCL-1 | MCL-1_1_63 | 5.85 | 5.19 | 6.42 | 6.49 | 98.97 | 6.33 |
| MCL-1 | MCL-1_1_64 | 4.91 | 4.07 | 3.26 | 3.27 | 32.93 | 4.63 |
| MCL-1 | MCL-1_1_65 | 4.65 | 4.67 | 3.46 | 3.19 | 36.50 | 3.99 |
| KRAS^13,14^ | KRAS000 | 9.11 | 7.05 | 1.45 | 6.25 | 100.23 | 7.67 |
| KRAS | KRAS001 | 8.84 | 6.84 | 4.89 | 5.83 | 105.57 | 7.60 |
| KRAS | KRAS002 | 9.05 | 7.12 | 0.69 | 6.30 | 95.59 | 7.46 |
| KRAS | KRAS003 | 8.88 | 7.01 | 5.89 | 5.62 | 97.50 | 7.29 |
| KRAS | KRAS004 | 8.92 | 6.86 | 5.39 | 5.87 | 98.45 | 7.27 |
| KRAS | KRAS005 | 8.86 | 6.68 | 2.06 | 5.94 | 102.90 | 7.16 |
| KRAS | KRAS006 | 8.89 | 7.17 | 6.26 | 5.81 | 106.13 | 7.09 |
| KRAS | KRAS007 | 8.89 | 7.07 | 5.40 | 5.73 | 102.79 | 7.07 |
| KRAS | KRAS008 | 8.21 | 6.70 | 2.08 | 4.99 | 122.10 | 6.99 |
| KRAS | KRAS009 | 8.83 | 7.02 | 4.63 | 5.68 | 104.86 | 6.93 |
| KRAS | KRAS010 | 8.88 | 6.96 | 2.63 | 5.88 | 93.14 | 6.89 |
| KRAS | KRAS011 | 8.85 | 7.16 | 5.89 | 5.80 | 106.33 | 6.78 |
| KRAS | KRAS012 | 8.95 | 6.96 | 5.71 | 6.22 | 96.35 | 6.64 |
| KRAS | KRAS013 | 8.73 | 7.00 | 4.52 | 5.76 | 89.01 | 6.51 |
| KRAS | KRAS014 | 8.67 | 6.31 | 6.06 | 5.33 | 120.85 | 6.16 |
| KRAS | KRAS015 | 8.25 | 5.45 | 5.37 | 5.07 | 124.42 | 6.04 |
| KRAS | KRAS016 | 7.67 | 5.58 | 4.55 | 5.28 | 118.92 | 6.02 |
| KRAS | KRAS017 | 8.23 | 5.94 | 5.29 | 5.49 | 118.64 | 5.81 |
| KRAS | KRAS018 | 8.16 | 5.40 | 5.48 | 5.05 | 118.39 | 5.45 |
| KRAS | KRAS019 | 8.09 | 5.35 | 5.45 | 5.13 | 125.29 | 5.24 |
| KRAS | KRAS020 | 8.05 | 5.65 | 0.62 | 4.76 | 128.44 | 5.03 |
| KRAS | KRAS021 | 7.62 | 5.17 | 3.00 | 4.81 | 99.46 | 4.69 |
| KRAS | KRAS022 | 8.08 | 6.29 | 2.49 | 4.42 | 99.97 | 6.96 |
| KRAS | KRAS023 | 7.93 | 6.29 | -22.77 | 5.42 | 93.69 | 6.94 |
| KRAS | KRAS024 | 8.11 | 6.19 | 4.60 | 4.43 | 98.73 | 6.85 |
| KRAS | KRAS025 | 7.74 | 6.32 | 3.14 | 4.70 | 105.61 | 6.82 |
| KRAS | KRAS026 | 7.75 | 6.37 | 4.19 | 4.46 | 88.58 | 6.75 |
| KRAS | KRAS027 | 7.88 | 6.15 | 3.24 | 4.46 | 101.10 | 6.52 |
| KRAS | KRAS028 | 7.55 | 6.09 | 2.93 | 4.56 | 91.54 | 6.43 |
| KRAS | KRAS029 | 7.57 | 5.62 | 5.41 | 3.27 | 95.28 | 6.19 |
| KRAS | KRAS030 | 7.47 | 6.15 | 4.56 | 3.83 | 96.95 | 5.89 |
| KRAS | KRAS031 | 7.75 | 6.22 | 4.81 | 4.23 | 89.22 | 5.88 |
| KRAS | KRAS032 | 7.66 | 6.11 | 4.94 | 3.44 | 97.31 | 5.86 |
| KRAS | KRAS033 | 7.79 | 6.05 | 4.56 | 4.23 | 91.61 | 5.69 |
| KRAS | KRAS034 | 7.67 | 6.26 | 4.53 | 4.25 | 93.88 | 5.67 |
| KRAS | KRAS035 | 7.40 | 4.99 | 4.57 | 3.77 | 107.66 | 5.37 |
| KRAS | KRAS036 | 7.17 | 6.03 | 3.45 | 3.61 | 95.42 | 5.17 |
| KRAS | KRAS037 | 7.07 | 5.60 | 3.67 | 3.66 | 96.50 | 4.84 |

Section 5. Establishment of the CMX-2025 set and detailed results

In analog to the popular non-covalent CASF-2016 core set, we established a covalent binding affinity data set, namely CMX-2025, based on IC_50_, *K*_i_ or *K*_d_ reported in the literatures concerning development of covalent ligands, metalloproteins and halogenated drug. In this work, we collected 36 covalent drugs, 40 metalloproteins and 48 halogenated drugs. The detailed predictions and activity data are shown in Table S4.

**Table S5.** Predicted binding affinity against the CMX-2025 test set

| Target | Ligand_ID | DeepDOX1 | SableBind | ∆_vina_XGB | | TankBind | | DeepDOCK | | Expt. |
| --- | --- | --- | --- | --- | --- | --- | --- | --- | --- | --- |
| Covalent drug | | | | | | | | | | |
| BTK^15^ | BTK_1 | 7.68 | 7.74 | | 4.69 | | 6.89 | | 49.64 | 5.42 |
| BTK | BTK_2 | 8.21 | 7.97 | | 4.83 | | 6.95 | | 49.00 | 6.94 |
| BTK | BTK_3 | 8.46 | 8.82 | | 0.21 | | 7.35 | | 55.27 | 8.15 |
| BTK | BTK_4 | 8.26 | 8.93 | | 0.20 | | 7.15 | | 53.15 | 7.49 |
| JNK^15^ | JNK_1 | 7.25 | 6.61 | | 3.65 | | 6.36 | | 39.05 | 5.11 |
| JNK | JNK_2 | 7.60 | 6.99 | | 1.15 | | 7.52 | | 22.87 | 9.12 |
| JNK | JNK_3 | 7.12 | 5.99 | | 4.91 | | 7.29 | | 82.04 | 6.14 |
| JNK | JNK_4 | 8.10 | 7.39 | | 2.21 | | 7.07 | | 21.95 | 7.95 |
| AVP8^15^ | AVP8_1 | 7.53 | 9.84 | | 5.52 | | 7.19 | | 121.37 | 9.22 |
| AVP8 | AVP8_2 | 6.29 | 8.43 | | 4.15 | | 6.41 | | 84.44 | 5.56 |
| AVP8 | AVP8_3 | 6.81 | 7.57 | | 3.88 | | 6.68 | | 86.70 | 7.39 |
| AVP8 | AVP8_4 | 6.48 | 8.97 | | 4.13 | | 6.33 | | 84.60 | 6.02 |
| ERK2^16^ | ERK2_1 | 6.91 | 7.38 | | 2.74 | | 5.95 | | 35.73 | 6.43 |
| ERK2 | ERK2_2 | 7.38 | 7.34 | | 4.32 | | 6.93 | | 79.41 | 8.07 |
| ERK2 | ERK2_3 | 6.36 | 7.92 | | 4.23 | | 6.16 | | 28.88 | 7.37 |
| ERK2 | ERK2_4 | 7.05 | 8.50 | | 4.32 | | 6.62 | | 25.54 | 8.50 |
| 3Cpro^15^ | 3Cpro_1 | 6.89 | 6.45 | | 1.88 | | 6.78 | | 103.98 | 8.22 |
| 3Cpro | 3Cpro_2 | 6.82 | 7.57 | | 4.87 | | 5.46 | | 140.07 | 5.49 |
| 3Cpro | 3Cpro_3 | 7.15 | 7.45 | | 3.79 | | 6.01 | | 90.78 | 7.20 |
| 3Cpro | 3Cpro_4 | 6.57 | 8.76 | | 5.36 | | 6.62 | | 87.58 | 6.12 |
| Src^15^ | Src_1 | 5.96 | 6.18 | | 1.52 | | 4.67 | | 22.73 | 5.27 |
| Src | Src_2 | 6.03 | 6.31 | | 1.09 | | 5.35 | | 30.72 | 6.53 |
| Src | Src_3 | 6.22 | 6.94 | | 0.44 | | 6.87 | | 43.97 | 7.13 |
| Src | Src_4 | 5.90 | 6.13 | | 1.33 | | 6.89 | | 32.93 | 6.06 |
| FBA^17^ | FBA_1 | 4.36 | 5.22 | | 0.83 | | 1.59 | | 14.29 | 1.30 |
| FBA | FBA_2 | 4.96 | 6.09 | | 1.71 | | 1.08 | | 42.55 | 2.24 |
| FBA | FBA_3 | 5.53 | 6.32 | | 2.86 | | 1.03 | | 57.62 | 3.43 |
| FBA | FBA_4 | 5.05 | 6.94 | | 3.39 | | 1.38 | | 26.85 | 3.34 |
| CASP3^15^ | CASP3_1 | 7.33 | 8.97 | | 5.46 | | 7.35 | | 103.74 | 9.11 |
| CASP3 | CASP3_2 | 7.89 | 8.73 | | 5.73 | | 7.95 | | 128.84 | 8.52 |
| CASP3 | CASP3_3 | 6.50 | 5.74 | | 1.34 | | 6.56 | | 79.78 | 5.76 |
| CASP3 | CASP3_4 | 7.78 | 7.48 | | 4.57 | | 7.16 | | 112.09 | 7.75 |
| KRAS^13,14^ | KRAS_1 | 9.11 | 7.05 | | 1.43 | | 6.25 | | 100.23 | 7.67 |
| KRAS | KRAS_2 | 8.88 | 6.96 | | 4.65 | | 5.88 | | 93.14 | 6.89 |
| KRAS | KRAS_3 | 8.23 | 5.94 | | 5.27 | | 5.49 | | 118.64 | 5.81 |
| KRAS | KRAS_4 | 7.62 | 5.17 | | 2.81 | | 4.81 | | 99.46 | 4.69 |
| Metalloprotein | | | | | | | | | | |
| MAN2A1^18^ | MAN2A1_1 | 5.09 | 1.98 | | 0.50 | | 4.23 | | 102.85 | 2.11 |
| MAN2A1 | MAN2A1_2 | 5.22 | 3.63 | | 4.59 | | 5.50 | | 85.59 | 4.09 |
| MAN2A1 | MAN2A1_3 | 6.14 | 8.20 | | 4.11 | | 4.25 | | 93.00 | 7.44 |
| MAN2A1 | MAN2A1_4 | 5.56 | 6.69 | | 4.57 | | 3.69 | | 55.79 | 6.51 |
| APN^19^ | APN_1 | 6.26 | 9.12 | | 7.71 | | 6.76 | | 72.27 | 8.81 |
| APN | APN_2 | 6.18 | 7.55 | | 6.54 | | 6.23 | | 61.59 | 7.32 |
| APN | APN_3 | 5.37 | 3.93 | | 5.81 | | 5.31 | | 83.41 | 4.30 |
| APN | APN_4 | 5.72 | 7.96 | | 8.13 | | 5.44 | | 86.97 | 7.46 |
| CA12^18^ | CA12_1 | 6.39 | 7.75 | | 6.82 | | 7.19 | | 80.87 | 6.07 |
| CA12 | CA12_2 | 6.17 | 5.52 | | 5.45 | | 7.40 | | 52.05 | 5.48 |
| CA12 | CA12_3 | 7.55 | 9.76 | | 5.93 | | 8.05 | | 54.14 | 9.23 |
| CA12 | CA12_4 | 6.51 | 7.09 | | 6.93 | | 7.99 | | 68.09 | 6.78 |
| VIM-2^19^ | VIM-2_1 | 5.63 | 4.51 | | 2.71 | | 4.08 | | 45.08 | 5.57 |
| VIM-2 | VIM-2_2 | 4.97 | 6.06 | | 4.11 | | 4.46 | | 36.22 | 5.17 |
| VIM-2 | VIM-2_3 | 6.24 | 5.92 | | 4.89 | | 5.31 | | 35.84 | 7.72 |
| VIM-2 | VIM-2_4 | 5.41 | 5.06 | | 3.71 | | 3.23 | | 47.20 | 4.86 |
| TML^19^ | TML_1 | 7.53 | 7.04 | | 6.48 | | 6.69 | | 75.68 | 7.30 |
| TML | TML_2 | 5.60 | 5.09 | | 5.73 | | 5.87 | | 91.80 | 5.63 |
| TML | TML_3 | 6.68 | 2.87 | | 6.28 | | 5.51 | | 68.23 | 3.47 |
| TML | TML_4 | 7.06 | 5.77 | | 6.39 | | 6.17 | | 92.07 | 8.03 |
| MMP-8^19^ | MMP-8_1 | 5.38 | 3.80 | | 6.70 | | 6.31 | | 90.32 | 4.00 |
| MMP-8 | MMP-8_2 | 5.90 | 3.02 | | 7.05 | | 6.42 | | 43.22 | 2.92 |
| MMP-8 | MMP-8_3 | 6.46 | 6.59 | | 7.30 | | 7.56 | | 85.01 | 6.15 |
| MMP-8 | MMP-8_4 | 6.62 | 8.71 | | 6.44 | | 6.40 | | 64.28 | 8.27 |
| MMP-12^19^ | MMP-12_1 | 6.23 | 8.86 | | 5.62 | | 7.41 | | 62.56 | 8.69 |
| MMP-12 | MMP-12_2 | 6.11 | 7.99 | | 5.40 | | 6.87 | | 100.38 | 7.59 |
| MMP-12 | MMP-12_3 | 5.41 | 6.86 | | 5.53 | | 6.25 | | 62.63 | 5.84 |
| MMP-12 | MMP-12_4 | 8.91 | 9.83 | | 8.44 | | 8.16 | | 87.43 | 9.59 |
| HPPD^19^ | HPPD_1 | 5.58 | 6.80 | | 5.15 | | 4.91 | | 40.94 | 5.92 |
| HPPD | HPPD_2 | 6.01 | 8.73 | | 4.77 | | 5.17 | | 37.17 | 6.83 |
| HPPD | HPPD_3 | 6.40 | 9.19 | | 7.57 | | 5.66 | | 47.86 | 7.13 |
| HPPD | HPPD_4 | 6.34 | 8.30 | | 7.51 | | 6.58 | | 50.63 | 8.40 |
| nACE^20^ | nACE_1 | 5.00 | 7.61 | | 5.77 | | 7.70 | | 76.39 | 5.22 |
| nACE | nACE_2 | 6.72 | 11.43 | | 7.48 | | 8.27 | | 67.82 | 8.69 |
| nACE | nACE_3 | 7.72 | 6.86 | | 7.40 | | 7.88 | | 90.24 | 7.69 |
| nACE | nACE_4 | 5.23 | 8.03 | | 4.79 | | 6.79 | | 118.37 | 6.82 |
| cACE^20^ | cACE_1 | 6.78 | 8.77 | | 7.26 | | 7.68 | | 92.27 | 9.09 |
| cACE | cACE_2 | 6.63 | 8.76 | | 7.98 | | 7.71 | | 95.13 | 8.04 |
| cACE | cACE_3 | 6.47 | 8.83 | | 4.13 | | 6.93 | | 99.02 | 7.09 |
| cACE | cACE_4 | 6.00 | 8.37 | | 4.47 | | 7.87 | | 74.67 | 5.38 |
| Halogenated drug | | | | | | | | | | |
| Ser190^21^ | Ser190_1 | 5.44 | 5.06 | | 5.45 | | 5.27 | | 88.02 | 5.42 |
| Ser190 | Ser190_2 | 6.22 | 5.32 | | 5.78 | | 5.83 | | 102.08 | 7.11 |
| Ser190 | Ser190_3 | 6.09 | 6.82 | | 5.22 | | 5.34 | | 137.31 | 6.34 |
| Ser190 | Ser190_4 | 6.49 | 7.12 | | 5.45 | | 6.93 | | 156.77 | 5.74 |
| TRY1^22^ | TRY1_1 | 6.09 | 6.99 | | 3.61 | | 5.01 | | 33.14 | 4.84 |
| TRY1 | TRY1_2 | 6.63 | 7.87 | | 3.38 | | 5.21 | | 41.12 | 5.16 |
| TRY1 | TRY1_3 | 6.49 | 7.68 | | 4.75 | | 4.94 | | 53.27 | 6.65 |
| TRY1 | TRY1_4 | 6.65 | 8.15 | | 5.37 | | 5.20 | | 45.42 | 6.90 |
| THRβ1^23^ | THRβ1_1 | 6.96 | 8.81 | | 6.15 | | 7.34 | | 84.64 | 5.11 |
| THRβ1 | THRβ1_2 | 7.06 | 8.71 | | 6.06 | | 7.38 | | 88.86 | 5.15 |
| THRβ1 | THRβ1_3 | 7.21 | 8.80 | | 8.03 | | 7.11 | | 0.00 | 5.62 |
| THRβ1 | THRβ1_4 | 7.72 | 8.95 | | 5.60 | | 7.09 | | 95.02 | 6.33 |
| DHODH^24^ | DHODH_1 | 5.25 | 4.85 | | 5.52 | | 3.01 | | 0.00 | 3.69 |
| DHODH | DHODH_2 | 5.59 | 5.58 | | 5.16 | | 2.41 | | 0.00 | 5.37 |
| DHODH | DHODH_3 | 6.22 | 5.77 | | 6.69 | | 4.35 | | 2.23 | 6.55 |
| DHODH | DHODH_4 | 6.11 | 6.57 | | 7.28 | | 2.84 | | 4.50 | 5.0 |
| c-Met^25^ | c-Met_1 | 7.32 | 8.66 | | 5.92 | | 7.62 | | 68.44 | 5.10 |
| c-Met | c-Met_2 | 7.57 | 8.53 | | 6.45 | | 8.19 | | 53.38 | 6.33 |
| c-Met | c-Met_3 | 7.17 | 8.45 | | 6.17 | | 7.82 | | 70.95 | 5.54 |
| c-Met | c-Met_4 | 7.58 | 8.52 | | 5.75 | | 7.74 | | 69.72 | 5.76 |
| GPR119^26^ | GPR119_1 | 7.67 | 8.77 | | 8.38 | | 8.74 | | 89.14 | 8.23 |
| GPR119 | GPR119_2 | 7.32 | 8.52 | | 7.11 | | 8.16 | | 90.11 | 7.93 |
| GPR119 | GPR119_3 | 7.87 | 8.63 | | 8.23 | | 8.94 | | 89.99 | 9.16 |
| GPR119 | GPR119_4 | 7.82 | 8.53 | | 7.04 | | 8.67 | | 84.46 | 8.82 |
| Menin^27^ | Menin_1 | 6.52 | 6.97 | | 6.11 | | 6.72 | | 43.32 | 5.92 |
| Menin | Menin_2 | 6.57 | 7.66 | | 6.31 | | 7.24 | | 43.82 | 6.58 |
| Menin | Menin_3 | 6.78 | 7.90 | | 6.38 | | 7.27 | | 44.94 | 7.18 |
| Menin | Menin_4 | 7.11 | 7.58 | | 6.96 | | 7.17 | | 49.46 | 7.33 |
| CDK2^28^ | CDK2_1 | 6.15 | 6.70 | | 6.32 | | 6.38 | | 30.26 | 7.67 |
| CDK2 | CDK2_2 | 7.12 | 6.86 | | 6.52 | | 6.58 | | 17.16 | 8.09 |
| CDK2 | CDK2_3 | 6.11 | 6.59 | | 5.72 | | 6.40 | | 33.46 | 7.19 |
| CDK2 | CDK2_4 | 6.34 | 6.41 | | 6.79 | | 6.32 | | 22.06 | 7.95 |
| PI3Kα^29^ | PI3Kα_1 | 6.71 | 8.06 | | 6.82 | | 7.50 | | 52.74 | 9.46 |
| PI3Kα | PI3Kα_2 | 5.81 | 8.25 | | 6.54 | | 7.60 | | 64.03 | 9.72 |
| PI3Kα | PI3Kα_3 | 6.87 | 8.15 | | 6.27 | | 7.31 | | 57.99 | 10.37 |
| PI3Kα | PI3Kα_4 | 7.34 | 8.37 | | 6.26 | | 7.03 | | 56.15 | 9.82 |
| AURKA^30^ | AURKA_1 | 6.15 | 8.09 | | 5.46 | | 6.78 | | 63.95 | 8.00 |
| AURKA | AURKA_2 | 6.40 | 7.82 | | 5.65 | | 7.45 | | 66.68 | 8.43 |
| AURKA | AURKA_3 | 6.27 | 8.79 | | 6.25 | | 7.44 | | 70.17 | 9.09 |
| AURKA | AURKA_4 | 6.55 | 7.75 | | 6.37 | | 7.49 | | 67.53 | 8.60 |
| THRB^31^ | THRB_1 | 7.22 | 8.29 | | 6.95 | | 7.64 | | 151.51 | 9.09 |
| THRB | THRB_2 | 7.37 | 8.32 | | 6.90 | | 7.56 | | 142.00 | 9.56 |
| THRB | THRB_3 | 7.33 | 8.69 | | 6.37 | | 8.51 | | 109.28 | 10.00 |
| THRB | THRB_4 | 7.70 | 8.71 | | 7.09 | | 8.43 | | 159.70 | 10.37 |
| EDNRB^32^ | EDNRB_1 | 8.25 | 8.92 | | 6.47 | | 8.07 | | 70.05 | 6.33 |
| EDNRB | EDNRB_2 | 8.02 | 9.00 | | 7.33 | | 7.90 | | 107.02 | 5.82 |
| EDNRB | EDNRB_3 | 8.45 | 9.44 | | 6.51 | | 8.23 | | 92.93 | 6.33 |
| EDNRB | EDNRB_4 | 8.74 | 8.72 | | 6.21 | | 8.28 | | 104.85 | 6.66 |

Section 6. DeepDOX1 screening identifies FBPase ligands

Employing the DeepDOX1 framework, we performed drug design and activity prediction for FBPase. Initial conformational sampling was carried out with Cov_DOX, followed by binding affinity scoring with DeepDOX1. Based on synthetic accessibility considerations, we selected and synthesized compound 11t, one of the most readily accessible candidates. Experimental validation confirmed that its protein binding affinity outperformed that of compound 11n, consistent with our computational predictions.

**Table S6.** DeepDOX1 screening identifies FBPase ligands

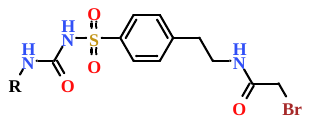

| Compds | *R* | DeepDOX1 |
| --- | --- | --- |
| 11 | 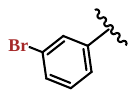 | 7.65 |
| 11n | 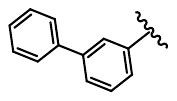 | 7.74 |
| 11t | 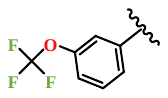 | 7.89 |
| 11u | 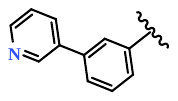 | 8.27 |
| 11v | 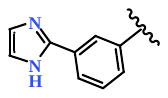 | 8.13 |
| 11w | 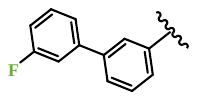 | 8.21 |
| 11-Virtu-6 | 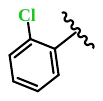 | 7.38 |
| 11-Virtu-7 | 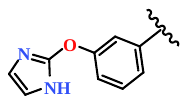 | 7.35 |
| 11-Virtu-8 | 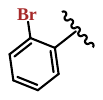 | 7.81 |
| 11-Virtu-9 | 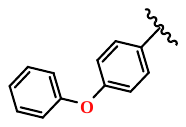 | 7.22 |
| 11-Virtu-10 | 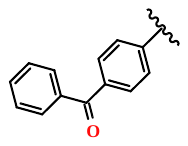 | 7.33 |
| 11-Virtu-11 | 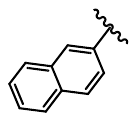 | 7.21 |
| 11-Virtu-12 | 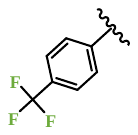 | 7.27 |
| 11-Virtu-13 | 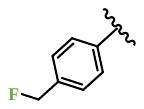 | 7.00 |
| 11-Virtu-14 | 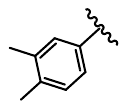 | 7.10 |
| 11-Virtu-15 | 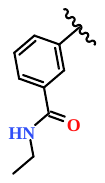 | 7.35 |
| 11-Virtu-16 | 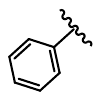 | 6.77 |
| 11-Virtu-17 | 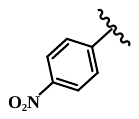 | 7.09 |
| 11-Virtu-18 | 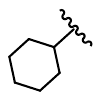 | 6.75 |
| 11-Virtu-19 | 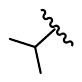 | 6.34 |
| 11-Virtu-20 | 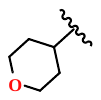 | 6.61 |

Figure S1. Predicted binding structure of 11t(A),11u(B),11v(C),11w(D) employing a combination of Cov_DOX and DeepDOX1.

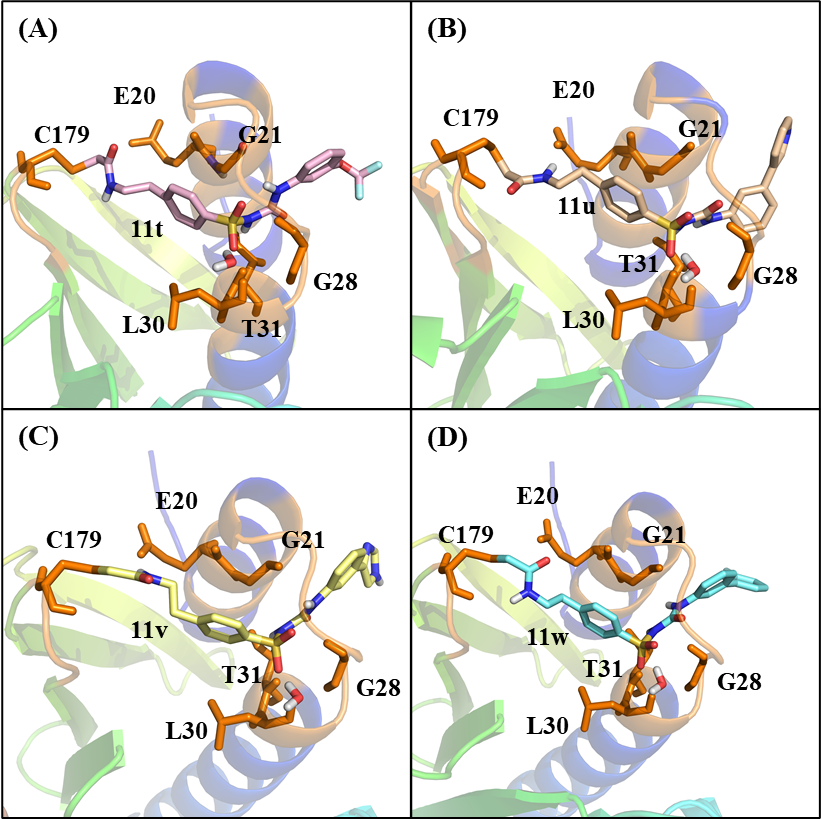

Section 7. Detailed FBPase DeepDOX1 Calculation

For the prediction of drug-binding poses to FBPase, we implemented a multi-stage computational pipeline. Initial conformational sampling was carried out using Cov_DOX, leveraging AutoDock4 and GSADOCK for initial pose generation. The resulting poses were geometry-optimized with the PM7 semiempirical method. Following this, single-point energy evaluations were conducted at the ωB97X-D/6-311+G(d,p) level of theory to obtain higher-level electronic energies. The conformational ensemble was then scored using the DeepDOX1 model, from which the pose with the most favorable score was selected as the predicted binding mode.

**Table S7.** DeepDOX1 results of different binding conformations

| Binding Conformation^a^ | DeepDOX1 Score (relative to Top1 pose) |
| --- | --- |
| Conf. 1 | -0.18 |
| Conf. 2 | -0.06 |
| Conf. 3 | 0.00 |
| Conf. 4 | -1.23 |
| Conf. 5 | -0.78 |
| Conf. 6 | -0.29 |

a. The number of conformation denotes its order ranked by molecular docking and semi-empirical theory.

Section 8. Detailed description of ligand atomic feature

For the ligand atomic feature, all ligand non-hydrogen atoms are divided into 64 atomic types based on the its element type and the atoms bonded with it, see Table S3 for a detailed description. Each row of the ligand atomic feature matrix is the one-hot encoding of a ligand atom atomic type number. Note that each atomic type number might correspond to one atomic type or several atomic types with similar chemical bonding environment. The atomic type encoding is in the form of ZXXXXX, while Z is the element type. The first X is the number of hydrogen atom bonded with corresponding atom. The second X is the number of carbon atom bonded with corresponding atom. The third X is the number of nitrogen atom and phosphorus atom bonded with corresponding atom. The fourth X is the number of oxygen atom and sulfur atom bonded with corresponding atom. The fifth X is the number of halogen atom bonded with corresponding atom. For example, C12000 is the carbon of benzene and N21000 is the nitrogen atom of -NH2 group.

**Table S8.** 64 atomic types defined for ligand atomic feature.

| Atomic type number | Atomic type encoding |
| --- | --- |
| 1 | C12000 |
| 2 | O01000 O00000 |
| 3 | C22000 C13000 C22010 C20000 |
| 4 | C03000 |
| 5 | N12000 N11000 N10010 |
| 6 | C01110 C10110 |
| 7 | C31000 C30000 |
| 8 | C12100 C11200 C10210 |
| 9 | O11000 O10000 O10100 O10010 |
| 10 | C21100 |
| 11 | C12010 C12001 C21001 C20110 |
| 12 | C02100 |
| 13 | C02010 |
| 14 | O02000 |
| 15 | N02000 |
| 16 | C21010 |
| 17 | C01020 C00030 C10020 |
| 18 | N03000 |
| 19 | C11100 |
| 20 | N21000 N20100 |
| 21 | C01200 C01101 C00201 |
| 22 | O00100 O00110 |
| 23 | C02001 |
| 24 | O00010 |
| 25 | F01000 I01000 Cl01000 Br01000 |
| 26 | N31000 N22000 N13000 N04000 N12100 N21100 N12010 N21010 N03100 N03010 N20110 |
| 27 | C30010 |
| 28 | C30100 C20200 |
| 29 | C10200 |
| 30 | C00300 |
| 31 | S02000 S00200 S02010 |
| 32 | N01100 |
| 33 | C00210 |
| 34 | O01100 O01010 |
| 35 | C11020 |
| 36 | P00040 P01030 P00140 P10030 P01040 P03010 P00310 |
| 37 | S01120 |
| 38 | C04000 C02200 |
| 39 | C11110 |
| 40 | N02100 N01110 |
| 41 | C03010 C02110 C02002 C03001 C11002 C10012 C01102 Si03010 Si02020 C11011 C02011 C10120 |
| 42 | C00120 |
| 43 | C11010 C01011 C20010 |
| 44 | N11100 N10200 N10110 N10100 |
| 45 | N11010 |
| 46 | N02010 |
| 47 | C01003 C00013 |
| 48 | C03100 |
| 49 | N01000 N00100 |
| 50 | C01100 C11000 C00110 C00200 |
| 51 | N01010 |
| 52 | O00200 |
| 53 | C02000 |
| 54 | C21000 C20100 |
| 55 | C02020 |
| 56 | N01020 N01200 N00120 |
| 57 | S02020 |
| 58 | N00200 N00110 |
| 59 | N20010 |
| 60 | C20020 |
| 61 | S11000 S01000 S01100 S01010 S10100 |
| 62 | S01030 S00130 S00220 S00040 |
| 63 | B01020 |
| 64 | Transition metal ion |

Section 9. Detailed description of the Rdkit feature

For molecular representation, we curated 12 RDKit-derived descriptors deemed critical for drug-likeness and protein-ligand interactions. The selected descriptors are summarized in the table below.

**Table S9.** Detailed data of the Rdkit feature

| Molecular Descriptors | Meaning |
| --- | --- |
| MolWt | The average molecular weight of the molecule |
| LogP | Atom-based calculation of LogP and MR using Crippen’s approach |
| Num_HDonors | Number of Hydrogen Bond Donors |
| Num_HAcceptors | Number of Hydrogen Bond Acceptors |
| QED | Calculate the weighted sum of ADS mapped properties |
| TPSA | Topological Polar Surface Area (TPSA) |
| NO_Count | Number of Nitrogens and Oxygens |
| FractionCSP3 | the fraction of C atoms that are SP3 hybridized |
| Num_RotatableBonds | Number of Rotatable Bonds |
| Num_AliphaticRings | the number of aliphatic (containing at least one non-aromatic bond) rings for a molecule |
| Num_SaturatedRings | the number of saturated rings for a molecule |
| RingCount | the number of rings for a molecule |

Section 10. DOX Calculation of protein–ligand binding energy

The initial crystallographic structure of the protein–ligand complex was first optimized using the PM7 semiempirical method using the MOPAC package. Subsequent single-point energy calculations were performed at the ωB97X-D/6-311G+(d,p) level of theory using eXtended ONIOM with Gaussian 09. We computed the overall binding energy from the single-point energy difference of the complex and its isolated components, along with a detailed decomposition of the individual residue contributions. Entropic contributions were approximated at the PM7 level, with the entropic change upon binding simplified to half the entropy of the free ligand.

Section 11. Used PDBBind PDB ID for Training set

| 10gs | 11gs | 13gs | 16pk | 184l | 185l | 186l | 187l | 188l | 1a07 |
| --- | --- | --- | --- | --- | --- | --- | --- | --- | --- |
| 1a1b | 1a1c | 1a1e | 1a28 | 1a42 | 1a46 | 1a4g | 1a4h | 1a4q | 1a4w |
| 1a52 | 1a5v | 1a61 | 1a69 | 1a8i | 1a94 | 1a99 | 1a9m | 1a9q | 1aaq |
| 1abf | 1acj | 1add | 1adl | 1ado | 1afk | 1ag9 | 1agw | 1ahx | 1ahy |
| 1ai4 | 1ai5 | 1ai6 | 1ai7 | 1aid | 1aj6 | 1ajn | 1ajp | 1ajq | 1akq |
| 1akr | 1akt | 1aku | 1akv | 1akw | 1al7 | 1al8 | 1alw | 1amk | 1amw |
| 1ao0 | 1apb | 1aq1 | 1aqj | 1avd | 1avn | 1awh | 1ax0 | 1ax1 | 1ax2 |
| 1axr | 1axs | 1axz | 1azl | 1b05 | 1b0h | 1b1h | 1b2h | 1b2i | 1b32 |
| 1b38 | 1b3f | 1b3g | 1b3h | 1b3l | 1b40 | 1b42 | 1b46 | 1b4h | 1b4z |
| 1b51 | 1b52 | 1b56 | 1b58 | 1b5g | 1b5h | 1b5i | 1b5j | 1b6h | 1b6j |
| 1b6k | 1b74 | 1b7h | 1b8n | 1b8o | 1b9j | 1b9s | 1b9t | 1b9v | 1bai |
| 1bap | 1bbz | 1bcd | 1bhf | 1bhx | 1bil | 1bji | 1bkm | 1bky | 1bl4 |
| 1bl6 | 1bl7 | 1bma | 1bmb | 1bmk | 1bmm | 1bn1 | 1bn4 | 1bnn | 1bnq |
| 1bnt | 1bnv | 1bnw | 1bo5 | 1bp0 | 1bq4 | 1br5 | 1br6 | 1br8 | 1bv7 |
| 1bv9 | 1bwa | 1bwb | 1bxo | 1bxq | 1bzf | 1bzh | 1bzj | 1c12 | 1c1u |
| 1c29 | 1c2t | 1c3e | 1c3x | 1c4u | 1c50 | 1c5c | 1c5f | 1c5o | 1c5q |
| 1c7e | 1c7f | 1c83 | 1c84 | 1c85 | 1c86 | 1c87 | 1c88 | 1c8k | 1c8v |
| 1c9d | 1cbr | 1cbx | 1ce5 | 1ceb | 1cet | 1cil | 1cim | 1cin | 1cj1 |
| 1cka | 1ckb | 1cnx | 1cny | 1cpi | 1cqp | 1cr6 | 1csh | 1csi | 1csr |
| 1css | 1ctr | 1cw2 | 1cwb | 1cwc | 1cx9 | 1cyn | 1czc | 1cze | 1czk |
| 1czl | 1czo | 1czr | 1d04 | 1d09 | 1d3d | 1d3p | 1d3q | 1d4i | 1d4k |
| 1d4p | 1d4t | 1d4y | 1d5r | 1d6n | 1d6v | 1d6w | 1d7j | 1dar | 1db1 |
| 1dbb | 1dbk | 1det | 1df8 | 1dfo | 1dgm | 1dhi | 1dhj | 1di8 | 1di9 |
| 1dif | 1dis | 1dkd | 1dl7 | 1dm2 | 1dmb | 1dmp | 1drj | 1drk | 1drv |
| 1dtq | 1dtt | 1dub | 1dud | 1duv | 1dva | 1dwd | 1dx6 | 1dy4 | 1dzj |
| 1dzk | 1dzm | 1dzp | 1e00 | 1e02 | 1e06 | 1e1v | 1e1x | 1e1y | 1e2k |
| 1e2l | 1e3g | 1e3v | 1e4h | 1e5a | 1e6q | 1e6s | 1e72 | 1e8h | 1e9h |
| 1eb2 | 1ebw | 1ebz | 1ec0 | 1ec3 | 1ec9 | 1ecq | 1ecv | 1eef | 1eei |
| 1efi | 1efy | 1egh | 1ej4 | 1ejn | 1ek1 | 1ek2 | 1ela | 1elb | 1elc |
| 1eld | 1ele | 1elr | 1em6 | 1ent | 1enu | 1epo | 1epp | 1erb | 1err |
| 1eve | 1evh | 1ez9 | 1ezq | 1f0q | 1f0t | 1f0u | 1f28 | 1f2o | 1f3e |
| 1f3j | 1f4e | 1f4f | 1f4g | 1f4x | 1f5k | 1f5l | 1f73 | 1f74 | 1f8a |
| 1f8b | 1f8c | 1f8d | 1f8e | 1f90 | 1f92 | 1f9e | 1f9g | 1fbm | 1fch |
| 1fcx | 1fcy | 1fcz | 1fd0 | 1fd7 | 1fe3 | 1fgi | 1fhd | 1fig | 1fiv |
| 1fj4 | 1fkb | 1fkf | 1fkg | 1fkh | 1fki | 1fkn | 1fkw | 1fl3 | 1flm |
| 1flr | 1fo0 | 1fpc | 1fpi | 1fpy | 1fq5 | 1fq6 | 1fq7 | 1fq8 | 1fsg |
| 1fta | 1ftj | 1ftk | 1ftl | 1ftm | 1fv9 | 1fvt | 1fvv | 1fw0 | 1fwe |
| 1fyr | 1fzj | 1fzk | 1fzo | 1fzq | 1g1d | 1g2o | 1g30 | 1g32 | 1g3f |
| 1g3m | 1g45 | 1g46 | 1g4j | 1g50 | 1g52 | 1g53 | 1g54 | 1g5f | 1g74 |
| 1g7f | 1g7g | 1g7p | 1g7v | 1g85 | 1g98 | 1g9t | 1ga8 | 1ga9 | 1gai |
| 1gca | 1gfy | 1gfz | 1ggn | 1gi7 | 1gj5 | 1gj7 | 1gj8 | 1gja | 1gjb |
| 1gjc | 1gjd | 1gni | 1gnj | 1gnm | 1gnn | 1gno | 1gny | 1gqs | 1grp |
| 1gsf | 1gsz | 1gt4 | 1gu1 | 1gu3 | 1gui | 1gux | 1gvx | 1gwm | 1gwq |
| 1gwr | 1gx8 | 1gym | 1gz3 | 1gz4 | 1gzv | 1h01 | 1h08 | 1h0a | 1h0w |
| 1h1p | 1h1s | 1h24 | 1h27 | 1h35 | 1h36 | 1h37 | 1h39 | 1h3a | 1h3b |
| 1h3c | 1h46 | 1h4n | 1h4w | 1h5u | 1h60 | 1h61 | 1h62 | 1h6e | 1h6h |
| 1h7a | 1h9z | 1ha2 | 1hbv | 1hdt | 1hee | 1hef | 1hge | 1hgi | 1hgj |
| 1hgt | 1hi3 | 1hih | 1hii | 1him | 1hiv | 1hk1 | 1hk2 | 1hlf | 1hmr |
| 1hms | 1hmt | 1hn2 | 1hp5 | 1hpv | 1hpx | 1hqg | 1hqh | 1hrn | 1hs6 |
| 1hsl | 1hte | 1htf | 1htg | 1hti | 1hty | 1hvi | 1hvj | 1hvk | 1hvl |
| 1hvr | 1hvs | 1hvy | 1hwr | 1hxb | 1hxk | 1hxw | 1hyo | 1i00 | 1i1e |
| 1i2s | 1i32 | 1i33 | 1i37 | 1i41 | 1i43 | 1i48 | 1i5r | 1i6v | 1i7c |
| 1i7g | 1i7i | 1i7m | 1i7z | 1i80 | 1i8z | 1i91 | 1i9l | 1i9n | 1i9o |
| 1i9p | 1i9q | 1ida | 1idb | 1ie9 | 1iep | 1iew | 1ihy | 1ii5 | 1iih |
| 1iiq | 1ik4 | 1ikx | 1iky | 1il3 | 1il4 | 1il5 | 1il9 | 1inf | 1ing |
| 1inh | 1inq | 1iq1 | 1is0 | 1it6 | 1iup | 1ivp | 1iy7 | 1iyl | 1izh |
| 1izi | 1j01 | 1j14 | 1j15 | 1j16 | 1j17 | 1j4r | 1jak | 1jaq | 1jet |
| 1jeu | 1jev | 1jg0 | 1jii | 1jij | 1jik | 1jil | 1jj9 | 1jjk | 1jld |
| 1jlq | 1jmf | 1jmg | 1jmi | 1jn2 | 1jn4 | 1jp5 | 1jpl | 1jq3 | 1jqd |
| 1jqe | 1jqy | 1jr1 | 1jt1 | 1jtq | 1juf | 1juq | 1jut | 1juy | 1jvp |
| 1jwt | 1jyq | 1jys | 1jzs | 1k03 | 1k06 | 1k08 | 1k1j | 1k1m | 1k1n |
| 1k1o | 1k1p | 1k4g | 1k4h | 1kak | 1kc5 | 1kcs | 1kdk | 1ke5 | 1ke6 |
| 1ke7 | 1ke8 | 1ke9 | 1kf0 | 1kf6 | 1kfy | 1kkq | 1kl3 | 1kl5 | 1klg |
| 1klu | 1km3 | 1kmv | 1koj | 1kqb | 1ktt | 1kv2 | 1kv5 | 1kwr | 1kyv |
| 1kz8 | 1kzk | 1kzn | 1l0a | 1l2s | 1l5q | 1l5r | 1l5s | 1l7x | 1l83 |
| 1l8g | 1laf | 1lag | 1lah | 1lan | 1lb6 | 1lbf | 1lbk | 1lee | 1lek |
| 1lev | 1lf2 | 1lf8 | 1lf9 | 1lfo | 1lgw | 1lhu | 1li2 | 1li3 | 1li6 |
| 1lkk | 1lkl | 1loq | 1lor | 1los | 1lox | 1lpk | 1lpz | 1lq2 | 1lqd |
| 1lrt | 1lst | 1lt6 | 1lvc | 1lvk | 1lzo | 1lzq | 1m0b | 1m13 | 1m2p |
| 1m2q | 1m2r | 1m4h | 1m51 | 1m5b | 1m5c | 1m5d | 1m5e | 1m5f | 1m74 |
| 1m7q | 1m7y | 1m83 | 1m9n | 1mai | 1mau | 1maw | 1mcz | 1me7 | 1me8 |
| 1mes | 1met | 1meu | 1mfg | 1mik | 1mkd | 1ml1 | 1mm6 | 1mm7 | 1moq |
| 1mq5 | 1mqd | 1mqg | 1mqh | 1mqi | 1mqj | 1mrw | 1mrx | 1ms0 | 1ms7 |
| 1msm | 1msn | 1mto | 1mtr | 1mui | 1mv0 | 1mx1 | 1mxu | 1my2 | 1my3 |
| 1my4 | 1mzs | 1n0t | 1n1m | 1n1t | 1n1v | 1n2v | 1n3i | 1n43 | 1n46 |
| 1n4h | 1n4k | 1n5z | 1n8v | 1n94 | 1n9a | 1n9m | 1nax | 1nd5 | 1ndj |
| 1ndv | 1ndw | 1ndy | 1ndz | 1nf8 | 1ngw | 1nhg | 1nhu | 1nhv | 1nhw |
| 1nhx | 1nhz | 1nj1 | 1nj5 | 1njb | 1njd | 1njs | 1nl9 | 1nli | 1nlo |
| 1nlp | 1nlt | 1nm6 | 1nmk | 1nnb | 1nnk | 1nnu | 1nny | 1no6 | 1noi |
| 1noj | 1nok | 1nox | 1np0 | 1npa | 1nq0 | 1nq7 | 1nt1 | 1ntk | 1ntv |
| 1nu3 | 1nvs | 1nw4 | 1nw5 | 1nw7 | 1nwl | 1nxy | 1nyx | 1nz7 | 1o0f |
| 1o0m | 1o0n | 1o0o | 1o2n | 1o2p | 1o2q | 1o42 | 1o46 | 1o49 | 1o4f |
| 1o4g | 1o4h | 1o4j | 1o4l | 1o4m | 1o4n | 1o4o | 1o4p | 1o4q | 1o5c |
| 1o5m | 1o5r | 1o6h | 1o6i | 1o6q | 1o6r | 1o79 | 1o86 | 1oai | 1oar |
| 1oba | 1ocn | 1od8 | 1odi | 1odj | 1ody | 1oe7 | 1oe8 | 1ofz | 1ogu |
| 1oh4 | 1oi9 | 1oif | 1oim | 1oiq | 1oir | 1oit | 1oiu | 1oiy | 1oj5 |
| 1ok7 | 1okv | 1okw | 1oky | 1okz | 1ol1 | 1ol2 | 1om1 | 1om9 | 1ony |
| 1onz | 1os5 | 1osg | 1oss | 1oth | 1ouk | 1ouy | 1ov3 | 1ove | 1ow6 |
| 1ow7 | 1ow8 | 1owd | 1owe | 1owi | 1owj | 1owk | 1ox9 | 1oz0 | 1ozv |
| 1p0y | 1p19 | 1p1o | 1p28 | 1p2a | 1p2g | 1p4r | 1p4u | 1p5e | 1p93 |
| 1pa9 | 1pb8 | 1pbk | 1pbq | 1pcg | 1pdz | 1pf7 | 1pf8 | 1pfy | 1pg2 |
| 1pgp | 1ph0 | 1phw | 1pip | 1pkx | 1pl0 | 1pmn | 1pmu | 1pmv | 1pot |
| 1ppm | 1pr1 | 1prm | 1pum | 1pvn | 1pwp | 1pwu | 1pwy | 1pxh | 1pxj |
| 1pxk | 1pxl | 1pxm | 1pxo | 1pxp | 1py1 | 1pyg | 1pyn | 1pyw | 1pzi |
| 1pzo | 1q0b | 1q1g | 1q1m | 1q3d | 1q3w | 1q41 | 1q4l | 1q4w | 1q4x |
| 1q5k | 1q5l | 1q63 | 1q65 | 1q66 | 1q6k | 1q6p | 1q6s | 1q6t | 1q72 |
| 1q83 | 1q8w | 1q95 | 1q9d | 1qaw | 1qb9 | 1qbn | 1qbr | 1qbs | 1qbt |
| 1qbu | 1qf2 | 1qft | 1qj1 | 1qj6 | 1qj7 | 1qja | 1qjb | 1qk3 | 1qka |
| 1qkb | 1qkn | 1qku | 1qm4 | 1qm5 | 1qon | 1qpb | 1qpe | 1qpl | 1qq9 |
| 1qs4 | 1qsc | 1qti | 1qvt | 1qvu | 1qwf | 1qwu | 1qx1 | 1qxk | 1qxl |
| 1qy1 | 1qy2 | 1qyg | 1r0p | 1r1h | 1r1i | 1r4w | 1r58 | 1r5h | 1r5n |
| 1r5w | 1r6g | 1r6n | 1r6z | 1r78 | 1r9l | 1rbo | 1rbp | 1rd4 | 1rdt |
| 1rev | 1rin | 1rjk | 1rlp | 1rlq | 1rnm | 1ro7 | 1rpa | 1rpf | 1rpj |
| 1rq2 | 1rql | 1rri | 1rrw | 1rry | 1rs4 | 1rsd | 1rsi | 1rst | 1rt1 |
| 1rt2 | 1rt9 | 1rtf | 1rth | 1rv1 | 1rw8 | 1rwq | 1rwx | 1rzx | 1s19 |
| 1s26 | 1s39 | 1s4d | 1s50 | 1s9t | 1s9v | 1sc8 | 1sh9 | 1shd | 1siv |
| 1sje | 1skj | 1sld | 1sle | 1slg | 1sm3 | 1sme | 1so2 | 1soj | 1sps |
| 1sqb | 1sqc | 1sqn | 1sqo | 1sqp | 1sqq | 1sqt | 1sr7 | 1sre | 1ssq |
| 1stc | 1stp | 1str | 1sts | 1sve | 1svg | 1svh | 1sw1 | 1sw2 | 1swi |
| 1swk | 1swn | 1swp | 1swr | 1syo | 1sz0 | 1szd | 1szm | 1t13 | 1t1r |
| 1t1s | 1t29 | 1t2v | 1t46 | 1t48 | 1t49 | 1t4j | 1t4s | 1t4v | 1t5a |
| 1t79 | 1t7f | 1t7j | 1t7r | 1ta2 | 1ta6 | 1tet | 1tg5 | 1thl | 1thr |
| 1ths | 1thz | 1tjp | 1tkx | 1tl7 | 1tl9 | 1tlp | 1tmn | 1tog | 1toi |
| 1toj | 1tok | 1tom | 1tou | 1tow | 1tqf | 1trd | 1tsl | 1tsm | 1tsv |
| 1tsy | 1tt1 | 1ttm | 1tu6 | 1tv6 | 1tve | 1tvr | 1tze | 1u0g | 1u1w |
| 1u2r | 1u2y | 1u33 | 1u3q | 1u3r | 1u59 | 1u65 | 1u71 | 1u8t | 1u9e |
| 1ua4 | 1udt | 1udu | 1uho | 1ui0 | 1uj0 | 1ujj | 1ujk | 1uk0 | 1uk1 |
| 1uml | 1umw | 1unl | 1uou | 1upf | 1upk | 1ur9 | 1urc | 1urw | 1usi |
| 1usk | 1utc | 1utj | 1utl | 1utm | 1utn | 1utp | 1utt | 1utz | 1uu7 |
| 1uu8 | 1uu9 | 1uv5 | 1uvr | 1uw6 | 1uwb | 1uwh | 1uwt | 1uwu | 1uxa |
| 1uxb | 1uy6 | 1uy7 | 1uy8 | 1uy9 | 1uyc | 1uyd | 1uye | 1uyf | 1uyg |
| 1uyh | 1uyi | 1uym | 1uys | 1uz1 | 1uz4 | 1uz8 | 1v0l | 1v0m | 1v0n |
| 1v0o | 1v0p | 1v1j | 1v1k | 1v2h | 1v2j | 1v2l | 1v2m | 1v2s | 1v2u |
| 1v2v | 1v41 | 1v79 | 1v7a | 1vcj | 1vcu | 1vea | 1veb | 1vfn | 1vj5 |
| 1vja | 1vjb | 1vjc | 1vjd | 1vjy | 1vkj | 1vot | 1vr1 | 1vrt | 1vru |
| 1vwf | 1vwl | 1vwn | 1vyf | 1vyg | 1vyq | 1vyw | 1vyz | 1vzq | 1w0x |
| 1w0z | 1w13 | 1w1p | 1w1t | 1w1v | 1w1y | 1w25 | 1w2g | 1w2x | 1w3j |
| 1w3k | 1w4l | 1w4p | 1w4q | 1w51 | 1w5v | 1w5w | 1w5x | 1w5y | 1w6h |
| 1w6j | 1w6r | 1w76 | 1w7h | 1w80 | 1w82 | 1w83 | 1w84 | 1w8l | 1w8m |
| 1w96 | 1wax | 1way | 1wbn | 1wbo | 1wbs | 1wbt | 1wbv | 1wbw | 1wc6 |
| 1wcc | 1wcq | 1wdn | 1wdq | 1wdy | 1we2 | 1wht | 1wm1 | 1wok | 1wug |
| 1wvj | 1wxz | 1wzy | 1x1z | 1x38 | 1x39 | 1x6u | 1x70 | 1x76 | 1x78 |
| 1x7b | 1x7e | 1x7q | 1x7r | 1x8b | 1x8s | 1xap | 1xbb | 1xbc | 1xbo |
| 1xd0 | 1xdg | 1xfv | 1xh4 | 1xh5 | 1xh6 | 1xh7 | 1xh8 | 1xhy | 1xjd |
| 1xk9 | 1xka | 1xkk | 1xlx | 1xlz | 1xm4 | 1xm6 | 1xmu | 1xn2 | 1xnx |
| 1xnz | 1xo2 | 1xoe | 1xog | 1xom | 1xon | 1xoq | 1xor | 1xos | 1xow |
| 1xoz | 1xq0 | 1xqc | 1xr9 | 1xs7 | 1xsc | 1xt8 | 1xuc | 1xud | 1xws |
| 1xxe | 1xxi | 1y0l | 1y1m | 1y1z | 1y20 | 1y2a | 1y2b | 1y2d | 1y2e |
| 1y2f | 1y2g | 1y2h | 1y2j | 1y2k | 1y4z | 1y57 | 1y6a | 1y6b | 1y6q |
| 1y98 | 1ya4 | 1ybo | 1yc4 | 1yda | 1ydb | 1ydk | 1yds | 1yet | 1yhs |
| 1yid | 1ym2 | 1ym4 | 1ynd | 1yon | 1ype | 1ypg | 1ypj | 1yrs | 1ysg |
| 1ysi | 1yt7 | 1yt9 | 1yvf | 1yvh | 1yvm | 1yw2 | 1ywr | 1yxd | 1yy4 |
| 1yye | 1yyr | 1yys | 1z1h | 1z1r | 1z34 | 1z4n | 1z4u | 1z5m | 1z6p |
| 1z6q | 1z6s | 1z71 | 1z9h | 1zaf | 1zaj | 1zc9 | 1zd3 | 1zdp | 1zea |
| 1zeo | 1zfk | 1zfq | 1zge | 1zgv | 1zh7 | 1zhl | 1zkk | 1zkl | 1zkn |
| 1zky | 1zoe | 1zog | 1zoh | 1zrz | 1zsb | 1zsf | 1zub | 1zuc | 1zxv |
| 1zyj | 1zyr | 1zz2 | 1zzl | 220l | 223l | 2a0c | 2a2x | 2a3a | 2a3b |
| 2a3c | 2a4l | 2a4m | 2a4z | 2a5b | 2a5c | 2a5s | 2a5u | 2a8g | 2aa6 |
| 2aa9 | 2aac | 2aay | 2adm | 2adu | 2ael | 2afw | 2afx | 2agv | 2aj8 |
| 2am1 | 2am2 | 2ama | 2amv | 2ank | 2anl | 2anm | 2ao6 | 2aoc | 2aod |
| 2aoe | 2aov | 2aox | 2aq7 | 2aqb | 2aqu | 2ate | 2ath | 2auz | 2avi |
| 2aw1 | 2ax6 | 2ax9 | 2axi | 2ay1 | 2ay2 | 2ay3 | 2ay4 | 2ay5 | 2ay6 |
| 2ay7 | 2ay8 | 2ay9 | 2ayp | 2az5 | 2az8 | 2az9 | 2azc | 2azm | 2azr |
| 2b07 | 2b1g | 2b1i | 2b1v | 2b1z | 2b2v | 2b4l | 2b4m | 2b52 | 2b53 |
| 2b54 | 2b55 | 2b7a | 2b7f | 2b8l | 2b8v | 2b9a | 2baj | 2bak | 2bal |
| 2ban | 2bb7 | 2bcd | 2bdj | 2bdy | 2be2 | 2bfq | 2bfr | 2bgd | 2bge |
| 2bgn | 2bgr | 2bjm | 2bkz | 2bmc | 2bo4 | 2bpm | 2bpv | 2bpy | 2bqv |
| 2br8 | 2brc | 2brg | 2brh | 2brm | 2brn | 2bro | 2bt9 | 2btr | 2bts |
| 2bua | 2bub | 2buc | 2bvd | 2bvr | 2bvx | 2bxt | 2bxu | 2byh | 2byi |
| 2byp | 2bys | 2bz5 | 2bz8 | 2c1a | 2c1b | 2c1n | 2c1q | 2c2l | 2c3j |
| 2c3k | 2c3l | 2c4g | 2c4v | 2c57 | 2c5n | 2c5o | 2c5x | 2c5y | 2c68 |
| 2c69 | 2c6e | 2c6i | 2c6k | 2c6l | 2c6m | 2c6n | 2c6o | 2c8w | 2c8x |
| 2c8y | 2c90 | 2c93 | 2c97 | 2c9b | 2c9d | 2c9t | 2cbj | 2cc7 | 2ccb |
| 2ccs | 2cct | 2ccu | 2ce9 | 2cej | 2ceq | 2cer | 2ces | 2cex | 2cf8 |
| 2cf9 | 2cgf | 2cgu | 2cgv | 2cgw | 2cgx | 2chm | 2chw | 2chx | 2chz |
| 2cia | 2ckm | 2cle | 2clf | 2clh | 2cli | 2clk | 2clm | 2clv | 2clx |
| 2cm8 | 2cmb | 2cmc | 2cmf | 2cmo | 2cn0 | 2cn8 | 2cne | 2csm | 2csn |
| 2ctc | 2cvd | 2d06 | 2d0k | 2d1x | 2d3u | 2d3z | 2dbl | 2drc | 2dri |
| 2duv | 2dw7 | 2dwx | 2dxs | 2e1w | 2e2p | 2e2r | 2e5y | 2e7f | 2e7l |
| 2e9n | 2e9o | 2e9u | 2e9v | 2ea4 | 2eh8 | 2epn | 2er6 | 2er9 | 2erz |
| 2esm | 2etk | 2etm | 2etr | 2euf | 2evc | 2evl | 2evm | 2evo | 2ewb |
| 2ewp | 2ewy | 2exc | 2exg | 2exm | 2ez5 | 2ez7 | 2f01 | 2f0z | 2f10 |
| 2f14 | 2f18 | 2f1a | 2f1b | 2f1g | 2f2c | 2f2h | 2f34 | 2f35 | 2f3e |
| 2f3f | 2f3k | 2f4b | 2f4j | 2f6t | 2f6v | 2f6y | 2f6z | 2f70 | 2f71 |
| 2f7i | 2f7o | 2f7p | 2f80 | 2f81 | 2f8g | 2f8i | 2fah | 2fai | 2fdp |
| 2ff1 | 2fgh | 2fgu | 2fgv | 2fhy | 2fie | 2fix | 2fjm | 2fjn | 2fjp |
| 2fky | 2fl2 | 2fl5 | 2fl6 | 2fle | 2flh | 2fmb | 2fme | 2fo4 | 2fq6 |
| 2fqx | 2fqy | 2fr3 | 2fr8 | 2fsv | 2fts | 2fum | 2fvc | 2fw3 | 2fw6 |
| 2fwp | 2fwy | 2fwz | 2fx7 | 2fxu | 2fxv | 2fyv | 2fzc | 2fzg | 2fzk |
| 2g01 | 2g0g | 2g0h | 2g1q | 2g1r | 2g1y | 2g24 | 2g2r | 2g5u | 2g79 |
| 2g8r | 2g94 | 2g97 | 2g9q | 2g9r | 2g9u | 2g9v | 2g9x | 2gbi | 2gc8 |
| 2gcd | 2gde | 2gdo | 2gej | 2gek | 2gfa | 2gfd | 2gfs | 2gg0 | 2gg2 |
| 2gg3 | 2gg5 | 2gg7 | 2gga | 2ggb | 2ggd | 2gh7 | 2gh9 | 2ghg | 2gj4 |
| 2gj5 | 2gl0 | 2glm | 2glp | 2gm9 | 2gmv | 2gmx | 2gnf | 2gnh | 2gni |
| 2gnj | 2gnl | 2go4 | 2gpp | 2gqn | 2gss | 2gst | 2gtk | 2gtv | 2gu8 |
| 2gvj | 2gz2 | 2gz7 | 2gz8 | 2h02 | 2h03 | 2h15 | 2h21 | 2h23 | 2h2d |
| 2h2e | 2h2g | 2h2h | 2h2j | 2h3e | 2h42 | 2h4g | 2h4k | 2h5a | 2h5e |
| 2h6b | 2h6q | 2h6t | 2h8h | 2h96 | 2h9m | 2ha2 | 2ha3 | 2ha6 | 2ha7 |
| 2hai | 2haw | 2hb9 | 2hd1 | 2hd6 | 2hdq | 2hdr | 2hds | 2hdu | 2hdx |
| 2hfp | 2hhn | 2hiw | 2hiz | 2hj4 | 2hjb | 2hk5 | 2hmb | 2hmv | 2hmw |
| 2hnc | 2hnx | 2hny | 2hoc | 2hog | 2hpa | 2hrp | 2hs1 | 2hs2 | 2hu6 |
| 2hug | 2hvc | 2hvx | 2hw2 | 2hwg | 2hwh | 2hwi | 2hxm | 2hxq | 2hy0 |
| 2hz4 | 2hzi | 2hzl | 2hzn | 2hzy | 2i0a | 2i0d | 2i0e | 2i0g | 2i0h |
| 2i0v | 2i19 | 2i1r | 2i2b | 2i2c | 2i3v | 2i40 | 2i4j | 2i4t | 2i4u |
| 2i4v | 2i4w | 2i4x | 2i4z | 2i5j | 2i6a | 2i6b | 2i7c | 2i80 | 2idw |
| 2ieh | 2ien | 2ieo | 2igv | 2igw | 2igy | 2ihj | 2ihq | 2iit | 2iiv |
| 2iko | 2iku | 2il2 | 2ilp | 2ima | 2imb | 2in6 | 2io6 | 2ioa | 2ipo |
| 2iqg | 2isc | 2isv | 2isw | 2itk | 2ito | 2itp | 2itt | 2ity | 2itz |
| 2iu0 | 2iuz | 2iv9 | 2ivu | 2iw6 | 2iw8 | 2iw9 | 2iws | 2iwu | 2iyf |
| 2izx | 2j27 | 2j2i | 2j3q | 2j47 | 2j4a | 2j4g | 2j4k | 2j4z | 2j50 |
| 2j62 | 2j6m | 2j75 | 2j77 | 2j79 | 2j7b | 2j7d | 2j7e | 2j7f | 2j7g |
| 2j7x | 2j9h | 2j9l | 2j9n | 2ja3 | 2jaj | 2jb5 | 2jbk | 2jbl | 2jbo |
| 2jbp | 2jbu | 2jc0 | 2jdl | 2jdo | 2jds | 2jdt | 2jdv | 2jf4 | 2jfh |
| 2jfz | 2jg0 | 2jgs | 2jiu | 2jiw | 2jj3 | 2jjb | 2jjk | 2jjr | 2jk9 |
| 2jke | 2jkk | 2jkm | 2jko | 2jkq | 2jkr | 2jkt | 2jle | 2jnp | 2jqi |
| 2jql | 2jst | 2jup | 2jxr | 2k1q | 2k2r | 2kbs | 2kce | 2kdh | 2khh |
| 2kmx | 2kup | 2l1r | 2l6j | 2l7u | 2l84 | 2lbv | 2lcs | 2lh8 | 2lkk |
| 2lnw | 2lo6 | 2lsv | 2ltv | 2ltw | 2ltx | 2ltz | 2ly0 | 2lya | 2lyb |
| 2lzg | 2m0u | 2m0v | 2m41 | 2mas | 2mc1 | 2nm1 | 2nmy | 2nmz | 2nnd |
| 2nng | 2nnk | 2nno | 2nnp | 2nnq | 2nns | 2nnv | 2no3 | 2np8 | 2nq7 |
| 2nsj | 2nsl | 2nsx | 2nt7 | 2nta | 2ntf | 2nv7 | 2nw4 | 2nwl | 2nww |
| 2nxl | 2o0u | 2o1v | 2o22 | 2o2u | 2o3p | 2o3z | 2o48 | 2o4h | 2o4j |
| 2o4k | 2o4l | 2o4n | 2o4p | 2o4s | 2o4z | 2o5d | 2o5k | 2o63 | 2o64 |
| 2o65 | 2o7v | 2o8h | 2o9i | 2o9j | 2o9k | 2o9v | 2oa0 | 2oag | 2oah |
| 2oax | 2obj | 2oei | 2of2 | 2of4 | 2off | 2og8 | 2ogy | 2ogz | 2oh0 |
| 2oh4 | 2ohk | 2ohl | 2ohm | 2ohp | 2ohq | 2ohr | 2ohs | 2oht | 2ohu |
| 2ohv | 2oi9 | 2oic | 2oiq | 2oj9 | 2ojf | 2ojg | 2ojj | 2olb | 2ole |
| 2on3 | 2onb | 2onc | 2oo8 | 2op9 | 2oph | 2oqi | 2oqv | 2osf | 2ot1 |
| 2ov4 | 2ovv | 2ovy | 2ow3 | 2ow6 | 2ow7 | 2ow9 | 2owb | 2oxd | 2oxn |
| 2oxx | 2oxy | 2oyk | 2oyl | 2oym | 2oz5 | 2oz7 | 2ozr | 2p0d | 2p2a |
| 2p2h | 2p2i | 2p33 | 2p3a | 2p3b | 2p3c | 2p3d | 2p3g | 2p3i | 2p4j |
| 2p4s | 2p7a | 2p7g | 2p7z | 2p83 | 2p8n | 2p8s | 2p9a | 2pax | 2pbw |
| 2pcp | 2pe0 | 2pe1 | 2pe2 | 2peh | 2pem | 2pfy | 2pg2 | 2pgj | 2pgl |
| 2pgz | 2ph8 | 2ph9 | 2pix | 2piy | 2pj1 | 2pj3 | 2pj4 | 2pj6 | 2pj7 |
| 2pj9 | 2pja | 2pjb | 2pjc | 2pk5 | 2pk6 | 2pks | 2pl0 | 2pmc | 2pmk |
| 2pmn | 2pnc | 2poq | 2pou | 2pov | 2pow | 2pq9 | 2pqb | 2pqc | 2pqj |
| 2pql | 2pqz | 2prj | 2psj | 2psu | 2psv | 2pt9 | 2pu0 | 2pu2 | 2pv1 |
| 2pvh | 2pvj | 2pvk | 2pvl | 2pvm | 2pvn | 2pvu | 2pvv | 2pwc | 2pwd |
| 2pwr | 2pyi | 2pym | 2pyn | 2pyy | 2pzi | 2pzy | 2q11 | 2q15 | 2q1l |
| 2q1q | 2q2a | 2q2c | 2q2z | 2q38 | 2q54 | 2q55 | 2q5k | 2q63 | 2q64 |
| 2q6c | 2q6h | 2q70 | 2q72 | 2q7m | 2q7q | 2q7y | 2q80 | 2q88 | 2q89 |
| 2q8g | 2q8h | 2q8m | 2q8s | 2q92 | 2q93 | 2q94 | 2q95 | 2q96 | 2qbs |
| 2qbu | 2qbw | 2qc6 | 2qcd | 2qcf | 2qcg | 2qch | 2qci | 2qcm | 2qd6 |
| 2qd7 | 2qd8 | 2qd9 | 2qe2 | 2qe5 | 2qf6 | 2qfo | 2qft | 2qfu | 2qg0 |
| 2qg2 | 2qhc | 2qhn | 2qhr | 2qhy | 2qhz | 2qi0 | 2qi1 | 2qi3 | 2qi4 |
| 2qi5 | 2qi6 | 2qi7 | 2qiq | 2qju | 2qk5 | 2qk8 | 2qlm | 2qln | 2qm7 |
| 2qm9 | 2qmd | 2qmf | 2qmg | 2qn1 | 2qn2 | 2qn3 | 2qnb | 2qnn | 2qnp |
| 2qnx | 2qo1 | 2qo8 | 2qoa | 2qoe | 2qoh | 2qp6 | 2qp8 | 2qpq | 2qpu |
| 2qrg | 2qrh | 2qrk | 2qrl | 2qrm | 2qrp | 2qrq | 2qt5 | 2qt9 | 2qtg |
| 2qtn | 2qtr | 2qtt | 2qu3 | 2qu5 | 2qu6 | 2qv7 | 2qw1 | 2qwb | 2qwc |
| 2qwd | 2qwe | 2qwf | 2qwg | 2qyk | 2qyl | 2qyn | 2qzk | 2qzl | 2qzr |
| 2qzx | 2r02 | 2r03 | 2r05 | 2r0u | 2r0y | 2r0z | 2r1w | 2r1x | 2r1y |
| 2r23 | 2r2b | 2r2m | 2r2w | 2r38 | 2r3c | 2r3f | 2r3h | 2r3i | 2r3j |
| 2r3k | 2r3l | 2r3m | 2r3n | 2r3o | 2r3p | 2r3t | 2r3w | 2r43 | 2r4b |
| 2r58 | 2r5a | 2r5b | 2r5q | 2r6f | 2r6w | 2r6y | 2r75 | 2r7b | 2r8q |
| 2r9b | 2r9s | 2r9x | 2rc8 | 2rc9 | 2rcb | 2rcu | 2rd6 | 2reg | 2rf2 |
| 2rfh | 2rg5 | 2rg6 | 2rgp | 2rgu | 2rip | 2rjp | 2rk7 | 2rk8 | 2rka |
| 2rke | 2rkf | 2rkg | 2rkm | 2rkn | 2rku | 2rl5 | 2rly | 2rm0 | 2rnx |
| 2rr4 | 2sfp | 2sim | 2std | 2toh | 2tsr | 2upj | 2uue | 2uuo | 2uup |
| 2uw0 | 2uw3 | 2uw4 | 2uw5 | 2uw6 | 2uw7 | 2uw8 | 2uwd | 2uxi | 2uxu |
| 2uxx | 2uxz | 2uy3 | 2uy5 | 2uyi | 2uym | 2uyq | 2uyw | 2uz6 | 2uz9 |
| 2uzb | 2uze | 2uzl | 2uzn | 2uzo | 2uzv | 2v0n | 2v0z | 2v10 | 2v11 |
| 2v12 | 2v13 | 2v16 | 2v22 | 2v25 | 2v2c | 2v2h | 2v2q | 2v2v | 2v3d |
| 2v3e | 2v54 | 2v57 | 2v58 | 2v59 | 2v5a | 2v5x | 2v77 | 2v7d | 2v95 |
| 2va5 | 2va6 | 2va7 | 2vaq | 2vb8 | 2vba | 2vc7 | 2vc9 | 2vcb | 2vci |
| 2vcj | 2vcq | 2vcw | 2vcx | 2vd0 | 2vd4 | 2vd7 | 2vfk | 2vgo | 2vgp |
| 2vhj | 2vhq | 2vi5 | 2vin | 2vio | 2vip | 2viq | 2vj1 | 2vj7 | 2vj8 |
| 2vj9 | 2vjx | 2vk2 | 2vk6 | 2vl4 | 2vle | 2vmc | 2vmd | 2vmf | 2vnm |
| 2vnn | 2vnt | 2vo4 | 2vpn | 2vpo | 2vpp | 2vqt | 2vr4 | 2vrj | 2vrx |
| 2vt3 | 2vta | 2vtd | 2vte | 2vth | 2vti | 2vtj | 2vtl | 2vtm | 2vtn |
| 2vto | 2vtp | 2vtq | 2vtr | 2vts | 2vtt | 2vu3 | 2vuk | 2vur | 2vv9 |
| 2vvo | 2vvs | 2vvt | 2vw1 | 2vw2 | 2vwc | 2vwu | 2vwv | 2vwx | 2vx0 |
| 2vxa | 2vxn | 2vyt | 2w05 | 2w06 | 2w0j | 2w17 | 2w1c | 2w1d | 2w1e |
| 2w1f | 2w1g | 2w1h | 2w1i | 2w3l | 2w3o | 2w4i | 2w54 | 2w5g | 2w5i |
| 2w67 | 2w68 | 2w6c | 2w6m | 2w6n | 2w6o | 2w6p | 2w6q | 2w6z | 2w70 |
| 2w71 | 2w7x | 2w7y | 2w8f | 2w8g | 2w8j | 2w8w | 2w8y | 2w92 | 2w97 |
| 2w9h | 2wa3 | 2wa4 | 2waj | 2wb5 | 2wbb | 2wbd | 2wc3 | 2wc4 | 2wcx |
| 2wd1 | 2wd3 | 2wd7 | 2we3 | 2web | 2wec | 2weh | 2wei | 2wej | 2weo |
| 2weq | 2wev | 2wez | 2wf0 | 2wf1 | 2wf2 | 2wf3 | 2wgj | 2wgs | 2who |
| 2wi1 | 2wi2 | 2wi3 | 2wi4 | 2wi5 | 2wi6 | 2wi7 | 2wib | 2wih | 2wk2 |
| 2wk6 | 2wks | 2wky | 2wl5 | 2wly | 2wlz | 2wm0 | 2wmr | 2wmu | 2wmv |
| 2wmw | 2wmx | 2wnl | 2won | 2wos | 2wot | 2wou | 2wp1 | 2wpa | 2wpb |
| 2wqb | 2wqp | 2wr8 | 2wtc | 2wtd | 2wti | 2wtj | 2wtw | 2wtx | 2wu6 |
| 2wu7 | 2wuf | 2wxd | 2wxf | 2wxg | 2wxh | 2wxi | 2wxj | 2wxk | 2wxl |
| 2wxm | 2wxn | 2wxo | 2wxp | 2wxq | 2wxv | 2wyi | 2wzm | 2wzs | 2wzy |
| 2x09 | 2x0y | 2x24 | 2x2k | 2x2l | 2x2m | 2x2r | 2x38 | 2x52 | 2x5o |
| 2x6d | 2x6e | 2x6j | 2x6k | 2x7c | 2x7d | 2x7o | 2x7s | 2x7t | 2x7x |
| 2x81 | 2x8d | 2x8e | 2x8i | 2x95 | 2x97 | 2x9e | 2x9f | 2xab | 2xae |
| 2xaf | 2xaj | 2xb7 | 2xb9 | 2xbj | 2xcg | 2xch | 2xck | 2xcs | 2xd6 |
| 2xd9 | 2xda | 2xde | 2xdk | 2xdx | 2xel | 2xey | 2xez | 2xf0 | 2xfi |
| 2xfk | 2xg3 | 2xg5 | 2xg9 | 2xgm | 2xgo | 2xgs | 2xhr | 2xhs | 2xht |
| 2xhx | 2xib | 2xix | 2xiy | 2xiz | 2xj0 | 2xj1 | 2xj2 | 2xjg | 2xjj |
| 2xjx | 2xk4 | 2xk6 | 2xk7 | 2xk8 | 2xk9 | 2xkc | 2xkd | 2xke | 2xkf |
| 2xm1 | 2xm2 | 2xm8 | 2xm9 | 2xml | 2xmy | 2xn3 | 2xn5 | 2xn7 | 2xne |
| 2xng | 2xnm | 2xnn | 2xno | 2xnp | 2xo8 | 2xog | 2xoi | 2xp2 | 2xp3 |
| 2xp4 | 2xp5 | 2xp6 | 2xp7 | 2xp8 | 2xpa | 2xpb | 2xpc | 2xpk | 2xqq |
| 2xrw | 2xsb | 2xtk | 2xuc | 2xui | 2xup | 2xvd | 2xvn | 2xwd | 2xx2 |
| 2xx4 | 2xxr | 2xxt | 2xxw | 2xxx | 2xxy | 2xy9 | 2xyd | 2xyr | 2xyt |
| 2xzg | 2y06 | 2y07 | 2y1n | 2y1o | 2y1w | 2y2k | 2y3p | 2y4k | 2y4s |
| 2y54 | 2y56 | 2y57 | 2y58 | 2y59 | 2y5f | 2y5l | 2y67 | 2y68 | 2y76 |
| 2y7i | 2y7k | 2y7p | 2y7w | 2y8c | 2y8i | 2y8o | 2y8q | 2ya6 | 2ya7 |
| 2ya8 | 2yac | 2yay | 2yb0 | 2ybp | 2ybu | 2ych | 2ycq | 2ycr | 2ycs |
| 2yde | 2ydf | 2ydi | 2ydj | 2ydk | 2ydo | 2ydt | 2ydv | 2ydw | 2ye9 |
| 2yek | 2yel | 2yem | 2yer | 2yex | 2yfa | 2yfx | 2yg2 | 2yga | 2ygf |
| 2ygu | 2yhd | 2yi0 | 2yi5 | 2yi7 | 2yiq | 2yir | 2yis | 2yiu | 2yiv |
| 2yiw | 2yix | 2yjw | 2yjx | 2yk1 | 2yk9 | 2ykb | 2ykc | 2yke | 2ykj |
| 2yln | 2ylo | 2ylp | 2ylq | 2ym3 | 2ym4 | 2ym5 | 2ym6 | 2ym7 | 2ym8 |
| 2yme | 2ymt | 2ypi | 2ypp | 2ywp | 2yxj | 2yz3 | 2z1w | 2z4o | 2z60 |
| 2z7r | 2z8e | 2z92 | 2z9g | 2zas | 2zaz | 2zb0 | 2zbk | 2zc9 | 2zcs |
| 2zdt | 2zdz | 2ze1 | 2zfp | 2zg1 | 2zg3 | 2zgx | 2zjf | 2zjw | 2zkj |
| 2zlf | 2zm1 | 2zm3 | 2zmd | 2zmm | 2zn7 | 2zns | 2znt | 2znu | 2zo3 |
| 2zof | 2zoq | 2zpk | 2zu4 | 2zv2 | 2zv9 | 2zwz | 2zx5 | 2zx6 | 2zx7 |
| 2zx8 | 2zx9 | 2zxa | 2zxb | 2zxd | 2zxg | 2zym | 2zyn | 2zz1 | 3a29 |
| 3a2c | 3a2o | 3a3y | 3a4o | 3a4p | 3a5y | 3a6t | 3a73 | 3abt | 3acx |
| 3ad7 | 3ad8 | 3ads | 3adt | 3adu | 3adv | 3agm | 3ah8 | 3ai8 | 3aje |
| 3ama | 3amb | 3anq | 3anr | 3ans | 3ant | 3ao2 | 3ap7 | 3apc | 3aqt |
| 3ara | 3arn | 3arr | 3art | 3arw | 3arx | 3arz | 3as1 | 3as2 | 3as3 |
| 3asx | 3at1 | 3at3 | 3at4 | 3atl | 3atp | 3atu | 3atv | 3au6 | 3av9 |
| 3ava | 3avg | 3avj | 3avk | 3avl | 3avz | 3ax5 | 3axk | 3axm | 3axz |
| 3ay0 | 3ay9 | 3aya | 3ayd | 3b0w | 3b24 | 3b25 | 3b26 | 3b28 | 3b2q |
| 3b2t | 3b2w | 3b3x | 3b4f | 3b4p | 3b50 | 3b5j | 3b66 | 3b67 | 3b78 |
| 3b7j | 3b7u | 3b95 | 3b9g | 3bc4 | 3bcs | 3be2 | 3be9 | 3bea | 3bel |
| 3bex | 3bft | 3bfu | 3bgl | 3bgp | 3bgq | 3bgs | 3bi0 | 3bi6 | 3biz |
| 3bjc | 3bkk | 3bkl | 3bl0 | 3bl1 | 3bl7 | 3bl9 | 3bla | 3blr | 3blt |
| 3bm6 | 3bm9 | 3bmn | 3bmy | 3bpr | 3bqc | 3bqn | 3br9 | 3bra | 3brn |
| 3bsc | 3bt9 | 3bti | 3btj | 3bu1 | 3bug | 3buh | 3bv3 | 3bva | 3bvb |
| 3bwj | 3bx5 | 3bxe | 3bxf | 3bxg | 3bxs | 3bxz | 3bym | 3bys | 3bz3 |
| 3c1k | 3c1n | 3c1x | 3c2f | 3c2o | 3c2r | 3c2u | 3c39 | 3c43 | 3c4f |
| 3c4h | 3c5u | 3c6t | 3c6u | 3c79 | 3c7n | 3c7p | 3c7q | 3c84 | 3c8e |
| 3cbp | 3cbs | 3ccb | 3ccc | 3ccn | 3cct | 3ccw | 3ccz | 3cd0 | 3cd7 |
| 3cd8 | 3cdb | 3cde | 3ce0 | 3ce3 | 3cf1 | 3cf8 | 3cf9 | 3cft | 3cgf |
| 3cgo | 3chd | 3chg | 3chp | 3chq | 3chs | 3cib | 3cic | 3cid | 3cj2 |
| 3cj3 | 3cj5 | 3cjo | 3ck7 | 3ck8 | 3ckb | 3cke | 3ckr | 3ckt | 3ckz |
| 3cl0 | 3cl2 | 3clp | 3cm2 | 3cn0 | 3co9 | 3coh | 3cow | 3cp9 | 3cpb |
| 3cpc | 3cph | 3cpj | 3cqw | 3cso | 3cth | 3ctt | 3cvk | 3cwe | 3cwj |
| 3cwk | 3cx9 | 3cy2 | 3cyw | 3cyx | 3cyz | 3cz1 | 3czv | 3d04 | 3d0b |
| 3d14 | 3d1g | 3d1x | 3d1y | 3d1z | 3d20 | 3d28 | 3d2e | 3d2r | 3d2t |
| 3d3p | 3d4l | 3d4y | 3d50 | 3d51 | 3d52 | 3d5m | 3d6o | 3d6p | 3d78 |
| 3d7b | 3d7h | 3d7m | 3d7z | 3d83 | 3d8w | 3d8y | 3d8z | 3d91 | 3d9n |
| 3d9o | 3d9p | 3d9v | 3d9z | 3da6 | 3da9 | 3daj | 3daz | 3db6 | 3db8 |
| 3dbd | 3dbs | 3dbu | 3dc2 | 3dc3 | 3dcc | 3dct | 3dcv | 3dcw | 3ddf |
| 3ddg | 3ddp | 3ddq | 3ddu | 3deh | 3dei | 3dej | 3dg8 | 3dgq | 3dhk |
| 3di6 | 3djo | 3djp | 3djq | 3dk1 | 3dkf | 3dln | 3dnd | 3dne | 3dng |
| 3dnj | 3dnt | 3dog | 3doy | 3doz | 3dp0 | 3dp1 | 3dp2 | 3dp3 | 3dp4 |
| 3dp9 | 3dpf | 3dpk | 3drf | 3drg | 3dri | 3drp | 3drr | 3drs | 3ds6 |
| 3dst | 3dt1 | 3dtc | 3du8 | 3dux | 3dv5 | 3dvp | 3dwb | 3dx0 | 3dx3 |
| 3dx4 | 3dxh | 3dxj | 3dxk | 3dxm | 3dya | 3dyo | 3dz2 | 3dzt | 3e01 |
| 3e12 | 3e2m | 3e3b | 3e3c | 3e51 | 3e5u | 3e62 | 3e63 | 3e64 | 3e6y |
| 3e7b | 3e7o | 3e85 | 3e8n | 3e9h | 3e9i | 3eax | 3eb1 | 3ebb | 3ebh |
| 3ebi | 3ebl | 3ebo | 3ecn | 3ed0 | 3ee2 | 3efj | 3efk | 3efr | 3eft |
| 3efw | 3egt | 3ehn | 3ehx | 3eid | 3eig | 3eio | 3ej1 | 3ejp | 3ejq |
| 3ejs | 3ejt | 3eju | 3eka | 3ekn | 3eko | 3ekp | 3ekr | 3eks | 3ekt |
| 3ekv | 3ekx | 3eky | 3el0 | 3el5 | 3el7 | 3el8 | 3el9 | 3elc | 3elj |
| 3emg | 3eml | 3eoc | 3eos | 3eou | 3eq7 | 3eqb | 3eql | 3eqr | 3erk |
| 3ern | 3ert | 3esj | 3ess | 3et7 | 3eta | 3evc | 3evd | 3ew2 | 3ewh |
| 3ewz | 3exo | 3eyg | 3eyh | 3eys | 3eyu | 3ezr | 3ezv | 3f07 | 3f17 |
| 3f2a | 3f3t | 3f3u | 3f3v | 3f3w | 3f48 | 3f5j | 3f5k | 3f5l | 3f66 |
| 3f69 | 3f6g | 3f6h | 3f70 | 3f78 | 3f7b | 3f7u | 3f7z | 3f81 | 3f82 |
| 3f88 | 3f8c | 3f8f | 3f8s | 3f8w | 3f9n | 3faa | 3fas | 3fat | 3fbr |
| 3fc1 | 3fc2 | 3fc8 | 3fcb | 3fcf | 3fci | 3fcl | 3fdn | 3fee | 3feg |
| 3fei | 3fej | 3ff3 | 3ff6 | 3ffp | 3fgc | 3fh5 | 3fh7 | 3fh8 | 3fhb |
| 3fhe | 3fhr | 3fi2 | 3fi3 | 3fjg | 3fjz | 3fk1 | 3fkt | 3fl5 | 3fl8 |
| 3fl9 | 3fmz | 3fnu | 3fpm | 3fq7 | 3fqh | 3fqk | 3fql | 3fr2 | 3fr4 |
| 3fr5 | 3frg | 3frz | 3ft5 | 3ft8 | 3ftq | 3fts | 3ftu | 3ftv | 3ftw |
| 3fty | 3ftz | 3fu0 | 3fu3 | 3fu5 | 3fu6 | 3fud | 3fue | 3fuf | 3fuh |
| 3fui | 3fuj | 3fuk | 3ful | 3fum | 3fun | 3fup | 3fvg | 3fvk | 3fvl |
| 3fvn | 3fw3 | 3fwv | 3fx6 | 3fxb | 3fxv | 3fxw | 3fyj | 3fyk | 3fz1 |
| 3fzr | 3fzt | 3g0b | 3g0e | 3g0g | 3g15 | 3g19 | 3g1d | 3g1m | 3g2h |
| 3g2i | 3g2j | 3g2k | 3g2l | 3g2u | 3g2y | 3g30 | 3g32 | 3g34 | 3g35 |
| 3g3n | 3g45 | 3g4g | 3g4i | 3g4l | 3g58 | 3g5d | 3g5y | 3g6g | 3g6h |
| 3g6m | 3g6z | 3g70 | 3g72 | 3g86 | 3g8e | 3g8o | 3g90 | 3g9e | 3g9l |
| 3g9n | 3ga5 | 3gba | 3gbe | 3gc4 | 3gc7 | 3gcq | 3gcs | 3gcu | 3gcv |
| 3gdt | 3gf2 | 3gfe | 3gfw | 3ggu | 3ggv | 3gi4 | 3gi5 | 3gi6 | 3gjd |
| 3gjw | 3gk2 | 3gk4 | 3gm0 | 3gn7 | 3gnv | 3gol | 3gp0 | 3gpe | 3gpo |
| 3gqo | 3gqz | 3gs6 | 3gs7 | 3gsg | 3gsm | 3gss | 3gst | 3gt9 | 3gtc |
| 3gur | 3gus | 3guz | 3gvb | 3gvu | 3gws | 3gwt | 3gwu | 3gwv | 3gww |
| 3gwx | 3gx0 | 3gxl | 3gxt | 3gy3 | 3gy7 | 3gyn | 3gz9 | 3h0a | 3h0b |
| 3h0e | 3h0j | 3h0q | 3h0s | 3h0v | 3h0w | 3h0y | 3h0z | 3h23 | 3h2c |
| 3h2n | 3h30 | 3h5b | 3h5s | 3h6z | 3h78 | 3h98 | 3h9f | 3h9k | 3h9o |
| 3ha6 | 3hab | 3hac | 3hav | 3hb4 | 3hb8 | 3hcm | 3hdm | 3hdn | 3hec |
| 3heg | 3hek | 3hf6 | 3hf8 | 3hfb | 3hfj | 3hfv | 3hfz | 3hhk | 3hhu |
| 3hjo | 3hk1 | 3hkq | 3hku | 3hkw | 3hky | 3hl7 | 3hl8 | 3hll | 3hmm |
| 3hmo | 3hmp | 3hmv | 3hng | 3hnz | 3ho2 | 3ho9 | 3hp2 | 3hp5 | 3hp9 |
| 3hq5 | 3hqh | 3hqw | 3hqy | 3hqz | 3hr1 | 3hrb | 3hrf | 3hs4 | 3hub |
| 3huc | 3hv3 | 3hv4 | 3hv5 | 3hv6 | 3hv7 | 3hv8 | 3hvc | 3hw1 | 3hx3 |
| 3hxb | 3hxc | 3hxd | 3hxe | 3hzk | 3hzm | 3hzv | 3hzy | 3i02 | 3i0r |
| 3i0s | 3i1y | 3i28 | 3i4b | 3i4y | 3i51 | 3i5n | 3i5z | 3i60 | 3i6c |
| 3i6m | 3i6o | 3i6z | 3i73 | 3i7b | 3i7c | 3i7e | 3i7g | 3i81 | 3i97 |
| 3ia6 | 3ibi | 3ibl | 3ibn | 3ibu | 3idp | 3ie3 | 3iej | 3ieo | 3ies |
| 3ifl | 3ifo | 3ifp | 3ig6 | 3igb | 3igp | 3igv | 3ii5 | 3iiw | 3iiy |
| 3ij0 | 3ijh | 3ijy | 3ijz | 3ik1 | 3ik3 | 3ikc | 3ikd | 3ikg | 3il6 |
| 3imc | 3ime | 3img | 3imy | 3in4 | 3inf | 3inh | 3iny | 3iob | 3ioc |
| 3iod | 3ioe | 3iok | 3ion | 3iop | 3ip5 | 3ip6 | 3ip9 | 3ipa | 3ipb |
| 3ipe | 3ipq | 3ips | 3ipy | 3iqj | 3iqu | 3iqv | 3is9 | 3isj | 3iss |
| 3ith | 3itu | 3itz | 3iub | 3iuc | 3iue | 3ivc | 3ivh | 3ivi | 3ivq |
| 3ivv | 3ivx | 3iw4 | 3iw5 | 3iw6 | 3iw7 | 3iw8 | 3jdw | 3jpv | 3jq7 |
| 3jqa | 3jqf | 3jsi | 3jsw | 3juk | 3jvk | 3jwq | 3jwr | 3jxw | 3jy0 |
| 3jy9 | 3jyj | 3jzb | 3jzc | 3jzf | 3jzg | 3jzh | 3jzi | 3jzk | 3k00 |
| 3k05 | 3k0h | 3k0k | 3k15 | 3k16 | 3k1j | 3k22 | 3k23 | 3k27 | 3k2f |
| 3k37 | 3k39 | 3k3a | 3k3b | 3k3g | 3k3h | 3k3j | 3k41 | 3k4d | 3k54 |
| 3k5c | 3k5d | 3k5f | 3k5i | 3k5k | 3k5u | 3k5x | 3k8c | 3k8d | 3k8o |
| 3k8q | 3k97 | 3k98 | 3k99 | 3kab | 3kac | 3kad | 3kaf | 3kag | 3kah |
| 3kai | 3kb7 | 3kbz | 3kc0 | 3kc1 | 3kc3 | 3kce | 3kcf | 3kd7 | 3kdc |
| 3kdd | 3kdm | 3kdt | 3kek | 3ken | 3kf7 | 3kfa | 3kfc | 3kgq | 3kgt |
| 3kgu | 3khj | 3khv | 3kig | 3kjd | 3kjn | 3kjq | 3kku | 3kl8 | 3kmm |
| 3kmx | 3kmy | 3kn0 | 3koo | 3kpu | 3kpv | 3kpw | 3kqm | 3kqo | 3kqp |
| 3kqt | 3kqw | 3kqy | 3kr0 | 3kr1 | 3kr2 | 3kr4 | 3kr5 | 3krj | 3krl |
| 3krr | 3krw | 3krx | 3kvw | 3kwf | 3kwj | 3kx1 | 3kxz | 3kyr | 3kze |
| 3l08 | 3l0k | 3l0n | 3l13 | 3l1n | 3l1s | 3l38 | 3l3a | 3l3l | 3l3m |
| 3l3n | 3l4t | 3l4u | 3l4v | 3l4w | 3l4x | 3l4y | 3l4z | 3l54 | 3l58 |
| 3l59 | 3l5b | 3l5e | 3l5f | 3l6h | 3l79 | 3l7a | 3l7c | 3l7d | 3l8s |
| 3l8v | 3l8x | 3l9h | 3l9l | 3l9m | 3l9n | 3lau | 3lbj | 3lbk | 3lbl |
| 3lbz | 3lc3 | 3lc5 | 3lcd | 3lco | 3lcu | 3lcv | 3ldp | 3ldq | 3ldw |
| 3le8 | 3lf0 | 3lfn | 3lfs | 3lgs | 3lhg | 3lhj | 3lik | 3lil | 3lir |
| 3liw | 3lj3 | 3ljg | 3lk0 | 3lk1 | 3lkh | 3lkz | 3lmk | 3lmp | 3lnk |
| 3loo | 3lpb | 3lpg | 3lpj | 3lpk | 3lpp | 3lq5 | 3ls4 | 3luo | 3lvp |
| 3lvw | 3lw0 | 3lxg | 3lxk | 3lxl | 3lxo | 3ly2 | 3lzb | 3lzs | 3lzu |
| 3m11 | 3m2u | 3m3x | 3m3z | 3m40 | 3m5e | 3m67 | 3m6f | 3m8p | 3m8u |
| 3m96 | 3m9f | 3ma3 | 3mam | 3mb6 | 3mb7 | 3mct | 3mdz | 3mf5 | 3mfw |
| 3mhc | 3mhi | 3mhl | 3mhm | 3mho | 3mhw | 3mi2 | 3mi3 | 3mj1 | 3mj2 |
| 3mjl | 3mkn | 3mks | 3ml2 | 3ml5 | 3mlb | 3mmf | 3mna | 3mo2 | 3mo5 |
| 3mof | 3moh | 3mp1 | 3mp6 | 3mpt | 3mqf | 3mrt | 3mrv | 3mrx | 3ms2 |
| 3ms4 | 3ms7 | 3ms9 | 3msc | 3msj | 3msk | 3msl | 3mt7 | 3mt8 | 3mt9 |
| 3mta | 3mtb | 3mtd | 3muf | 3muk | 3muz | 3mv5 | 3mvj | 3mvl | 3mvm |
| 3mw1 | 3mwe | 3mwu | 3mww | 3mxc | 3mxd | 3mxe | 3mxf | 3mxy | 3my1 |
| 3my5 | 3myq | 3mz3 | 3mzc | 3n0h | 3n0n | 3n1c | 3n1v | 3n1w | 3n2e |
| 3n2p | 3n35 | 3n3g | 3n3j | 3n3l | 3n49 | 3n4b | 3n4l | 3n51 | 3n5h |
| 3n5j | 3n5k | 3n7r | 3n87 | 3n8k | 3n8n | 3n9r | 3nal | 3nam | 3nb5 |
| 3nc4 | 3nc9 | 3ncg | 3ncz | 3ndm | 3nee | 3nef | 3neg | 3neo | 3nes |
| 3new | 3nex | 3nf6 | 3nf7 | 3nf9 | 3nfl | 3ng4 | 3nga | 3nhi | 3nht |
| 3ni5 | 3nif | 3nim | 3nin | 3nlb | 3nm6 | 3nmq | 3nnu | 3nnv | 3nnw |
| 3nnx | 3nox | 3np7 | 3np9 | 3npa | 3nq3 | 3nrm | 3nrz | 3ns9 | 3nsh |
| 3nsq | 3ntp | 3nu3 | 3nu4 | 3nu5 | 3nu6 | 3nu9 | 3nuj | 3nuo | 3nus |
| 3nuu | 3nuy | 3nw3 | 3nw5 | 3nw6 | 3nw7 | 3nww | 3nxq | 3nyd | 3nyn |
| 3nyx | 3nzc | 3nzs | 3nzu | 3o2m | 3o4k | 3o56 | 3o57 | 3o5x | 3o75 |
| 3o84 | 3o8g | 3o8h | 3o8p | 3o95 | 3o96 | 3o99 | 3o9a | 3o9b | 3o9c |
| 3o9d | 3o9e | 3o9f | 3o9g | 3o9p | 3oad | 3oag | 3oaw | 3oay | 3ob0 |
| 3obq | 3obx | 3oc0 | 3ocg | 3ocp | 3odk | 3odu | 3oe6 | 3oe9 | 3ogm |
| 3ogp | 3ogq | 3ohf | 3ohh | 3ohi | 3oik | 3oil | 3oim | 3ok9 | 3oka |
| 3okh | 3oki | 3okp | 3oku | 3okv | 3olf | 3omm | 3oob | 3oof | 3ook |
| 3oot | 3ooz | 3opm | 3oq5 | 3oqf | 3oqk | 3osi | 3osw | 3ot3 | 3ot8 |
| 3otf | 3otq | 3ov1 | 3ove | 3ovz | 3ow3 | 3ow6 | 3owb | 3owd | 3owj |
| 3owk | 3owl | 3oxi | 3oy0 | 3oy1 | 3oy3 | 3oyl | 3oyn | 3oyq | 3oys |
| 3ozj | 3ozp | 3ozr | 3p0g | 3p17 | 3p1d | 3p23 | 3p2e | 3p2h | 3p3g |
| 3p3r | 3p3t | 3p3u | 3p44 | 3p4q | 3p4r | 3p4v | 3p4w | 3p50 | 3p55 |
| 3p58 | 3p5k | 3p79 | 3p7a | 3p7c | 3p7i | 3p8h | 3p8n | 3p8o | 3p8z |
| 3p9j | 3pa3 | 3pa4 | 3pa5 | 3pax | 3pb3 | 3pb9 | 3pbb | 3pd8 | 3pd9 |
| 3pdc | 3pe1 | 3pe2 | 3peq | 3pgu | 3phe | 3pi5 | 3pix | 3pj1 | 3pj8 |
| 3pjc | 3pjt | 3pju | 3pm1 | 3po1 | 3poa | 3poz | 3pp0 | 3pp1 | 3pp7 |
| 3ppj | 3ppk | 3ppm | 3ppo | 3ppp | 3ppq | 3ppr | 3pqz | 3prf | 3prz |
| 3ps1 | 3ps6 | 3psb | 3psl | 3pty | 3pvu | 3pvw | 3pwd | 3pwh | 3pwk |
| 3pxe | 3pxy | 3pxz | 3py0 | 3py1 | 3q0z | 3q1x | 3q2j | 3q32 | 3q3b |
| 3q3t | 3q43 | 3q44 | 3q4j | 3q6k | 3q6w | 3q6z | 3q71 | 3q7j | 3q8d |
| 3q92 | 3q96 | 3qa2 | 3qaa | 3qai | 3qak | 3qaq | 3qar | 3qbh | 3qbn |
| 3qc4 | 3qc9 | 3qce | 3qcf | 3qch | 3qci | 3qcj | 3qcl | 3qcq | 3qcs |
| 3qcx | 3qcy | 3qd0 | 3qdd | 3qel | 3qem | 3qfv | 3qfy | 3qfz | 3qg6 |
| 3qgw | 3qi1 | 3qi3 | 3qi4 | 3qiy | 3qiz | 3qj0 | 3qj9 | 3qk0 | 3qk5 |
| 3qkd | 3qkl | 3qnd | 3qo3 | 3qo9 | 3qox | 3qpn | 3qpo | 3qpp | 3qps |
| 3qqa | 3qqk | 3qqu | 3qrk | 3qs1 | 3qs4 | 3qs5 | 3qs6 | 3qs8 | 3qsb |
| 3qt6 | 3qt7 | 3qtf | 3qti | 3qtq | 3qtr | 3qts | 3qtu | 3qtw | 3qtx |
| 3qtz | 3qu0 | 3que | 3qup | 3qvu | 3qvv | 3qx9 | 3qxm | 3qxp | 3qzt |
| 3qzv | 3r00 | 3r01 | 3r02 | 3r0t | 3r0w | 3r0y | 3r16 | 3r17 | 3r1v |
| 3r21 | 3r22 | 3r2a | 3r2f | 3r2y | 3r42 | 3r4m | 3r4n | 3r4o | 3r4p |
| 3r5j | 3r5n | 3r5t | 3r6c | 3r6g | 3r6u | 3r7b | 3r7n | 3r7o | 3r7q |
| 3r7r | 3r8i | 3r8u | 3r8v | 3r8z | 3r91 | 3r92 | 3r9d | 3r9n | 3rah |
| 3rak | 3ral | 3rcd | 3rcj | 3rde | 3rdv | 3rey | 3rhx | 3ri1 | 3rik |
| 3ril | 3rin | 3rjc | 3rjw | 3rk7 | 3rk9 | 3rl7 | 3rl8 | 3rlp | 3rlq |
| 3rm4 | 3rm8 | 3rme | 3rmf | 3rni | 3ro4 | 3roc | 3rpr | 3rpv | 3rpy |
| 3rq7 | 3rse | 3rsr | 3rsv | 3rt6 | 3rt8 | 3rtf | 3rth | 3rti | 3rtm |
| 3rtn | 3rtp | 3ru1 | 3rup | 3rv3 | 3rv4 | 3rv6 | 3rv7 | 3rv8 | 3rv9 |
| 3rvg | 3rw9 | 3rwp | 3rwq | 3rx5 | 3rx7 | 3rx8 | 3rxe | 3ry8 | 3ryv |
| 3ryx | 3ryy | 3ryz | 3rz0 | 3rz1 | 3rz3 | 3rz5 | 3rz7 | 3rz8 | 3rzb |
| 3rzi | 3s00 | 3s0b | 3s0d | 3s0e | 3s0j | 3s0o | 3s1g | 3s1h | 3s1y |
| 3s2a | 3s2o | 3s2p | 3s2v | 3s3i | 3s3k | 3s3q | 3s3r | 3s3v | 3s43 |
| 3s45 | 3s4q | 3s53 | 3s54 | 3s5y | 3s6t | 3s71 | 3s72 | 3s73 | 3s74 |
| 3s75 | 3s76 | 3s77 | 3s78 | 3s7a | 3s7b | 3s7f | 3s7l | 3s7m | 3s8l |
| 3s8n | 3s8o | 3s9e | 3s9t | 3sap | 3sax | 3saz | 3sbh | 3sbi | 3sc1 |
| 3sd5 | 3sfc | 3sff | 3sfg | 3sfh | 3sfi | 3sgx | 3sh0 | 3sh1 | 3sha |
| 3shc | 3shv | 3shy | 3shz | 3sie | 3sio | 3sk2 | 3ska | 3ske | 3skf |
| 3skg | 3skh | 3sl4 | 3sl5 | 3sl8 | 3sm0 | 3sm2 | 3smq | 3sn7 | 3sni |
| 3snl | 3so9 | 3soq | 3sov | 3spf | 3spk | 3sqq | 3sr4 | 3srb | 3src |
| 3srv | 3st5 | 3st6 | 3su0 | 3su1 | 3su2 | 3su3 | 3su4 | 3su5 | 3sud |
| 3sue | 3suf | 3sug | 3sur | 3sus | 3sut | 3suu | 3suv | 3suw | 3sv2 |
| 3sw9 | 3sww | 3sx4 | 3sx9 | 3sxf | 3sxu | 3sym | 3sz1 | 3szm | 3t01 |
| 3t03 | 3t07 | 3t0t | 3t19 | 3t1a | 3t1l | 3t1m | 3t1n | 3t2q | 3t2t |
| 3t2w | 3t3c | 3t3d | 3t3e | 3t3g | 3t3h | 3t3i | 3t3u | 3t3v | 3t4n |
| 3t5i | 3t5u | 3t60 | 3t64 | 3t6b | 3t6j | 3t6y | 3t70 | 3t82 | 3t83 |
| 3t84 | 3t85 | 3tao | 3tay | 3tb6 | 3tc5 | 3tcg | 3tcp | 3tct | 3tcy |
| 3td4 | 3tdh | 3tdj | 3te5 | 3tf6 | 3tfu | 3tfv | 3tge | 3tgs | 3thb |
| 3thd | 3ti1 | 3ti3 | 3ti4 | 3ti5 | 3ti6 | 3ti8 | 3tia | 3tib | 3tic |
| 3tif | 3tiy | 3tjc | 3tkh | 3tki | 3tkm | 3tku | 3tkw | 3tkz | 3tl5 |
| 3tll | 3tn8 | 3tne | 3tnh | 3tpp | 3tpr | 3ts4 | 3tt0 | 3tti | 3ttj |
| 3ttm | 3ttn | 3ttp | 3ttz | 3tu1 | 3tu7 | 3tu9 | 3tv4 | 3tv5 | 3tv6 |
| 3tv7 | 3tv8 | 3tvw | 3tvx | 3twd | 3twj | 3txo | 3ty0 | 3tyv | 3tz0 |
| 3tz2 | 3tz4 | 3tza | 3u10 | 3u15 | 3u2k | 3u3u | 3u3z | 3u4o | 3u4u |
| 3u4w | 3u5l | 3u6a | 3u6h | 3u6i | 3u6j | 3u6w | 3u78 | 3u7s | 3u81 |
| 3u8j | 3u8l | 3u8m | 3u8w | 3u90 | 3u92 | 3u93 | 3u9c | 3u9n | 3ua8 |
| 3ua9 | 3ubd | 3ubx | 3udd | 3udj | 3udk | 3udm | 3udn | 3udp | 3udq |
| 3udr | 3udy | 3uec | 3ug2 | 3uh4 | 3uhm | 3uil | 3uix | 3uj9 | 3ujb |
| 3ujc | 3ujd | 3ukr | 3uli | 3umo | 3ump | 3umq | 3umw | 3unj | 3unn |
| 3unz | 3uo5 | 3uod | 3uoh | 3uok | 3uol | 3uph | 3upi | 3upx | 3upz |
| 3uqf | 3uqg | 3ur0 | 3usx | 3uu1 | 3uug | 3uvl | 3uvn | 3uvp | 3uvq |
| 3uvx | 3uwl | 3uwo | 3ux0 | 3uxm | 3uyt | 3uz5 | 3uza | 3uzc | 3uzj |
| 3uzp | 3v01 | 3v0l | 3v0p | 3v3l | 3v3m | 3v3v | 3v49 | 3v51 | 3v5g |
| 3v5j | 3v5l | 3v5p | 3v5q | 3v5t | 3v78 | 3v7c | 3v7x | 3v8s | 3v8t |
| 3v8w | 3vbg | 3vbq | 3vbt | 3vbv | 3vbw | 3vbx | 3vby | 3vc4 | 3vd7 |
| 3veh | 3veu | 3vf3 | 3vf5 | 3vf8 | 3vf9 | 3vfa | 3vfb | 3vfq | 3vg1 |
| 3vha | 3vhc | 3vhe | 3vhk | 3vi2 | 3vi5 | 3vi7 | 3vid | 3vjc | 3vje |
| 3vnt | 3vo3 | 3voz | 3vp1 | 3vp2 | 3vp3 | 3vp4 | 3vqh | 3vqu | 3vrt |
| 3vru | 3vrv | 3vrw | 3vry | 3vs3 | 3vsw | 3vsx | 3vtc | 3vtd | 3vtr |
| 3vv6 | 3vv7 | 3vv8 | 3vva | 3vvy | 3vvz | 3vw0 | 3vw6 | 3vw7 | 3vws |
| 3vx3 | 3vyd | 3vye | 3vzd | 3vzg | 3vzv | 3w0l | 3w1f | 3w2o | 3w2r |
| 3w2s | 3w32 | 3w33 | 3w54 | 3w55 | 3w5e | 3w5n | 3w9r | 3wab | 3wav |
| 3waw | 3wb4 | 3wb5 | 3wbl | 3wc7 | 3wd1 | 3wd2 | 3wd9 | 3wf5 | 3wf6 |
| 3wf7 | 3wf9 | 3wff | 3wfg | 3wgw | 3wha | 3wi2 | 3wig | 3wix | 3wiy |
| 3wiz | 3wjw | 3wk4 | 3wk5 | 3wk6 | 3wk7 | 3wk8 | 3wk9 | 3wkb | 3wkc |
| 3wkd | 3wke | 3wmb | 3wpn | 3wq6 | 3wqh | 3wqm | 3ws8 | 3ws9 | 3wt5 |
| 3wt7 | 3wtn | 3wyk | 3wym | 3wyx | 3wyy | 3wz6 | 3wz7 | 3wzj | 3wzk |
| 3zbf | 3zbx | 3zc5 | 3zc6 | 3zcl | 3zcw | 3zdh | 3zep | 3zhf | 3zhz |
| 3zi0 | 3zi8 | 3zj8 | 3zk6 | 3zki | 3zlk | 3zll | 3zln | 3zlo | 3zlq |
| 3zlr | 3zls | 3zlx | 3zly | 3zm4 | 3zm5 | 3zm6 | 3zm9 | 3zmg | 3zmm |
| 3zmq | 3zmt | 3zmu | 3zmv | 3zmz | 3zn0 | 3zn1 | 3znr | 3zo1 | 3zo2 |
| 3zo4 | 3zos | 3zov | 3zpq | 3zpr | 3zpt | 3zpu | 3zqe | 3zqt | 3zrk |
| 3zrl | 3zrm | 3zs0 | 3zs1 | 3zsq | 3zst | 3zsw | 3zsy | 3zsz | 3zt1 |
| 3zt3 | 3zt4 | 3ztc | 3ztd | 3ztx | 3zv7 | 3zvv | 3zw3 | 3zxe | 3zxz |
| 3zya | 3zyb | 3zyu | 3zze | 3zzf | 3zzh | 4a16 | 4a22 | 4a23 | 4a4l |
| 4a4o | 4a4q | 4a4v | 4a4w | 4a4x | 4a50 | 4a6v | 4a6w | 4a7b | 4a7c |
| 4a9i | 4a9m | 4a9n | 4a9r | 4a9s | 4a9t | 4a9u | 4aa0 | 4aa4 | 4aa5 |
| 4aac | 4aaw | 4abf | 4abk | 4ac3 | 4acc | 4acd | 4acg | 4ach | 4aci |
| 4acm | 4acu | 4acx | 4ael | 4af3 | 4afe | 4afg | 4afh | 4afj | 4aft |
| 4ag8 | 4agc | 4agd | 4agm | 4ago | 4ah9 | 4ahu | 4ahv | 4ai5 | 4aia |
| 4aif | 4aj1 | 4aj2 | 4aj4 | 4aje | 4aji | 4ajk | 4ajl | 4ajn | 4ajo |
| 4akn | 4al4 | 4alg | 4alu | 4alv | 4alw | 4alx | 4amw | 4an2 | 4an3 |
| 4an9 | 4anb | 4anm | 4anu | 4anw | 4aoi | 4ap7 | 4apo | 4app | 45751 |
| 4aq3 | 4aq4 | 4aqc | 4ara | 4arb | 4ark | 4arw | 4as9 | 4asd | 4ase |
| 4asj | 4asy | 4at3 | 4at4 | 4at5 | 4au8 | 4aua | 4avu | 4avw | 4aw5 |
| 4awi | 4awj | 4axa | 4axd | 4ay5 | 4ay6 | 4ayt | 4ayv | 4ayw | 4ayx |
| 4az2 | 4az5 | 4az6 | 4azb | 4azc | 4aze | 4azf | 4azg | 4azi | 4azy |
| 4b00 | 4b05 | 4b0b | 4b0c | 4b0g | 4b0j | 4b11 | 4b12 | 4b13 | 4b14 |
| 4b1c | 4b1d | 4b1j | 4b2d | 4b2i | 4b2l | 4b32 | 4b33 | 4b34 | 4b35 |
| 4b3b | 4b3c | 4b3d | 4b3u | 4b4g | 4b4m | 4b5b | 4b5d | 4b6c | 4b6e |
| 4b6f | 4b6o | 4b6p | 4b6q | 4b6r | 4b6s | 4b70 | 4b71 | 4b72 | 4b73 |
| 4b74 | 4b76 | 4b77 | 4b78 | 4b7j | 4b7n | 4b7p | 4b7q | 4b7r | 4b7z |
| 4b82 | 4b83 | 4b84 | 4b85 | 4b8p | 4b9k | 4ba3 | 4bak | 4bam | 4ban |
| 4baq | 4bb2 | 4bb4 | 4bb9 | 4bbe | 4bbf | 4bbg | 4bbh | 4bc5 | 4bcd |
| 4bcf | 4bcg | 4bch | 4bci | 4bcj | 4bck | 4bcm | 4bcn | 4bco | 4bcp |
| 4bcq | 4bcs | 4bcw | 4bda | 4bdb | 4bdd | 4bde | 4bdf | 4bdg | 4bdh |
| 4bdi | 4bdj | 4bdk | 4bds | 4bdt | 4bek | 4bf1 | 4bf6 | 4bfd | 4bfp |
| 4bfr | 4bfy | 4bfz | 4bg6 | 4bgg | 4bgh | 4bgm | 4bgx | 4bgy | 4bh3 |
| 4bh4 | 4bhn | 4bhz | 4bi0 | 4bi1 | 4bi2 | 4bi7 | 4bib | 4bic | 4bid |
| 4bie | 4bio | 4bj8 | 4bj9 | 4bjb | 4bjc | 4bjx | 4bkj | 4bks | 4bkz |
| 4blb | 4bnt | 4bnu | 4bnv | 4bnx | 4bny | 4bnz | 4bo0 | 4bo1 | 4bo3 |
| 4bo4 | 4bo5 | 4bo7 | 4bo8 | 4bo9 | 4bqg | 4bqh | 4bqs | 4bqt | 4brx |
| 4bs0 | 4bs4 | 4bt9 | 4btb | 4btk | 4btl | 4btm | 4btw | 4btx | 4bty |
| 4bup | 4bvb | 4bw1 | 4bw2 | 4bw3 | 4bw4 | 4bxk | 4byi | 4bzn | 4bzo |
| 4bzr | 4bzs | 4c1d | 4c1e | 4c1m | 4c1t | 4c1u | 4c1w | 4c1y | 4c2v |
| 4c35 | 4c36 | 4c37 | 4c38 | 4c4e | 4c4f | 4c4g | 4c4h | 4c4i | 4c4j |
| 4c52 | 4c5d | 4c61 | 4c66 | 4c68 | 4c6z | 4c70 | 4c72 | 4c73 | 4c7t |
| 4c94 | 4c9w | 4c9x | 4ca5 | 4ca8 | 4cc5 | 4cc6 | 4cc7 | 4cd0 | 4cd1 |
| 4cd5 | 4cdr | 4ce1 | 4ce2 | 4ce3 | 4ceb | 4cfl | 4cfm | 4cft | 4cfu |
| 4cfv | 4cfw | 4cfx | 4cgi | 4cgj | 4ch8 | 4ci1 | 4ci2 | 4ci3 | 4cix |
| 4ciz | 4cj4 | 4cjn | 4cjp | 4cjq | 4cjr | 4ck3 | 4cki | 4ckj | 4ckr |
| 4cku | 4cl9 | 4clb | 4cli | 4clj | 4clp | 4clz | 4cmo | 4cmt | 4cmu |
| 4cnh | 4cp7 | 4cps | 4cpt | 4cpx | 4cpy | 4cpz | 4cqf | 4cqg | 4cr5 |
| 4crd | 4csd | 4csj | 4ctj | 4ctk | 4cts | 4cu1 | 4cu7 | 4cu8 | 4cwf |
| 4cwn | 4cwo | 4cwp | 4cwq | 4cwr | 4cws | 4cwt | 4cxw | 4cxx | 4d0w |
| 4d0x | 4d1a | 4d1b | 4d1c | 4d1d | 4d1j | 4d1s | 4d1y | 4d2d | 4d2p |
| 4d2r | 4d2s | 4d2t | 4d2v | 4d2w | 4d83 | 4d85 | 4d88 | 4d89 | 4d8a |
| 4d8c | 4d8s | 4d8z | 4da5 | 4daf | 4db7 | 4dbm | 4dbn | 4dce | 4dcs |
| 4dcv | 4ddl | 4ddm | 4dds | 4ddy | 4de0 | 4de5 | 4de7 | 4dea | 4deg |
| 4deh | 4dei | 4del | 4der | 4des | 4det | 4deu | 4dew | 4dff | 4dfg |
| 4dfu | 4dgb | 4dgg | 4dgm | 4dgn | 4dgr | 4dij | 4dj7 | 4djo | 4djp |
| 4djq | 4djr | 4dju | 4djw | 4djx | 4djy | 4dk5 | 4dk7 | 4dlj | 4dma |
| 4dmn | 4dmw | 4do3 | 4do4 | 4do5 | 4dpf | 4dpi | 4dpt | 4dpu | 4dpy |
| 4dq2 | 4drk | 4drm | 4drn | 4dru | 4dst | 4dsu | 4dtk | 4dtt | 4du8 |
| 4duh | 4dus | 4dve | 4dwk | 4dxg | 4dy6 | 4dzw | 4dzy | 4e0w | 4e0x |
| 4e1e | 4e1k | 4e20 | 4e26 | 4e28 | 4e35 | 4e3d | 4e3f | 4e3g | 4e3h |
| 4e49 | 4e4a | 4e4l | 4e4n | 4e4x | 4e5d | 4e6c | 4e6d | 4e70 | 4e7r |
| 4e8w | 4e8y | 4e8z | 4e96 | 4e9u | 4ea1 | 4ebv | 4ebw | 4ec0 | 4edy |
| 4edz | 4ee0 | 4eeh | 4eev | 4ef4 | 4ef6 | 4efk | 4eft | 4efu | 4eg4 |
| 4eg5 | 4eg6 | 4eg7 | 4ega | 4egh | 4egi | 4egk | 4eh2 | 4eh3 | 4eh4 |
| 4eh5 | 4eh6 | 4eh7 | 4eh9 | 4ehe | 4ehg | 4ehv | 4ehz | 4ei4 | 4ej8 |
| 4ejl | 4ejn | 4ek9 | 4eke | 4ekg | 4elb | 4ele | 4elf | 4elg | 4elh |
| 4em7 | 4emt | 4en4 | 4enx | 4eny | 4eo4 | 4eo6 | 4eoh | 4eoi | 4eok |
| 4eol | 4eon | 4eop | 4eos | 4ep2 | 4epy | 4eqc | 4ere | 4erk | 4erw |
| 4etz | 4eu0 | 4eu3 | 4euc | 4euo | 4euv | 4ewh | 4ewn | 4exg | 4exh |
| 4eym | 4ez3 | 4ez5 | 4ezj | 4ezk | 4ezl | 4ezw | 4ezx | 4ezy | 4ezz |
| 4f08 | 4f1l | 4f1s | 4f3k | 4f5y | 4f63 | 4f64 | 4f6s | 4f6u | 4f6w |
| 4f70 | 4f7j | 4f7n | 4f8h | 4f8j | 4f9g | 4f9u | 4f9y | 4fab | 4fai |
| 4fbe | 4fbx | 4fc0 | 4fcb | 4fcd | 4fcq | 4fcr | 4fem | 4feq | 4few |
| 4ff8 | 4fgz | 4fhh | 4fhi | 4fht | 4fic | 4fjz | 4fkk | 4fl1 | 4flh |
| 4flp | 4fm7 | 4fm8 | 4fmn | 4fmo | 4fmq | 4fn5 | 4fns | 4fny | 4fnz |
| 4fob | 4foc | 4fod | 4fpf | 4fpk | 4fr3 | 4fri | 4frj | 4frk | 4frs |
| 4fs3 | 4fs4 | 4fse | 4fsl | 4fut | 4fxf | 4fxj | 4fxp | 4fxq | 4fxz |
| 4fys | 4fz3 | 4fz6 | 4fzj | 4g0l | 4g0y | 4g11 | 4g16 | 4g17 | 4g19 |
| 4g1f | 4g2f | 4g2j | 4g2l | 4g2r | 4g2w | 4g2y | 4g31 | 4g34 | 4g3f |
| 4g3g | 4g4p | 4g55 | 4g5f | 4g5y | 4g69 | 4g8m | 4g8n | 4g8r | 4g8v |
| 4g8y | 4g90 | 4g9c | 4gah | 4gao | 4gb9 | 4gbd | 4gby | 4gbz | 4gcj |
| 4ge1 | 4ge2 | 4ge4 | 4ge7 | 4gfd | 4gfo | 4gg5 | 4gg7 | 4ggl | 4ggz |
| 4gh6 | 4ghi | 4gih | 4giu | 4gj2 | 4gj3 | 4gj6 | 4gj8 | 4gj9 | 4gja |
| 4gjb | 4gjc | 4gjd | 4gk2 | 4gk3 | 4gk4 | 4gkh | 4gki | 4glw | 4glx |
| 4gm3 | 4gm8 | 4gmc | 4gmy | 4gne | 4gny | 4gpk | 4gq4 | 4gq6 | 4gql |
| 4gqq | 4gqr | 4gs8 | 4gs9 | 4gsc | 4gsy | 4gts | 4gtv | 4gu6 | 4gue |
| 4gui | 4guj | 4gvm | 4gw1 | 4gw5 | 4gw6 | 4gw8 | 4gwi | 4gwk | 4gxl |
| 4gxs | 4gz3 | 4gzp | 4gzt | 4gzw | 4gzx | 4h1m | 4h36 | 4h39 | 4h3b |
| 4h3i | 4h3q | 4h42 | 4h4b | 4h4d | 4h4e | 4h4m | 4h58 | 4h5e | 4h71 |
| 4h75 | 4h7q | 4h81 | 4h85 | 4ha5 | 4hai | 4hbm | 4hbn | 4hbv | 4hbw |
| 4hby | 4hco | 4hcz | 4hdb | 4hdc | 4hdf | 4hdp | 4he9 | 4heg | 4hej |
| 4heu | 4hf4 | 4hfz | 4hg7 | 4hgs | 4hgt | 4hhy | 4hhz | 4hiq | 4his |
| 4hj2 | 4hki | 4hkk | 4hkn | 4hkp | 4hl5 | 4hla | 4hlc | 4hld | 4hlf |
| 4hlg | 4hlh | 4hlk | 4hlm | 4hlw | 4hmh | 4hmk | 4hnf | 4hni | 4hnn |
| 4hod | 4hp0 | 4hpi | 4hra | 4hs8 | 4hso | 4ht0 | 4ht2 | 4ht6 | 4htp |
| 4htx | 4hu1 | 4hva | 4hvd | 4hvg | 4hvh | 4hvi | 4hvs | 4hw2 | 4hw3 |
| 4hw7 | 4hwb | 4hwp | 4hwr | 4hxj | 4hxl | 4hxm | 4hxr | 4hxs | 4hxw |
| 4hxz | 4hy1 | 4hy9 | 4hyb | 4hyf | 4hyh | 4hyi | 4hym | 4hys | 4hyu |
| 4hzm | 4hzt | 4hzw | 4hzx | 4hzz | 4i0d | 4i0f | 4i0s | 4i0t | 4i0z |
| 4i10 | 4i11 | 4i12 | 4i1c | 4i47 | 4i4e | 4i4f | 4i54 | 4i5c | 4i5h |
| 4i5m | 4i5p | 4i6b | 4i6f | 4i6h | 4i72 | 4i7f | 4i7j | 4i7k | 4i7l |
| 4i7m | 4i7p | 4i80 | 4i8n | 4i8w | 4i8x | 4i8z | 4i9c | 4i9h | 4i9i |
| 4i9u | 4i9z | 4ib5 | 4ibb | 4ibc | 4ibd | 4ibe | 4ibf | 4ibg | 4ibi |
| 4ibj | 4ibk | 4ibm | 4idn | 4ido | 4idt | 4idv | 4idz | 4ie0 | 4ie4 |
| 4ie6 | 4ie7 | 4ifh | 4ifi | 4igk | 4igr | 4igt | 4iho | 4iic | 4iid |
| 4iie | 4iif | 4ij1 | 4ijh | 4ijl | 4ijp | 4ikn | 4ikr | 4ikt | 4im0 |
| 4inb | 4io2 | 4io3 | 4io4 | 4io5 | 4io6 | 4io7 | 4io8 | 4ipf | 4ipn |
| 4iq6 | 4iqt | 4iqu | 4irx | 4is6 | 4isu | 4ith | 4iti | 4itj | 4iue |
| 4iuo | 4iur | 4iut | 4iuu | 4iuv | 4iva | 4ivk | 4ivs | 4ivt | 4iwd |
| 4iz0 | 4izm | 4izy | 4j04 | 4j06 | 4j08 | 4j09 | 4j0a | 4j0p | 4j0r |
| 4j0s | 4j0t | 4j0v | 4j0y | 4j0z | 4j17 | 4j1c | 4j1e | 4j1f | 4j1h |
| 4j1i | 4j1k | 4j22 | 4j26 | 4j2c | 4j2t | 4j3d | 4j3e | 4j3j | 4j3m |
| 4j4n | 4j4o | 4j4v | 4j51 | 4j52 | 4j53 | 4j6i | 4j73 | 4j74 | 4j77 |
| 4j78 | 4j79 | 4j7d | 4j7e | 4j7i | 4j81 | 4j82 | 4j84 | 4j86 | 4j8b |
| 4j8g | 4j8m | 4j8r | 4j93 | 4jaj | 4jal | 4jaz | 4jbo | 4jbp | 4jck |
| 4jfd | 4jfe | 4jff | 4jfi | 4jfj | 4jfk | 4jfl | 4jfm | 4jft | 4jfx |
| 4jfz | 4jg0 | 4jg6 | 4jh0 | 4jhz | 4jib | 4jik | 4jin | 4jj7 | 4jje |
| 4jjm | 4jjs | 4jju | 4jkt | 4jkw | 4jlg | 4jlh | 4jlj | 4jlm | 4jln |
| 4jls | 4jmg | 4jmu | 4jn4 | 4jnc | 4jnj | 4jnm | 4joe | 4jof | 4jog |
| 4joh | 4joj | 4jok | 4joo | 4jp9 | 4jpe | 4jps | 4jpx | 4jpy | 4jq7 |
| 4jq8 | 4jql | 4jr0 | 4jr3 | 4jr5 | 4jrg | 4jrv | 4jsa | 4jsc | 4jsr |
| 4jt8 | 4jt9 | 4ju3 | 4ju4 | 4ju6 | 4ju7 | 4jv6 | 4jv7 | 4jv8 | 4jv9 |
| 4jvb | 4jvq | 4jx7 | 4jx9 | 4jxv | 4jxw | 4jyb | 4jyc | 4jym | 4k0o |
| 4k0y | 4k1b | 4k2f | 4k2g | 4k3h | 4k3k | 4k3l | 4k3m | 4k3n | 4k3o |
| 4k3p | 4k3q | 4k3r | 4k42 | 4k43 | 4k4f | 4k4j | 4k55 | 4k5l | 4k5n |
| 4k5o | 4k5y | 4k63 | 4k64 | 4k66 | 4k67 | 4k6i | 4k6y | 4k6z | 4k72 |
| 4k76 | 4k7i | 4k7n | 4k7o | 4k8a | 4k8o | 4k8s | 4k9h | 4k9y | 4kab |
| 4kai | 4kb8 | 4kb9 | 4kbc | 4kbi | 4kbk | 4kcg | 4ke0 | 4ke1 | 4keq |
| 4kfp | 4kin | 4kip | 4kiq | 4kiu | 4kiw | 4klb | 4klv | 4km0 | 4km2 |
| 4kmd | 4kmu | 4kn0 | 4kn1 | 4knb | 4kni | 4knj | 4knn | 4knr | 4knx |
| 4kod | 4kom | 4kon | 4kot | 4kov | 4kow | 4kox | 4kp5 | 4kp6 | 4kp8 |
| 4kpx | 4kpz | 4kql | 4kqp | 4kqq | 4kqr | 4krs | 4ks1 | 4ks2 | 4ks3 |
| 4ks4 | 4ks5 | 4ksq | 4ksy | 4ktc | 4kup | 4kva | 4kvm | 4kwf | 4kwg |
| 4kwp | 4kww | 4kxb | 4kxn | 4kz0 | 4kz3 | 4kz4 | 4kz5 | 4kz7 | 4kz8 |
| 4kza | 4kzb | 4kzc | 4kzl | 4l02 | 4l09 | 4l0b | 4l0i | 4l0s | 4l0t |
| 4l0v | 4l10 | 4l1a | 4l1u | 4l23 | 4l2f | 4l2g | 4l2k | 4l2l | 4l31 |
| 4l32 | 4l33 | 4l34 | 4l3p | 4l4m | 4l4v | 4l4z | 4l50 | 4l51 | 4l52 |
| 4l53 | 4l5j | 4l6s | 4l70 | 4l7b | 4l7c | 4l7d | 4l7f | 4l7g | 4l7h |
| 4l7j | 4l7l | 4l7n | 4l7o | 4l7r | 4l7u | 4l8m | 4l9i | 4la7 | 4lbl |
| 4lbo | 4lbu | 4lch | 4led | 4leq | 4lgg | 4lgh | 4lh2 | 4lh3 | 4lh6 |
| 4lh7 | 4lhm | 4li6 | 4lj5 | 4lj8 | 4lkg | 4lkh | 4lkj | 4lkk | 4lko |
| 4ll3 | 4llj | 4llk | 4llp | 4lm0 | 4lm1 | 4lm2 | 4lm3 | 4lm4 | 4lm5 |
| 4lmn | 4lmu | 4ln2 | 4lnb | 4lnf | 4lng | 4lno | 4lnp | 4lnw | 4loh |
| 4loi | 4loo | 4lop | 4loq | 4loy | 4lp0 | 4lp9 | 4lpb | 4lpg | 4lph |
| 4lq3 | 4lq9 | 4lrr | 4lsj | 4lte | 4lts | 4luo | 4luv | 4luz | 4lv4 |
| 4lvt | 4lw1 | 4lwc | 4lwe | 4lwh | 4lwt | 4lwu | 4lwv | 4lww | 4lxb |
| 4lxd | 4ly9 | 4lys | 4lyw | 4lzr | 4m0e | 4m0f | 4m0r | 4m1d | 4m2r |
| 4m2v | 4m2w | 4m3d | 4m3e | 4m3g | 4m3m | 4m3q | 4m48 | 4m5g | 4m5i |
| 4m5j | 4m5k | 4m5l | 4m6p | 4m6q | 4m7j | 4m84 | 4m8e | 4m8h | 4m8x |
| 4m8y | 4man | 4mbc | 4mbi | 4mbj | 4mbl | 4mc1 | 4mc2 | 4mc6 | 4mc9 |
| 4mcb | 4mcc | 4mcd | 4mcv | 4md6 | 4mdn | 4mdq | 4mdr | 4mds | 4mdt |
| 4men | 4meo | 4mep | 4meq | 4mf0 | 4mf1 | 4mg6 | 4mg7 | 4mg8 | 4mg9 |
| 4mga | 4mgb | 4mgc | 4mgv | 4mh7 | 4mha | 4mho | 4mhs | 4mhy | 4mhz |
| 4mi3 | 4mi6 | 4mi9 | 4mib | 4mic | 4mji | 4mjo | 4mjp | 4mjq | 4mjr |
| 4mk0 | 4mk7 | 4mk8 | 4mk9 | 4mka | 4mlx | 4mm4 | 4mm5 | 4mm6 | 4mm7 |
| 4mm8 | 4mm9 | 4mma | 4mmf | 4mmm | 4mmp | 4mnp | 4mo4 | 4mo8 | 4mot |
| 4mp2 | 4mp7 | 4mpc | 4mpn | 4mq1 | 4mq2 | 4mq6 | 4mqp | 4mqu | 4mr3 |
| 4mr4 | 4mr5 | 4mr6 | 4mra | 4mre | 4mrf | 4mrg | 4mrh | 4mro | 4mrw |
| 4mrz | 4ms0 | 4msa | 4msc | 4msg | 4msk | 4msl | 4msn | 4msu | 4mt9 |
| 4mti | 4mue | 4muf | 4muk | 4mul | 4muw | 4mvh | 4mvw | 4mvx | 4mvy |
| 4mw0 | 4mw1 | 4mw2 | 4mw4 | 4mw5 | 4mw6 | 4mw7 | 4mw9 | 4mwc | 4mwe |
| 4mwq | 4mwr | 4mwu | 4mwv | 4mww | 4mwx | 4mwy | 4mx0 | 4mx9 | 4mxa |
| 4mxc | 4myd | 4myh | 4myq | 4mz4 | 4mz5 | 4mz6 | 4mzh | 4n00 | 4n07 |
| 4n1b | 4n1u | 4n3r | 4n3w | 4n4t | 4n4v | 4n5d | 4n6g | 4n6y | 4n6z |
| 4n70 | 4n7e | 4n7h | 4n7j | 4n7m | 4n8d | 4n8e | 4n8q | 4n98 | 4n99 |
| 4n9a | 4n9b | 4n9c | 4n9d | 4n9e | 4na4 | 4na7 | 4na8 | 4nah | 4nat |
| 4nb6 | 4nbk | 4ncg | 4ncm | 4ncn | 4nct | 4ndu | 4ngm | 4ngn | 4nh7 |
| 4nh8 | 4nh9 | 4nhx | 4nie | 4nj3 | 4njd | 4nka | 4nks | 4nku | 4nl1 |
| 4nld | 4nmo | 4nmp | 4nmq | 4nmr | 4nms | 4nmt | 4nmv | 4nnr | 4np3 |
| 4np9 | 4nra | 4nrb | 4nrc | 4nrk | 4nrl | 4nrm | 4nrt | 4nru | 4ntj |
| 4nuc | 4nud | 4nue | 4nus | 4nvp | 4nw5 | 4nw6 | 4nwc | 4nwd | 4nxu |
| 4nxv | 4ny3 | 4nyf | 4nyi | 4nyj | 4nym | 4o04 | 4o05 | 4o07 | 4o09 |
| 4o0a | 4o0b | 4o0r | 4o0t | 4o0v | 4o0x | 4o0y | 4o0z | 4o10 | 4o12 |
| 4o13 | 4o15 | 4o1b | 4o1d | 4o1l | 4o24 | 4o28 | 4o2a | 4o2b | 4o2c |
| 4o2e | 4o2f | 4o2p | 4o37 | 4o3a | 4o3b | 4o3c | 4o3f | 4o42 | 4o43 |
| 4o44 | 4o45 | 4o4r | 4o4y | 4o5g | 4o6e | 4o70 | 4o71 | 4o72 | 4o74 |
| 4o75 | 4o76 | 4o77 | 4o7a | 4o7b | 4o7c | 4o7e | 4o7f | 4o97 | 4o9s |
| 4oag | 4oar | 4obo | 4obp | 4obq | 4oc0 | 4oc2 | 4ocq | 4ocx | 4ocz |
| 4od0 | 4od7 | 4odf | 4oew | 4oex | 4og3 | 4og4 | 4og5 | 4og6 | 4og7 |
| 4og8 | 4ogi | 4ogn | 4ogt | 4ohk | 4ohm | 4ohp | 4ojq | 4ojr | 4ok3 |
| 4ok5 | 4ok6 | 4okg | 4okp | 4oks | 4olc | 4old | 4olh | 4oma | 4omj |
| 4omk | 4ona | 4onf | 4ono | 4oow | 4op1 | 4op2 | 4op3 | 4oq5 | 4oq6 |
| 4or0 | 4otg | 4oth | 4oty | 4oue | 4ov5 | 4ovf | 4ovg | 4ovh | 4ow0 |
| 4own | 4owo | 4owv | 4oya | 4oyb | 4oyi | 4oyk | 4oym | 4oyo | 4oyp |
| 4oys | 4oyt | 4oz1 | 4oz2 | 4oz3 | 4ozo | 4p00 | 4p02 | 4p0a | 4p0n |
| 4p0v | 4p0x | 4p1r | 4p1u | 4p2t | 4p3h | 4p45 | 4p4b | 4p4i | 4p4s |
| 4p4t | 4p58 | 4p5d | 4p5e | 4p5z | 4p6e | 4p6g | 4p6w | 4p6x | 4p72 |
| 4p73 | 4p74 | 4p75 | 4p7e | 4pax | 4pb1 | 4pb2 | 4pce | 4pci | 4pct |
| 4pd5 | 4pd6 | 4pd7 | 4pd8 | 4pd9 | 4pda | 4pde | 4pee | 4pf3 | 4pft |
| 4pfu | 4pg3 | 4ph4 | 4phu | 4phv | 4phw | 4pin | 4pio | 4pjt | 4pkt |
| 4pku | 4pkw | 4pl0 | 4pm0 | 4pml | 4pmm | 4pmp | 4pms | 4pmt | 4pn1 |
| 4pni | 4pnk | 4pnl | 4pnm | 4pnn | 4pnq | 4pnr | 4pns | 4pnt | 4pnu |
| 4pnw | 4po0 | 4poh | 4poj | 4pow | 4pox | 4pp0 | 4pp3 | 4pp5 | 4pp9 |
| 4ppa | 4ppb | 4ppc | 4pqn | 4pra | 4prg | 4pri | 4prj | 4prn | 4ps3 |
| 4ps5 | 4ps7 | 4ps8 | 4psb | 4psh | 4psq | 4psx | 4pte | 4ptg | 4puj |
| 4puk | 4pul | 4pum | 4puz | 4pv0 | 4pv7 | 4pvo | 4pvt | 4pvv | 4pxf |
| 4py1 | 4py2 | 4pyn | 4pyo | 4pyq | 4pyv | 4pyx | 4pyy | 4pz8 | 4pzh |
| 4pzv | 4pzw | 4pzx | 4q08 | 4q0a | 4q0k | 4q15 | 4q18 | 4q19 | 4q1e |
| 4q1f | 4q1n | 4q4e | 4q4o | 4q4p | 4q4q | 4q4r | 4q6d | 4q6e | 4q6r |
| 4q7p | 4q7s | 4q7v | 4q7w | 4q81 | 4q83 | 4q87 | 4q8x | 4q8y | 4q90 |
| 4q93 | 4q99 | 4q9m | 4q9o | 4q9s | 4q9y | 4q9z | 4qaa | 4qab | 4qag |
| 4qdk | 4qfg | 4qfr | 4qfs | 4qg7 | 4qga | 4qge | 4qgf | 4qgg | 4qgh |
| 4qht | 4qij | 4qiz | 4qj0 | 4qjm | 4qjp | 4qjx | 4ql8 | 4qmm | 4qmn |
| 4qmo | 4qmp | 4qmq | 4qms | 4qmt | 4qmu | 4qmw | 4qmx | 4qmy | 4qmz |
| 4qn7 | 4qna | 4qnu | 4qo4 | 4qp1 | 4qp9 | 4qpd | 4qq5 | 4qr4 | 4qrc |
| 4qsh | 4qsm | 4qsu | 4qsv | 4qsw | 4qsx | 4qt0 | 4qta | 4qtb | 4qtc |
| 4qtl | 4qtn | 4qxo | 4qxq | 4qxr | 4qxs | 4qye | 4qyg | 4qyh | 4qyy |
| 4r06 | 4r0a | 4r3c | 4r3w | 4r4c | 4r4i | 4r4o | 4r4q | 4r4t | 4r5a |
| 4r5b | 4r5n | 4r5t | 4r5v | 4r5x | 4r6e | 4r6t | 4r6w | 4r6x | 4r74 |
| 4r75 | 4r76 | 4r7m | 4r8y | 4r91 | 4r92 | 4r93 | 4r95 | 4ra1 | 4ra5 |
| 4rak | 4rcd | 4rce | 4rcf | 4rcg | 4rd6 | 4rdn | 4re2 | 4re4 | 4re9 |
| 4res | 4ret | 4rfc | 4rfd | 4rfy | 4rfz | 4rg0 | 4rio | 4riu | 4riv |
| 4rj3 | 4rj4 | 4rj5 | 4rj6 | 4rj7 | 4rj8 | 4rlk | 4rll | 4rn4 | 4rpn |
| 4rpo | 4rqk | 4rqv | 4rrn | 4rro | 4rrs | 4rrv | 4rs0 | 4rsc | 4rse |
| 4rsk | 4rss | 4rux | 4ruy | 4ruz | 4rvk | 4rvl | 4rvm | 4rvt | 4rx7 |
| 4rx8 | 4rx9 | 4ryl | 4tim | 4tju | 4tjw | 4tjy | 4tk0 | 4tk1 | 4tk2 |
| 4tk3 | 4tk4 | 4tk5 | 4tkf | 4tkg | 4tki | 4tln | 4tlr | 4tmp | 4tmr |
| 4tn2 | 4tn4 | 4tpm | 4tpp | 4tpt | 4tpw | 4ts1 | 4tt2 | 4tte | 4tu4 |
| 4tv3 | 4tvj | 4tw6 | 4tw7 | 4tw9 | 4twc | 4twd | 4tww | 4tx6 | 4txs |
| 4ty6 | 4ty8 | 4ty9 | 4tya | 4tyb | 4tyl | 4tyo | 4tz2 | 4tz8 | 4tzm |
| 4tzn | 4u0d | 4u0e | 4u0f | 4u0i | 4u0m | 4u0n | 4u0u | 4u2y | 4u3f |
| 4u43 | 4u44 | 4u45 | 4u4x | 4u54 | 4u58 | 4u5j | 4u5o | 4u5s | 4u5u |
| 4u5v | 4u69 | 4u6c | 4u6e | 4u6r | 4u6w | 4u6y | 4u71 | 4u73 | 4u79 |
| 4u7o | 4u8w | 4u90 | 4u91 | 4u93 | 4ua8 | 4uac | 4ual | 4uc5 | 4uco |
| 4ucr | 4ucs | 4uct | 4ucu | 4ucv | 4uda | 4udb | 4ufe | 4uff | 4ufg |
| 4ufh | 4ufi | 4ufj | 4ufk | 4ufl | 4ufm | 4ufz | 4uix | 4uiz | 4uj1 |
| 4uj2 | 4uj9 | 4uja | 4ujb | 4uma | 4umb | 4umc | 4umq | 4umr | 4umt |
| 4umu | 4und | 4urk | 4urn | 4uru | 4urv | 4urw | 4urx | 4ury | 4urz |
| 4us3 | 4us4 | 4usj | 4usw | 4utn | 4utr | 4utv | 4utx | 4uu7 | 4uu8 |
| 4uua | 4uub | 4uv8 | 4uv9 | 4uva | 4uvb | 4uvc | 4uwf | 4uwg | 4uwh |
| 4uwl | 4ux6 | 4uxb | 4uxl | 4uxq | 4uyd | 4uye | 4uyf | 4uyg | 4uyh |
| 4uyn | 4uzd | 4uzh | 4v05 | 4v24 | 4v25 | 4w4v | 4w4w | 4w4x | 4w4y |
| 4w4z | 4w50 | 4w52 | 4w53 | 4w54 | 4w55 | 4w57 | 4w7p | 4w7t | 4w97 |
| 4w9d | 4w9e | 4w9f | 4w9j | 4w9k | 4w9n | 4w9o | 4w9p | 4w9q | 4w9w |
| 4wa9 | 4waf | 4wag | 4wbo | 4wf2 | 4wf6 | 4wgi | 4wh7 | 4wh9 | 4whq |
| 4whr | 4whs | 4wht | 4why | 4whz | 4wi1 | 4wj5 | 4wj7 | 4wke | 4wki |
| 4wmu | 4wmv | 4wmx | 4wmy | 4wnk | 4wov | 4wp7 | 4wpn | 4wsy | 4wt6 |
| 4wuy | 4wvl | 4ww6 | 4wwn | 4wwo | 4wwp | 4wy1 | 4wy3 | 4wy6 | 4wym |
| 4wz8 | 4x11 | 4x1f | 4x2l | 4x3r | 4x47 | 4x48 | 4x49 | 4x5y | 4x5z |
| 4x60 | 4x63 | 4x6k | 4x6m | 4x6n | 4x6x | 4x6y | 4x7h | 4x7i | 4x7j |
| 4x7k | 4x7l | 4x7n | 4x7o | 4x7q | 4xaq | 4xar | 4xas | 4xe0 | 4xe1 |
| 4xg6 | 4xg7 | 4xg8 | 4xg9 | 4xh6 | 4xhl | 4xip | 4xiq | 4xir | 4xj0 |
| 4xkx | 4xm6 | 4xm7 | 4xm8 | 4xmo | 4xnv | 4xnw | 4xqu | 4xs2 | 4xta |
| 4xtv | 4xtx | 4xty | 4xtz | 4xu0 | 4xu1 | 4xu2 | 4xu3 | 4xuh | 4xum |
| 4xwk | 4xx9 | 4xxs | 4xy9 | 4xyc | 4xyf | 4y2b | 4y2j | 4y2p | 4y2q |
| 4y2s | 4y2t | 4y2u | 4y2v | 4y2x | 4y2y | 4y46 | 4y4v | 4y5h | 4y62 |
| 4y64 | 4y6m | 4y73 | 4y83 | 4y85 | 4y87 | 4y8c | 4y8d | 4y8x | 4y8y |
| 4y8z | 4ybj | 4ybk | 4ycl | 4ycm | 4ycn | 4ycu | 4ycv | 4ydq | 4yh3 |
| 4yht | 4yjn | 4yk0 | 4yll | 4ylu | 4ymj | 4ynd | 4yo6 | 4yog | 4yoi |
| 4yoj | 4yp8 | 4ypf | 4yqh | 4yrc | 4yrd | 4yrg | 4yrr | 4ytc | 4yth |
| 4yti | 4yur | 4yuw | 4yux | 4yuy | 4yuz | 4yv0 | 4yv1 | 4yvc | 4yve |
| 4yx4 | 4yxd | 4yxi | 4yxo | 4yxu | 4yyt | 4yz5 | 4yz9 | 4yzc | 4z1j |
| 4z1k | 4z1n | 4z22 | 4z2h | 4z2i | 4z2j | 4z2k | 4z83 | 4z84 | 4z8d |
| 4z93 | 4z9l | 4za0 | 4zed | 4zei | 4zek | 4zg9 | 4zim | 4zk5 | 4zki |
| 4zl4 | 4zla | 4zlo | 4zme | 4zmf | 4zow | 4zpe | 4zpf | 4zpg | 4zqt |
| 4zs0 | 4zs2 | 4zs3 | 4zsm | 4zsp | 4zsq | 4zsr | 4zt2 | 4zt4 | 4zt5 |
| 4zt6 | 4zt7 | 4ztl | 4ztn | 4ztq | 4ztr | 4zts | 4zw3 | 4zw5 | 4zw7 |
| 4zw8 | 4zx0 | 4zx1 | 4zx4 | 4zx5 | 4zx8 | 4zx9 | 4zy0 | 4zy1 | 4zy2 |
| 4zyr | 4zzx | 4zzy | 5a00 | 5a14 | 5a3n | 5a3t | 5a3w | 5a3x | 5a4e |
| 5a4l | 5a4q | 5a54 | 5a5s | 5a5z | 5a69 | 5a6a | 5a6b | 5a6h | 5a6k |
| 5a6n | 5a7y | 5a9u | 5aa8 | 5aa9 | 5aaa | 5aab | 5aac | 5aad | 5aae |
| 5aaf | 5aag | 5ab9 | 5abe | 5abf | 5abg | 5abp | 5acw | 5ael | 5aep |
| 5ai0 | 5ai4 | 5ai6 | 5ai8 | 5ai9 | 5aia | 5aib | 5aic | 5ajv | 5ajw |
| 5ajx | 5ajy | 5ajz | 5ak0 | 5ak2 | 5ak3 | 5ak4 | 5ak5 | 5ak6 | 5ake |
| 5akg | 5akh | 5aki | 5akj | 5akk | 5akl | 5akz | 5ald | 5ale | 5alf |
| 5alg | 5alh | 5ali | 5alj | 5alk | 5all | 5alm | 5aln | 5alo | 5alp |
| 5alr | 5als | 5alt | 5alu | 5alv | 5alw | 5alx | 5aly | 5am0 | 5am1 |
| 5am2 | 5am3 | 5am4 | 5am5 | 5amd | 5amg | 5aml | 5amn | 5anq | 5aom |
| 5ap0 | 5ap1 | 5ap2 | 5ap3 | 5ap4 | 5ap5 | 5ap6 | 5aqf | 5aqg | 5aqh |
| 5aqj | 5aqk | 5aqn | 5aqo | 5aqp | 5aqq | 5aqr | 5aqt | 5aqu | 5aqz |
| 5ar5 | 5ar7 | 5arf | 5arg | 5auu | 5auv | 5auw | 5aux | 5auy | 5auz |
| 5av0 | 5avi | 5ax9 | 5axq | 5ayy | 5b0x | 5b1s | 5b25 | 5b4k | 5b4l |
| 5bml | 5bms | 5bnm | 5bnr | 5bns | 5bpp | 5bqs | 5bue | 5bve | 5bvf |
| 5bvk | 5bvn | 5bvo | 5bvw | 5bw4 | 5byi | 5c1w | 5c26 | 5c42 | 5c4k |
| 5c5h | 5c6o | 5c6p | 5c8k | 5c8m | 5c8n | 5cal | 5can | 5cao | 5cap |
| 5caq | 5cas | 5cau | 5cav | 5cbm | 5cbr | 5cbs | 5cc2 | 5ccl | 5cdh |
| 5ceh | 5cf4 | 5cf8 | 5cgc | 5cgd | 5cgj | 5cj6 | 5cjf | 5ckr | 5cks |
| 5clm | 5cnj | 5cnm | 5cpr | 5cqu | 5cr7 | 5cs6 | 5csh | 5csp | 5ct7 |
| 5ctc | 5cte | 5cu2 | 5cu4 | 5cuq | 5cwa | 5cxh | 5cy3 | 5czb | 5d0r |
| 5d12 | 5d1n | 5d24 | 5d26 | 5d3l | 5d3n | 5d3p | 5d3s | 5d3t | 5d6j |
| 5d7a | 5da3 | 5db0 | 5db1 | 5db2 | 5db3 | 5dbm | 5dd9 | 5dda | 5ddb |
| 5ddc | 5ddd | 5dde | 5ddf | 5dgz | 5dh3 | 5dh5 | 5dhg | 5dhh | 5dhj |
| 5dhp | 5dhq | 5dhr | 5dhs | 5dht | 5dhu | 5diq | 5dit | 5diu | 5div |
| 5djr | 5dk4 | 5dnu | 5doh | 5dpx | 5dqc | 5dqf | 5dri | 5dro | 5drq |
| 5dry | 5dt2 | 5dtj | 5dtk | 5dtq | 5dtr | 5dts | 5dtt | 5dv2 | 5dva |
| 5dx4 | 5dxh | 5dxt | 5dxu | 5dy5 | 5dyt | 5dyw | 5dyy | 5e1e | 5e1s |
| 5e28 | 5e2k | 5e2l | 5e2n | 5e2r | 5e2s | 5e5g | 5e8r | 5e91 | 5ea3 |
| 5ea5 | 5ea7 | 5eak | 5ech | 5eci | 5ecv | 5edq | 5edr | 5eds | 5edu |
| 5ee7 | 5eek | 5efh | 5egs | 5egu | 5eh0 | 5eh5 | 5eh7 | 5ehe | 5ehg |
| 5ehi | 5ehn | 5ehp | 5ehr | 5ehv | 5ehw | 5ehy | 5ei2 | 5ei6 | 5ei8 |
| 5eif | 5eij | 5eiw | 5ejv | 5ekh | 5ekj | 5ekm | 5ekn | 5eko | 5ekx |
| 5elv | 5em5 | 5em6 | 5em7 | 5em8 | 5eng | 5enk | 5enm | 5ep7 | 5eqy |
| 5er1 | 5er2 | 5erg | 5es1 | 5etj | 5eud | 5eue | 5evb | 5evk | 5evz |
| 5ew3 | 5ew9 | 5ewj | 5ewm | 5exn | 5exw | 5ey0 | 5ey4 | 5ey8 | 5ey9 |
| 5eyd | 5eyk | 5eym | 5ezx | 5f00 | 5f01 | 5f1x | 5f20 | 5f2k | 5f2r |
| 5f2w | 5f32 | 5f3t | 5f3z | 5f41 | 5f4n | 5f5i | 5f60 | 5f94 | 5f95 |
| 5f9b | 5fcw | 5fcz | 5fd2 | 5fdc | 5fdi | 5fdo | 5fdp | 5fhm | 5fhn |
| 5fi2 | 5fi6 | 5fi7 | 5fkj | 5fky | 5fl0 | 5fl1 | 5flo | 5flp | 5flq |
| 5flt | 5fnc | 5fnd | 5fnf | 5fng | 5fnj | 5fnq | 5fnr | 5fns | 5fnt |
| 5fnu | 5fp0 | 5fqc | 5fqp | 5fqr | 5fqs | 5fqt | 5fqv | 5fsx | 5fsy |
| 5fto | 5fue | 5fv7 | 5fwa | 5fwj | 5fxq | 5fxr | 5g1n | 5g1p | 5g2b |
| 5g2n | 5g4n | 5g53 | 5g57 | 5ggz | 5ghv | 5gja | 5gjf | 5gjg | 5gmh |
| 5gmn | 5gn5 | 5gn6 | 5gs9 | 5gsa | 5gut | 5gv2 | 5gvk | 5gvl | 5gvm |
| 5gvp | 5h09 | 5h0b | 5h0e | 5h0g | 5h0h | 5h13 | 5h19 | 5h22 | 5h2u |
| 5h3q | 5h85 | 5h8b | 5h8e | 5h8g | 5ha9 | 5hbh | 5hct | 5hcv | 5hcx |
| 5hcy | 5hd0 | 5hdv | 5hdx | 5hdz | 5he0 | 5he1 | 5he2 | 5he4 | 5he7 |
| 5hes | 5hex | 5hfu | 5hg1 | 5hgq | 5hh5 | 5hh6 | 5his | 5hjq | 5hkm |
| 5hln | 5hlp | 5hls | 5hlw | 5hm0 | 5hmy | 5hn0 | 5hn7 | 5hn8 | 5hn9 |
| 5hna | 5hng | 5ho6 | 5ho7 | 5ho8 | 5hoa | 5hor | 5hu0 | 5hvp | 5hvu |
| 5hvy | 5hx6 | 5hx8 | 5hzn | 5i0b | 5i2r | 5i3v | 5i3w | 5i3x | 5i3y |
| 5i80 | 5i83 | 5i88 | 5i89 | 5i8b | 5i8g | 5i8p | 5i94 | 5i9i | 5i9x |
| 5i9y | 5ia0 | 5ia1 | 5ia3 | 5ia4 | 5ia5 | 5idp | 5ie1 | 5iee | 5ief |
| 5ifu | 5ih5 | 5ih6 | 5ih8 | 5ih9 | 5iha | 5ij7 | 5im3 | 5in9 | 5ism |
| 5itd | 5ito | 5itp | 5iu4 | 5iu7 | 5iu8 | 5iua | 5iub | 5iuh | 5iv4 |
| 5ivf | 5ivj | 5izk | 5izl | 5izm | 5j0d | 5j1v | 5j20 | 5j27 | 5j2x |
| 5j4n | 5j4y | 5j58 | 5j59 | 5j5r | 5j64 | 5j6a | 5j6d | 5j6l | 5j6m |
| 5j6n | 5j71 | 5j75 | 5j7b | 5j7w | 5j82 | 5j86 | 5j8m | 5j8u | 5j8z |
| 5j9l | 5j9x | 5jah | 5jal | 45662 | 5jao | 5jap | 5jar | 5jas | 5jat |
| 5jau | 5jcb | 5jfr | 5jga | 5jgb | 5jgd | 5jhb | 5jjr | 5jjs | 5jn8 |
| 5jn9 | 5jna | 5jnc | 5jox | 5jpt | 5jq5 | 5jq7 | 5jq8 | 5jq9 | 5jqb |
| 5jur | 5jv0 | 5jv1 | 5jv2 | 5jxn | 5jxq | 5jyp | 5jzn | 5jzy | 5k00 |
| 5k0m | 5k0t | 5k1i | 5k48 | 5k4i | 5k4j | 5k4l | 5k4x | 5k4z | 5k5e |
| 5k5n | 5k5s | 5k76 | 5k8n | 5k8v | 5k9w | 5ka1 | 5ka3 | 5ka7 | 5ka9 |
| 5kab | 5kad | 5kby | 5kcx | 5kdr | 5ke0 | 5khx | 5kit | 5kj2 | 5kjk |
| 5kjm | 5kjn | 5kkr | 5kks | 5kkt | 5kls | 5kmf | 5knj | 5kpk | 5kpl |
| 5kq5 | 5kqf | 5kr8 | 5ksx | 5ktx | 5ku3 | 5ku6 | 5kv8 | 5kv9 | 5kva |
| 5kw2 | 5kww | 5kz0 | 5kzq | 5l13 | 5l2i | 5l2m | 5l2n | 5l2o | 5l2s |
| 5l2t | 5l3a | 5l3e | 5l3j | 5l44 | 5l4h | 5l6h | 5l6j | 5l72 | 5l7e |
| 5l7g | 5l7h | 5l8c | 5l8y | 5l9g | 5l9h | 5l9i | 5l9l | 5l9o | 5lbq |
| 5lce | 5lch | 5ld8 | 5le1 | 5lg3 | 5lgn | 5lgt | 5lhg | 5lhh | 5lhi |
| 5lj1 | 5lj2 | 5ljj | 5lkr | 5ll4 | 5ll5 | 5ll9 | 5lla | 5llc | 5lle |
| 5llg | 5llh | 5llm | 5lm4 | 5lma | 5lmb | 5lmk | 5lo5 | 5lo6 | 5lom |
| 5lp6 | 5lpd | 5lqf | 5lqq | 5ls6 | 5lsc | 5lsg | 5luu | 5lvl | 5lvn |
| 5lwd | 5lwm | 5lwn | 5lxc | 5lyx | 5lyy | 5lz2 | 5lz4 | 5lz5 | 5lz7 |
| 5lz8 | 5lz9 | 5m04 | 5m0d | 5m0m | 5m0s | 5m39 | 5m44 | 5m4c | 5m4f |
| 5m4i | 5m4k | 5m4q | 5m4u | 5m51 | 5m53 | 5m55 | 5m56 | 5m57 | 5m6u |
| 5m7u | 45721 | 5mat | 5mby | 5meh | 5mek | 5mf6 | 5mfr | 5mfs | 5mhp |
| 5mhq | 5mi3 | 5mi5 | 5mi7 | 5mi9 | 5mim | 5mja | 5mjn | 5mks | 5ml5 |
| 5mli | 5mme | 5mmg | 5mo8 | 5mod | 5moe | 5mpk | 5mpn | 5mpz | 5mqe |
| 5mqv | 5mrb | 5mrd | 5mrh | 5mri | 5msb | 5mtx | 5mw3 | 5mxr | 5mz3 |
| 5mz8 | 5n0d | 5n0e | 5n1r | 5n1s | 5n25 | 5n2x | 5n2z | 5n34 | 5n4s |
| 5n4t | 5n53 | 5n55 | 5n58 | 5n7v | 5n84 | 5n87 | 5n93 | 5n9k | 5n9l |
| 5n9n | 5n9s | 5n9t | 5nad | 5nap | 5nau | 5ncy | 5ncz | 5ndf | 5ne5 |
| 5nea | 5nee | 5nev | 5nfh | 5ng9 | 5ngb | 5ngu | 5nhf | 5nhz | 5ni0 |
| 5nih | 5njz | 5nk2 | 5nk3 | 5nk4 | 5nk6 | 5nk7 | 5nk8 | 5nk9 | 5nka |
| 5nkb | 5nkc | 5nkd | 5nkg | 5nkh | 5nki | 5nkk | 5nlk | 5nn0 | 5nn4 |
| 5nn5 | 5nn6 | 5nr8 | 5nra | 5nrf | 5nu1 | 5nuu | 5nw7 | 5nx9 | 5nxg |
| 5nxi | 5nxo | 5nxp | 5nxv | 5nxw | 5ny1 | 5ny3 | 5ny6 | 5nya | 5nze |
| 5nzf | 5nzm | 5nzn | 5nzo | 5nzp | 5nzq | 5o07 | 5o0j | 5o11 | 5o1d |
| 5o1h | 5o2d | 5o5m | 5o7e | 5o83 | 5o91 | 5o9h | 5o9o | 5o9p | 5o9q |
| 5o9r | 5oa2 | 5obj | 5obr | 5oci | 5oei | 5of0 | 5ofu | 5ofv | 5ofw |
| 5ogk | 5ogl | 5oht | 5ohy | 5oku | 5one | 5op6 | 5op8 | 5oqu | 5orh |
| 5ork | 5orr | 5ort | 5orv | 5orw | 5orx | 5ory | 5orz | 5os0 | 5os1 |
| 5os2 | 5os3 | 5os4 | 5os5 | 5os7 | 5os8 | 5osd | 5ose | 5osk | 5osl |
| 5ot8 | 5ot9 | 5ota | 5otc | 5otr | 5otz | 5ou2 | 5ou3 | 5oul | 5ove |
| 5ovf | 5ovg | 5ovh | 5ovr | 5ovx | 5owh | 5owl | 5oxg | 5prc | 5pzm |
| 5q0g | 5q0h | 5qa4 | 5qa5 | 5qa6 | 5qa7 | 5qa8 | 5qa9 | 5qaa | 5qab |
| 5qac | 5qad | 5qae | 5qag | 5qah | 5qai | 5qaj | 5qak | 5qal | 5qam |
| 5qan | 5qao | 5qap | 5qaq | 5qar | 5qas | 5qat | 5qau | 5qav | 5qaw |
| 5qax | 5qay | 5qaz | 5qb0 | 5qb1 | 5qb2 | 5qb3 | 5qck | 5qcl | 5qin |
| 5qj3 | 5qqo | 5qqp | 5std | 5svk | 5svl | 5swg | 5sz0 | 5sz1 | 5sz2 |
| 5sz3 | 5sz4 | 5sz6 | 5sz7 | 5t19 | 5t1s | 5t1t | 5t1u | 5t1w | 5t23 |
| 5t27 | 5t28 | 5t2b | 5t2d | 5t2g | 5t2i | 5t2l | 5t2m | 5t31 | 5t4b |
| 5t4f | 5t4h | 5t68 | 5t8e | 5t8j | 5t8p | 5t8q | 5t92 | 5t97 | 5tb6 |
| 5tbe | 5tbj | 5tc0 | 5tci | 5tcj | 5tco | 5td2 | 5tex | 5tfx | 5tgc |
| 5thi | 5thj | 5thn | 5ti0 | 5ti2 | 5ti3 | 5ti4 | 5ti5 | 5ti6 | 5tiu |
| 5tjx | 5tkd | 5tks | 5tln | 5toe | 5tol | 5tpg | 5tq3 | 5tq4 | 5tq5 |
| 5tq6 | 5tq8 | 5tqu | 5tr6 | 5trh | 5trk | 5tt7 | 5ttf | 5tuq | 5tur |
| 5tuy | 5tw2 | 5tw3 | 5tw5 | 5twl | 5twz | 5tx5 | 5txy | 5ty1 | 5ty8 |
| 5ty9 | 5tya | 5tz3 | 5tzw | 5tzx | 5tzy | 5u0d | 5u0e | 5u0f | 5u0g |
| 5u0w | 5u0y | 5u0z | 5u11 | 5u12 | 5u13 | 5u14 | 5u2c | 5u2e | 5u2f |
| 5u4x | 5u5h | 5u5k | 5u62 | 5u6b | 5u6c | 5u6d | 5u7j | 5u7l | 5u7o |
| 5u8a | 5u8f | 5uah | 5ubt | 5ueu | 5uey | 5uga | 5ugb | 5ugh | 5ugm |
| 5uig | 5uiq | 5uir | 5uis | 5uit | 5uiu | 5ukj | 5ukk | 5ukl | 5ukm |
| 5ul5 | 5ulg | 5uln | 5ulp | 5un1 | 5uoo | 5uor | 5uox | 5up3 | 5upe |
| 5upf | 5upj | 5uqv | 5urj | 5urm | 5usf | 5usy | 5uuu | 5ux4 | 5uxm |
| 5uxn | 5uyu | 5uzj | 5v0n | 5v19 | 5v24 | 5v2l | 5v3h | 5v3o | 5v3y |
| 5v40 | 5v41 | 5v49 | 5v6u | 5v79 | 5v7a | 5v7i | 5v8q | 5v9p | 5v9t |
| 5vb6 | 5vb7 | 5vcv | 5vcw | 5vcx | 5vcy | 5vcz | 5vd1 | 5vd3 | 5vdo |
| 5vdp | 5vdq | 5vdu | 5vdv | 5vdw | 5vee | 5vex | 5vgy | 5vil | 5vio |
| 5vja | 5vlr | 5vo6 | 5vqe | 5vqq | 5vqs | 5vqt | 5vqu | 5vqw | 5vrl |
| 5vs6 | 5vsc | 5vse | 5vsj | 5vt4 | 5w0e | 5w0f | 5w0i | 5w0l | 5w0q |
| 5w1e | 5w2p | 5w2q | 5w5o | 5w5v | 5w84 | 5w8h | 5w8j | 5w99 | 5wa5 |
| 5wal | 5wb6 | 5wbf | 5wbm | 5wbp | 5wbq | 5wbr | 5wdj | 5wev | 5wex |
| 5wf5 | 5wf6 | 5wf7 | 5wfc | 5wfd | 5wg4 | 5wg5 | 5wg6 | 5wgp | 5wh5 |
| 5wh6 | 5wi0 | 5wj6 | 5wl0 | 5wlt | 5wlv | 5wmg | 5wo4 | 5wp5 | 5wqa |
| 5wqc | 5wr7 | 5ws3 | 5wuk | 5wvd | 5wyq | 5wyr | 5wyx | 5wyz | 5wzr |
| 5x26 | 5x27 | 5x28 | 5x33 | 5x5o | 5x8i | 5x9h | 5xaf | 5xag | 5xbt |
| 5xg4 | 5xhr | 5xig | 5xih | 5xij | 5xiw | 5xkm | 5xmp | 5xmr | 5xms |
| 5xmt | 5xmu | 5xmx | 5xqx | 5xsr | 5xst | 5xsu | 5xv7 | 5xvf | 5xvu |
| 5xyx | 5xyy | 5xzr | 5y1y | 5y2f | 5y5n | 5y5t | 5y5u | 5y6d | 5y6e |
| 5y6k | 5y7j | 5y7k | 5y7z | 5y80 | 5y86 | 5ya5 | 5yas | 5ybi | 5yc8 |
| 5ye7 | 5ye8 | 5ye9 | 5yea | 5yf1 | 5yfs | 5yft | 5yfz | 5yg2 | 5yg3 |
| 5yg4 | 5yh8 | 5yhe | 5yhg | 5yhl | 5yij | 5yjb | 5yjf | 5yji | 5yjo |
| 5yl2 | 5ylj | 5yls | 5ylt | 5ylu | 5ylv | 5yql | 5yqo | 5yve | 5yvt |
| 5ywg | 5ywy | 5yy6 | 5yyb | 5yz7 | 5yzc | 5z1c | 5z1e | 5z4o | 5z5f |
| 5z5v | 5z66 | 5z68 | 5z89 | 5z95 | 5z99 | 5z9e | 5za7 | 5za8 | 5zaf |
| 5zag | 5zc5 | 5zcu | 5ze6 | 5zef | 5zg0 | 5zg1 | 5zg3 | 5zh2 | 5zh3 |
| 5zh4 | 5zh5 | 5zhm | 5zhn | 5zk3 | 5zk8 | 5zkb | 5zkc | 5zlf | 5znc |
| 5zni | 5zo8 | 5zo9 | 5zr3 | 5ztn | 5zty | 5zun | 5zxi | 5zz2 | 6a1c |
| 6a3n | 6a93 | 6a94 | 6aah | 6aaj | 6aak | 6aam | 6aaq | 6abk | 6abp |
| 6abx | 6aej | 6afk | 6agk | 6agt | 6ahi | 6ajg | 6ajh | 6aji | 6ak4 |
| 6ak5 | 6akw | 6alc | 6alz | 6am8 | 6ami | 6an1 | 6anl | 6aol | 6aom |
| 6ap6 | 6ap7 | 6ap8 | 45753 | 6apw | 6aqf | 6aqq | 6aqs | 6arv | 6as6 |
| 6asu | 6aum | 6awo | 6awp | 6axq | 6ay5 | 6b2q | 6b3e | 6b4d | 6b4h |
| 6b59 | 6b5a | 6b5i | 6b7h | 6b8j | 6b8u | 6baw | 6bbu | 6bbv | 6beb |
| 6bed | 6bee | 6beh | 6bfa | 6bfd | 6bfe | 6bfw | 6bfx | 6bh0 | 6bhv |
| 6bky | 6bm6 | 6bmv | 6bmx | 6bny | 6bo6 | 6bod | 6boe | 6boy | 6bqg |
| 6bqh | 6bs5 | 6bsk | 6bu1 | 6bx6 | 6bxy | 6by8 | 6c0n | 6c0r | 6c0s |
| 6c0t | 6c0u | 6c1s | 6c2r | 6c2t | 6c2x | 6c2y | 6c3e | 6c42 | 6c4d |
| 6c4g | 6c5q | 6c6o | 6c7g | 6c7x | 6c8p | 6c91 | 6c9n | 6c9p | 6c9q |
| 6c9r | 6c9s | 6c9v | 6cd5 | 6cf5 | 6cf7 | 6cfc | 6cgt | 6chh | 6cis |
| 6ciy | 6cj5 | 6cje | 6cjh | 6cjw | 6cjy | 6ck6 | 6ckc | 6cki | 6cmj |
| 6cmr | 6cms | 6cn5 | 6cn6 | 6cnj | 6cnk | 6coj | 6cpa | 6cpw | 6cq0 |
| 6cq4 | 6cq5 | 6cqf | 6cse | 6csf | 6csp | 6cvd | 6cyc | 6cyi | 6cz3 |
| 6cz4 | 6czv | 6d1a | 6d1b | 6d1g | 6d1h | 6d1i | 6d28 | 6d4o | 6d4q |
| 6d4u | 6d4v | 6d4w | 6d50 | 6d56 | 6d59 | 6d5e | 6d5h | 6d5j | 6d5w |
| 6d6t | 6d6u | 6d8v | 6dak | 6db3 | 6dcg | 6dd0 | 6dd1 | 6det | 6dgl |
| 6dgo | 6dgr | 6dhu | 6die | 6dih | 6dim | 6djc | 6djd | 6dji | 6djj |
| 6dkb | 6dkg | 6dki | 6dko | 6dlj | 6dne | 6dnp | 6dpx | 6dpz | 6drx |
| 6dry | 6drz | 6dsp | 6dtw | 6dtx | 6duf | 6dug | 6duh | 6dvo | 6dxg |
| 6dy7 | 6e0r | 6e13 | 6e1a | 6e1z | 6e2n | 6e2o | 6e3z | 6e4a | 6e4t |
| 6e4u | 6e4w | 6e59 | 6e7t | 6e7u | 6e7v | 6e8x | 6e99 | 6e9l | 6e9w |
| 6ea1 | 6ea2 | 6eaa | 6eab | 6ebe | 6ebw | 6ecz | 6ed6 | 6eda | 6edl |
| 6eds | 6ee2 | 6ee4 | 6eea | 6eed | 6eeh | 6eeo | 6eg9 | 6ega | 6egs |
| 6ei5 | 6eij | 6eil | 6eip | 6eiq | 6eis | 6ej2 | 6ej3 | 6ej4 | 6ekd |
| 6eks | 6eku | 6el5 | 6eln | 6elo | 6elp | 6emh | 6en4 | 6en5 | 6eo8 |
| 6eo9 | 6eq8 | 6eqm | 6eqp | 6es0 | 6etj | 6euc | 6euv | 6ew3 | 6ewk |
| 6exi | 6exs | 6ey9 | 6eyb | 6ez6 | 6ezg | 6ezh | 6ezq | 6f1j | 6f26 |
| 6f34 | 6f3d | 6f3e | 6f3g | 6f5h | 6f5u | 6f6n | 6f6s | 6f6u | 6f7b |
| 6f7c | 6f7q | 6f86 | 6f8u | 6f90 | 6f9g | 6f9v | 6fba | 6fer | 6few |
| 6fex | 6ffh | 6ffi | 6fh3 | 6fhk | 6fil | 6fnf | 6fni | 6fnj | 6fnq |
| 6fnr | 6fnx | 6fo5 | 6fo7 | 6fo8 | 6fo9 | 6fob | 6fpu | 6fqo | 6fqu |
| 6fr0 | 6fr2 | 6frf | 6fsd | 6fse | 6fsy | 6ft3 | 6ft4 | 6ft8 | 6ft9 |
| 6ftn | 6ftw | 6fu5 | 6fv4 | 6fyv | 6fzg | 6fzj | 6fzu | 6g01 | 6g07 |
| 6g1w | 6g33 | 6g34 | 6g39 | 6g3a | 6g3y | 6g4n | 6g4y | 6g4z | 6g5u |
| 6g6t | 6g6y | 6g6z | 6g7a | 6g93 | 6g9a | 6g9d | 6g9h | 6g9i | 6g9j |
| 6g9k | 6g9m | 6g9n | 6g9x | 6gbw | 6gcw | 6gcx | 6gdg | 6ge0 | 6gg3 |
| 6gg4 | 6gg5 | 6gg8 | 6gge | 6ggg | 6gh9 | 6gi6 | 6gih | 6gin | 6gip |
| 6gjb | 6gl3 | 6gl9 | 6gla | 6glb | 6gmd | 6gn1 | 6got | 6gqm | 6gu3 |
| 6gu4 | 6gu7 | 6guc | 6gue | 6guf | 6guk | 6gva | 6gwr | 6gx3 | 6gxa |
| 6gxb | 6gxe | 6gxq | 6gxu | 6gxw | 6gy0 | 6gz9 | 6gzd | 6gzh | 6gzm |
| 6h12 | 6h13 | 6h14 | 6h29 | 6h2z | 6h33 | 6h34 | 6h36 | 6h37 | 6h38 |
| 6h4d | 6h4u | 6h4z | 6h50 | 6h51 | 6h75 | 6h76 | 6h7d | 6h7f | 6h7j |
| 6h7l | 6h7m | 6h7n | 6h7o | 6h9b | 6h9v | 6h9x | 6hb5 | 6hb6 | 6hb7 |
| 6hbm | 6hbn | 6hcu | 6hcv | 6hcw | 6hgv | 6hh3 | 6hh5 | 6hhr | 6hjk |
| 6hk3 | 6hk4 | 6hk6 | 6hk7 | 6hke | 6hkx | 6hky | 6hlx | 6hlz | 6hm1 |
| 6hm6 | 6hm7 | 6hmb | 6hmx | 6hni | 6hoq | 6hor | 6hot | 6hou | 6hoy |
| 6hp0 | 6hq3 | 6hq4 | 6hq7 | 6hqy | 6hro | 6hrq | 6hs4 | 6hsz | 6ht8 |
| 6htg | 6hth | 6hti | 6htz | 6hu0 | 6hu1 | 6hu2 | 6hv0 | 6hvh | 6hvi |
| 6hvj | 6hys | 6hzp | 6hzu | 6hzv | 6hzy | 6i0b | 6i0z | 6i10 | 6i11 |
| 6i12 | 6i13 | 6i14 | 6i15 | 6i16 | 6i17 | 6i18 | 6i1r | 6i3u | 6i5g |
| 6i5i | 6i5y | 6i8m | 6i8t | 6i8z | 6i9a | 6ibl | 6ibs | 6ibv | 6iby |
| 6ibz | 6ic0 | 6ic2 | 6iik | 6iil | 6iin | 6iiu | 6iiv | 6ikm | 6ilq |
| 6ilz | 6im6 | 6imb | 6imd | 6imi | 6imr | 6ind | 6ipl | 6iql | 6irt |
| 6iup | 6j5l | 6j63 | 6j8r | 6j9w | 6j9y | 6jad | 6jag | 6jam | 45663 |
| 6jao | 6jap | 6jaq | 6jav | 6jaw | 6jb0 | 6jb4 | 6jbb | 6jbe | 6jib |
| 6jid | 6jki | 6jmf | 6jn6 | 6jse | 6jsf | 6jsg | 6jsn | 6jtc | 6jut |
| 6jwa | 6k0j | 6k1q | 6k3l | 6kjd | 6kla | 6kqi | 6kzc | 6kzd | 6m8q |
| 6m95 | 6m9l | 6m9t | 6ma1 | 6ma2 | 6ma3 | 6ma5 | 6mck | 6md0 | 6md4 |
| 6md7 | 6mdb | 6mdc | 6mdd | 6mdq | 6me4 | 6me5 | 6miv | 6miy | 6mjq |
| 6mjw | 6mlf | 6mlw | 6mnv | 6mo4 | 6mod | 6mom | 6moo | 6mq3 | 6mr5 |
| 6msn | 6mso | 6mul | 6mum | 6mvu | 6mvx | 6mx3 | 6mx8 | 6mxc | 6mxd |
| 6mxe | 6myn | 6n0m | 6n17 | 6n19 | 6n3k | 6n3n | 6n3o | 6n3z | 6n48 |
| 6n4b | 6n5f | 6n5g | 6n5h | 6n79 | 6n7a | 6n7c | 6n7d | 6n7y | 6n7z |
| 6n82 | 6n9l | 6nao | 6ndl | 6nfg | 6nfo | 6nfy | 6ng0 | 6nji | 6nmb |
| 6no9 | 6np5 | 6npt | 6nrg | 6nrh | 6nri | 6nrj | 6nsp | 6nss | 6nti |
| 6nu1 | 6nu5 | 6nv7 | 6nv9 | 6ny4 | 6nyh | 6nyv | 6nyw | 6nze | 6nzf |
| 6nzh | 6nzk | 6nzp | 6nzq | 6nzr | 6o0h | 6o1g | 6o2p | 6o4w | 6o50 |
| 6o9d | 6o9x | 6oa1 | 6oag | 6oah | 6ob0 | 6oco | 6ocq | 6ocu | 6od6 |
| 6odz | 6oe1 | 6oe3 | 6oh2 | 6oh3 | 6ohd | 6oi8 | 6oi9 | 6oio | 6oip |
| 6oiq | 6oir | 6oko | 6op9 | 6os6 | 6ox0 | 6oyh | 6oyw | 6oz6 | 6p10 |
| 6p11 | 6p12 | 6p1l | 6p83 | 6peb | 6pf3 | 6pf5 | 6pfj | 6pg3 | 6pg4 |
| 6pg6 | 6pg7 | 6pg8 | 6pga | 6pgb | 6pge | 6pi1 | 6pi5 | 6pi6 | 6pii |
| 6pl2 | 6plf | 6plg | 6pm9 | 6ppy | 6prc | 6ps0 | 6ps1 | 6ps3 | 6ps4 |
| 6ps5 | 6ps6 | 6psb | 6pt3 | 6pyu | 6pz4 | 6q2y | 6q30 | 6q3z | 6q4e |
| 6q54 | 6q60 | 6q6f | 6q73 | 6q89 | 6q8a | 6q8b | 6q8c | 6q8k | 6q8p |
| 6qab | 6qac | 6qad | 6qae | 6qat | 6qau | 6qcj | 6qef | 6ql1 | 6qmd |
| 6qme | 6qmj | 6qmk | 6qs5 | 6qwi | 6qxa | 6qxj | 6qxs | 6qyl | 6qyn |
| 6qyo | 6qz7 | 6qzh | 6r0k | 6r4s | 6r5f | 6r8q | 6r8r | 6rfj | 6rfn |
| 6rj2 | 6rj5 | 6rj6 | 6rk4 | 6rna | 6rot | 6rqk | 6s0e | 6s43 | 6s7k |
| 6s7s | 6s88 | 6s8a | 6s9b | 6s9c | 6s9d | 6sd9 | 6sdc | 6sdd | 6sfc |
| 6sfi | 6sfj | 6sfk | 6slg | 6sq0 | 6sy7 | 6sze | 6szj | 6t6a | 6tim |
| 6tl2 | 6tld | 6u26 | 6u6w | 6u80 | 6u9v | 6ufo | 6ugn | 6ugo | 6ugp |
| 6ugq | 6ugr | 6ugz | 6uh0 | 6uhu | 6uhv | 6uii | 6uil | 6uim | 6ul8 |
| 6upj | 6uvv | 6uvy | 6uwp | 6uwv | 7abp | 7cpa | 7gpb | 7prc | 7std |
| 8a3h | 8abp | 8cpa | 8gpb | 8hvp | 9abp | 9hvp | 9icd |  |  |

Section 12. Protein Crystallization and Structure Determination

To obtain complex structures, the WT FBPase protein (0.2 mg/mL at a final concentration) was incubated with 5 μM 11 for 40 min. And then the sample was concentrated by ultrafiltration to a protein concentration of 8 mg/mL (measured by absorbance at 280 nm). Crystals were obtained using the hanging-drop vapor diffusion method at 18 °C. Crystals grew from a mixture of 1 μL of protein and 1 μL of a well solution containing 1 mM AMP, 0.1 M Tris (pH = 8.4), and 15% v/v EtOH. Crystals were cryoprotected using a well solution supplemented with 40% PEG400 and flash-frozen in liquid nitrogen. X-ray diffraction data were collected at the BL19U1 and BL10U257 beamlines at the Shanghai Synchrotron Radiation Facility. The structure was resolved by molecular replacement using Phaser, and the structure of *Hu-*FBPase (PDB ID: 5ZWK) as the search model. The model was built using Coot and refined with PHENIX. Atomic restraints were generated for the inhibitor using eLBOW, and the model was validated using MolProbity. All structural figures were generated by using PyMOL. The crystal structure has been uploaded to the RCSB PDB, the PDB ID is 9XVI. All related files were uploaded as supplementary materials, see crystalstructure9XVI.zip file.

**Table S10.** Crystallography data collection and refinement statistics for *Hu*FBPase-11t

| **Parameters** | **Hu-FBPase-11t(9XVI)** |
| --- | --- |
| **Data collection** |  |
| Space group | P 21 21 21 |
| Wavelength | 0.979 |
| cell dimensions  a, b, c (Å)  α, β, γ (°) | 67.672 83.261 279.635  90 90 90 |
| resolution (Å) | 48.62 - 1.96 (2.03 - 1.96) |
| *R_merge_* | 0.1714 (0.8765) |
| *I/σ* (I) | 5.75 (1.99) |
| completeness (%) | 99.78 (99.76) |
| redundancy | 13.1(12.5) |
| **Refinement** |  |
| no. reflections | 114172 (11277) |
| *R_work_/R_free_* | 0.1852/0.2133 |
| no. atoms  protein  ligand/ion  water  all atom | 9822  199  863  10884 |
| B-factors  protein  ligand/ion  water  all atom | 26.09  32.00  34.17  26.84 |
| rms deviations |  |
| bond lengths (Å) | 0.008 |
| bond angles (°) | 1.06 |

Section 13. In Vitro Inhibitory Activity Evaluation.

FBPase activity was measured spectrophotometrically by employing the coupling enzymes PGI (Roche, 10128139001, Switzerland) and glucose-6-phosphate dehydrogenase (G6PDH) (Sigma, 9001-40-5).14 The reduction of NADP^+^ (Macklin, 53-59-8, China) to NADPH was monitored directly at 340 nm. Specifically, 0.11 mg/mL of recombinant FBPase was incubated with serial dilutions of each compound for 12−40 min. At each time point, the NADP^+^ linked enzymatic assay was applied as mentioned above. Relative enzyme activities for various inhibitor concentrations over time were fit to an exponential equation to generate *k*_obs_ values for each concentration tested. Using data from three independent experiments, the resulting *k*_obs_ values were then plotted versus inhibitor concentration, and *K*_i_ and *k*_inact_ values were generated according to the equation.

**Figure S2.** The plot of inhibition kinetic constant (*K*_i_), kinact and the *k*_inact_/*K*_i_ relationship

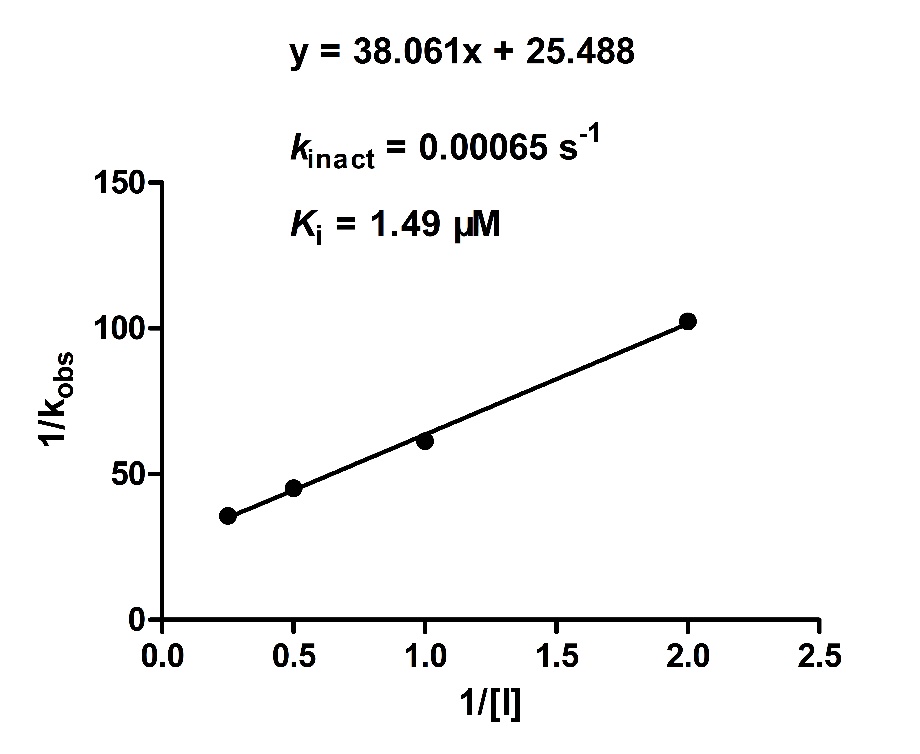

$$\frac{1}{k_{obs}}= \frac{K_{i}}{k_{inact}}\cdot\frac{1}{I}+ \frac{1}{k_{inact}}$$

**Figure S3.** ^1^H NMR of **11t**

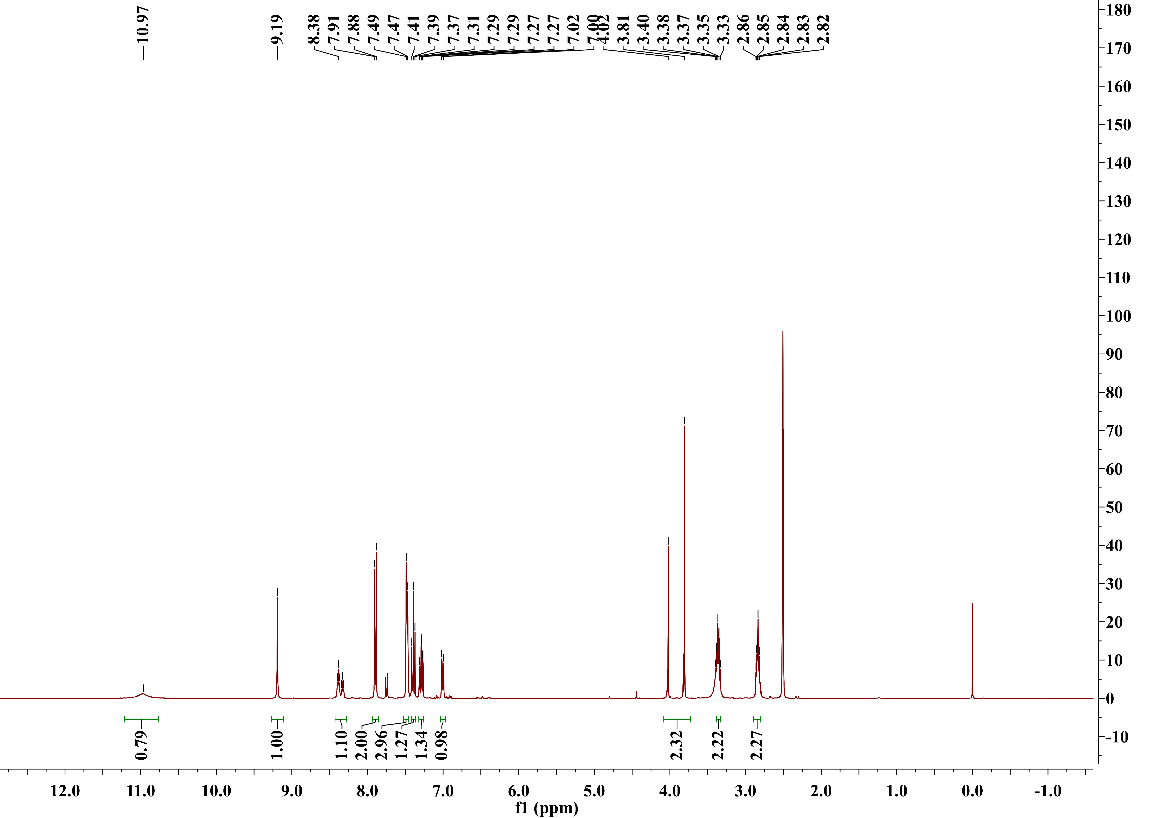

1H NMR (400 MHz, DMSO-d6) δ 10.97 (s, 1H), 9.19 (s, 1H), 8.36 (dt, J = 22.5, 5.5 Hz, 1H), 7.89 (d, J = 8.3 Hz, 2H), 7.48 (d, J = 7.4 Hz, 3H), 7.39 (t, J = 8.2 Hz, 1H), 7.32 – 7.26 (m, 1H), 7.01 (d, J = 8.2 Hz, 1H), 3.91 (d, J = 86.6 Hz, 2H), 3.36 (dd, J = 12.4, 6.6 Hz, 2H), 2.90 – 2.80 (m, 2H). 13C NMR (101 MHz, DMSO-d6) δ 166.54, 149.94, 148.98, 145.81, 140.32, 138.19, 131.05, 129.88, 128.04, 118.23, 115.72, 111.58, 43.06, 35.01, 29.88.

**Figure S4.** ^13^C NMR of **11t**

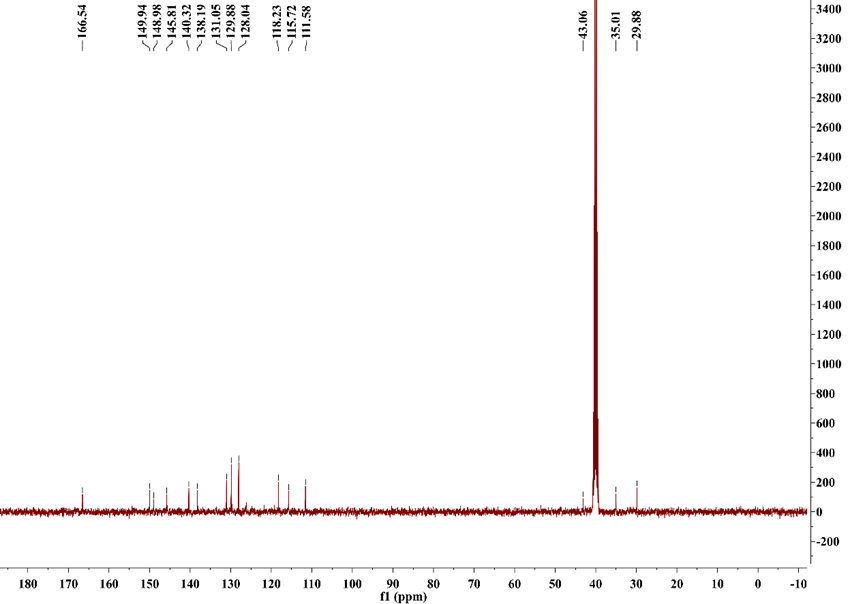

Section 14. Code Availability and Usage

For practical application, we provide a standalone prediction code on GitHub. To execute it, users need the trained model weights along with two essential input files: a DOX.log file (the output from our previously developed DOX or Cov_DOX methods) and a ligand.mol2 file (containing the ligand structure). Note: For covalent inhibitors, the ligand.mol2 file must represent the pre-reactive ligand structure. Following the instructions in the repository's README will run the code and output the final DeepDOX1 prediction.

**Figure S5.** Schematic overview of the DeepDOX1 architecture and its featurization pipeline.

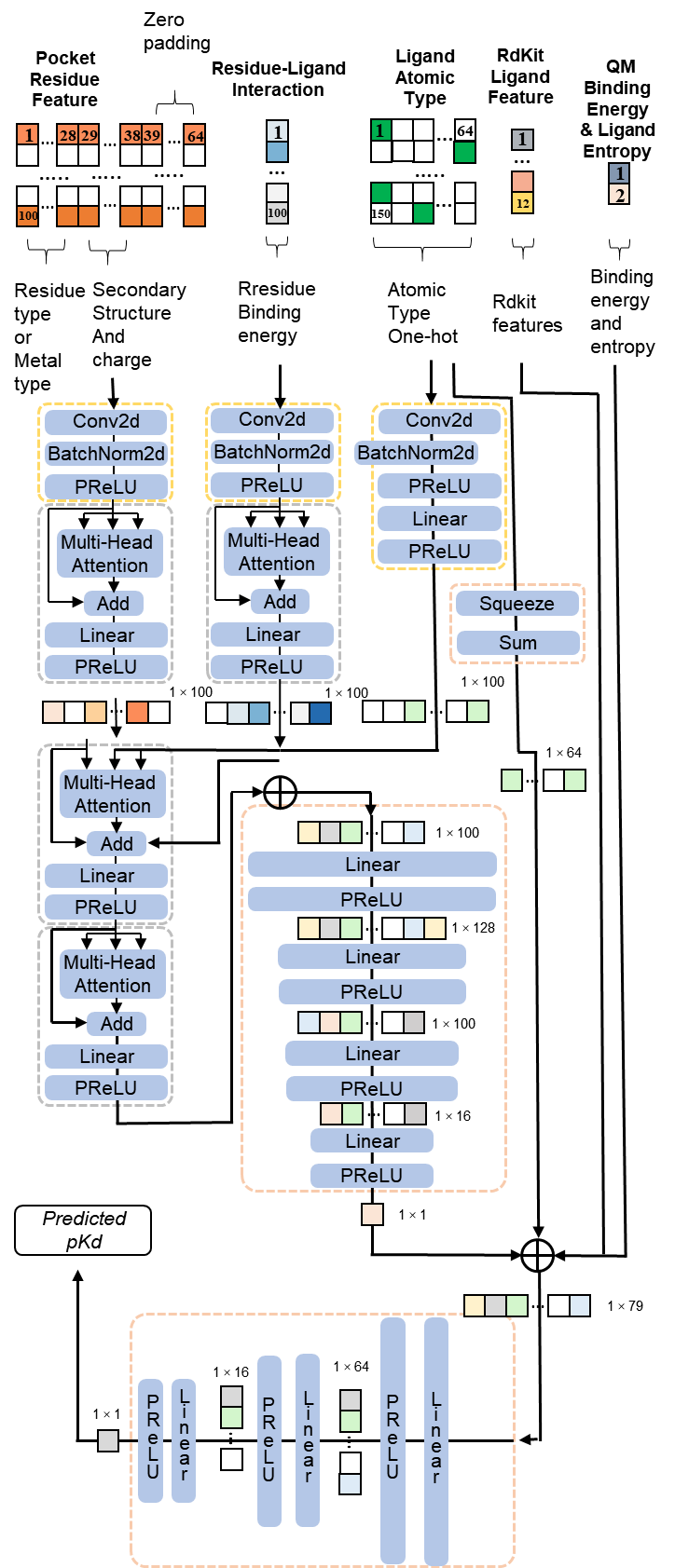

Reference

1.	Su, M. et al. Comparative Assessment of Scoring Functions: The CASF-2016 Update. *Journal of Chemical Information and Modeling* **59**, 895-913 (2018).

2.	Schindler, C.E.M. et al. Large-Scale Assessment of Binding Free Energy Calculations in Active Drug Discovery Projects. *J Chem Inf Model* **60**, 5457-5474 (2020).

3.	Ashwell, M.A. et al. Discovery and optimization of a series of 3-(3-phenyl-3H-imidazo[4,5-b]pyridin-2-yl)pyridin-2-amines: orally bioavailable, selective, and potent ATP-independent Akt inhibitors. *J Med Chem* **55**, 5291-310 (2012).

4.	Leger, P.R. et al. Discovery of Potent, Selective, and Orally Bioavailable Inhibitors of USP7 with In Vivo Antitumor Activity. *J Med Chem* **63**, 5398-5420 (2020).

5.	Kwiatkowski, J. et al. Stepwise Evolution of Fragment Hits against MAPK Interacting Kinases 1 and 2. *J Med Chem* **63**, 621-637 (2020).

6.	Flanagan, M.E. et al. Discovery of CP-690,550: a potent and selective Janus kinase (JAK) inhibitor for the treatment of autoimmune diseases and organ transplant rejection. *J Med Chem* **53**, 8468-84 (2010).

7.	Abel, R., Wang, L., Harder, E.D., Berne, B.J. & Friesner, R.A. Advancing Drug Discovery through Enhanced Free Energy Calculations. *Acc Chem Res* **50**, 1625-1632 (2017).

8.	Jiang, F. et al. Discovery of Benzo[cd]indol-2(1H)-ones and Pyrrolo[4,3,2-de]quinolin-2(1H)-ones as Bromodomain and Extra-Terminal Domain (BET) Inhibitors with Selectivity for the First Bromodomain with Potential High Efficiency against Acute Gouty Arthritis. *J Med Chem* **62**, 11080-11107 (2019).

9.	Rahm, F. et al. Creation of a Novel Class of Potent and Selective MutT Homologue 1 (MTH1) Inhibitors Using Fragment-Based Screening and Structure-Based Drug Design. *J Med Chem* **61**, 2533-2551 (2018).

10.	Ren, H. et al. Design, Synthesis, and Characterization of an Orally Active Dual-Specific ULK1/2 Autophagy Inhibitor that Synergizes with the PARP Inhibitor Olaparib for the Treatment of Triple-Negative Breast Cancer. *J Med Chem* **63**, 14609-14625 (2020).

11.	Feng, X. et al. A Novel Decalin-Based Bicyclic Scaffold for FKBP51-Selective Ligands. *J Med Chem* **63**, 231-240 (2020).

12.	Kump, K.J. et al. Discovery and Characterization of 2,5-Substituted Benzoic Acid Dual Inhibitors of the Anti-apoptotic Mcl-1 and Bfl-1 Proteins. *J Med Chem* **63**, 2489-2510 (2020).

13.	Shin, Y. et al. Discovery of N-(1-Acryloylazetidin-3-yl)-2-(1H-indol-1-yl)acetamides as Covalent Inhibitors of KRAS(G12C). *ACS Med Chem Lett* **10**, 1302-1308 (2019).

14.	Lanman, B.A. et al. Discovery of a Covalent Inhibitor of KRAS(G12C) (AMG 510) for the Treatment of Solid Tumors. *J Med Chem* **63**, 52-65 (2020).

15.	Wei, L. et al. Cov_DOX: A Method for Structure Prediction of Covalent Protein-Ligand Bindings. *J Med Chem* **65**, 5528-5538 (2022).

16.	Ward, R.A. et al. Structure-Guided Design of Highly Selective and Potent Covalent Inhibitors of ERK1/2. *J Med Chem* **58**, 4790-801 (2015).

17.	Wen, W. et al. Structure-Guided Discovery of the Novel Covalent Allosteric Site and Covalent Inhibitors of Fructose-1,6-Bisphosphate Aldolase to Overcome the Azole Resistance of Candidiasis. *J Med Chem* **65**, 2656-2674 (2022).

18.	Jiang, D. et al. MetalProGNet: a structure-based deep graph model for metalloprotein-ligand interaction predictions. *Chem Sci* **14**, 2054-2069 (2023).

19.	Liu, Z. et al. Forging the Basis for Developing Protein-Ligand Interaction Scoring Functions. *Acc Chem Res* **50**, 302-309 (2017).

20.	Cozier, G.E. et al. Design of Novel Mercapto-3-phenylpropanoyl Dipeptides as Dual Angiotensin-Converting Enzyme C-Domain-Selective/Neprilysin Inhibitors. *J Med Chem* **68**, 7720-7736 (2025).

21.	Katz, B.A. et al. Engineering inhibitors highly selective for the S1 sites of Ser190 trypsin-like serine protease drug targets. *Chem Biol* **8**, 1107-21 (2001).

22.	Heffron, T.P. et al. The Rational Design of Selective Benzoxazepin Inhibitors of the alpha-Isoform of Phosphoinositide 3-Kinase Culminating in the Identification of (S)-2-((2-(1-Isopropyl-1H-1,2,4-triazol-5-yl)-5,6-dihydrobenzo[f]imidazo[1,2-d][1,4]oxazepin-9-yl)oxy)propanamide (GDC-0326). *J Med Chem* **59**, 985-1002 (2016).

23.	Kelly, M.J. et al. Discovery of 2-[3,5-dichloro-4-(5-isopropyl-6-oxo-1,6-dihydropyridazin-3-yloxy)phenyl]-3,5-dioxo-2,3,4,5-tetrahydro[1,2,4]triazine-6-carbonitrile (MGL-3196), a Highly Selective Thyroid Hormone Receptor beta agonist in clinical trials for the treatment of dyslipidemia. *J Med Chem* **57**, 3912-23 (2014).

24.	Hurt, D.E., Widom, J. & Clardy, J. Structure of Plasmodium falciparum dihydroorotate dehydrogenase with a bound inhibitor. *Acta Crystallogr D Biol Crystallogr* **62**, 312-23 (2006).

25.	Cui, J.J. et al. Structure based drug design of crizotinib (PF-02341066), a potent and selective dual inhibitor of mesenchymal-epithelial transition factor (c-MET) kinase and anaplastic lymphoma kinase (ALK). *J Med Chem* **54**, 6342-63 (2011).

26.	Qian, Y. et al. Activation and signaling mechanism revealed by GPR119-G(s) complex structures. *Nat Commun* **13**, 7033 (2022).

27.	Pollock, J. et al. Rational Design of Orthogonal Multipolar Interactions with Fluorine in Protein-Ligand Complexes. *J Med Chem* **58**, 7465-74 (2015).

28.	Ali, S., Tian, X., Meccia, S.A. & Zhou, J. Highlights on U.S. FDA-approved halogen-containing drugs in 2024. *Eur J Med Chem* **287**, 117380 (2025).

29.	Hanan, E.J. et al. Discovery of GDC-0077 (Inavolisib), a Highly Selective Inhibitor and Degrader of Mutant PI3Kalpha. *J Med Chem* **65**, 16589-16621 (2022).

30.	Martin, M.P. et al. A novel mechanism by which small molecule inhibitors induce the DFG flip in Aurora A. *ACS Chem Biol* **7**, 698-706 (2012).

31.	Gillis, E.P., Eastman, K.J., Hill, M.D., Donnelly, D.J. & Meanwell, N.A. Applications of Fluorine in Medicinal Chemistry. *J Med Chem* **58**, 8315-59 (2015).

32.	Bolli, M.H. et al. The Discovery of N-[5-(4-Bromophenyl)-6-[2-[(5-bromo-2-pyrimidinyl)oxy]ethoxy]-4-pyrimidinyl]-N′-propylsulfamide (Macitentan), an Orally Active, Potent Dual Endothelin Receptor Antagonist. *Journal of Medicinal Chemistry* **55**, 7849-7861 (2012).
